# Structural Optimization of siRNA Conjugates for Albumin Binding Achieves Effective MCL1-Targeted Cancer Therapy

**DOI:** 10.1101/2023.02.14.528574

**Authors:** Ella N. Hoogenboezem, Shrusti S. Patel, Ashley B. Cavnar, Justin H. Lo, Lauren M. Babb, Nora Francini, Prarthana Patil, Juan M. Colazo, Danielle L. Michell, Violeta M. Sanchez, Joshua T. McCune, Jinqi Ma, Carlisle R. DeJulius, Linus H. Lee, Jonah C. Rosch, Ryan M. Allen, Larry D. Stokes, Jordan L. Hill, Kasey C. Vickers, Rebecca S. Cook, Craig L. Duvall

## Abstract

The high potential for therapeutic application of siRNAs to silence traditionally undruggable oncogenic drivers remains largely untapped due to the challenges of tumor cell delivery. Here, siRNAs were optimized for *in situ* binding to albumin through C18 lipid modifications to improve pharmacokinetics and tumor delivery. Systematic variation of siRNA conjugates revealed a lead structure with divalent C_18_ lipids each linked through three repeats of hexaethylene glycol connected by phosphorothioate bonds. Importantly, we discovered that locating the branch site of the divalent lipid structure proximally (adjacent to the RNA) rather than at a more distal site (after the linker segment) promotes association with albumin, while minimizing self-assembly and lipoprotein association. Comparison to higher albumin affinity (diacid) lipid variants and siRNA directly conjugated to albumin underscored the importance of conjugate hydrophobicity and reversibility of albumin binding for siRNA delivery and bioactivity in tumors. The lead conjugate increased tumor siRNA accumulation 12-fold in orthotopic mouse models of triple negative breast cancer over the parent siRNA. When applied for silencing of the anti-apoptotic oncogene MCL-1, this structure achieved approximately 80% MCL1 silencing in orthotopic breast tumors. Furthermore, application of the lead conjugate structure to target MCL1 yielded better survival outcomes in three independent, orthotopic, triple negative breast cancer models than an MCL1 small molecule inhibitor. These studies provide new structure-function insights on optimally leveraging siRNA-lipid conjugate structures that associate *in situ* with plasma albumin for molecular-targeted cancer therapy.

## Introduction

Short interfering RNA (siRNA) therapies are powerful tools to silence disease-driving genes that are classically ‘undruggable.’ However, in their unmodified form, siRNAs undergo rapid renal clearance and nuclease degradation, resulting in poor *in vivo* target tissue bioavailability. Initial clinical success of siRNA was achieved by the FDA-approved, first-in-class medicine patisiran, comprised of siRNA encapsulated in a lipid nanoparticle, preventing siRNA degradation and mediating hepatic delivery for treatment of hereditary transthyretin-mediated amyloidosis^1^. However, nanoparticle formulations are relatively complex to synthesize and are associated with injection reactions^2, 3^, necessitating premedication with corticosteroids and antihistamine, as in the case of patisiran^4^. The challenges of lipid nanoparticle carriers for siRNA delivery might be avoided with a carrier-free delivery system for chemically modified siRNAs. This notion has been explored successfully, yielding four FDA-approved drugs employing trivalent N-acetylgalactosamine (GalNAc)-based targeting of siRNAs to hepatocytes expressing the asiaglycoprotein receptor (givosiran, lumasiran, inclisiran, vutrisiran). However, clinical success of siRNAs against extrahepatic targets remains elusive.

The molecular precision provided by siRNA makes it an attractive class of drugs for many diseases including cancers, but oncological siRNA applications have not shown success in clinical trials. This may be due, at least in part, to reliance on the idea that the leaky, aberrant tumor vasculature and insufficient lymphatic drainage of tumors, collectively termed the enhanced permeability and retention (EPR) effect, would enable siRNA nanoparticle delivery to tumors, while avoiding healthy tissues. However, nanoparticles designed to leverage the EPR effect have yielded inconsistent results, particularly upon translation into clinical trials^5–8^. Importantly, siRNA delivered by bulky nanocarriers may have limited efficiency in reaching and permeating tumors that are poorly perfused or do not have large-scale lympho-vascular disruptions. In contrast, smaller siRNA conjugates, with structurally-optimized end modifications that improve systemic and cell-level pharmacokinetics, may achieve more homogeneous delivery into tumors^9^. These end modifications can be used to promote preferential *in situ* association with various plasma components as a way to reduce kidney clearance and influence siRNA pharmacokinetics and biodistribution. Albumin is a promising target for use as an endogenous carrier because it is abundant in plasma, has a long half-life, and naturally binds to fatty acids (FAs)^9^, which can be readily attached to siRNA structures.

Albumin also has several traits that motivate its use as an endogenous carrier specifically for tumor applications. Albumin accumulates in the tumor insterstitium in numerous solid cancer models^10–14^. Albumin exhibits >4-fold higher tumor tissue penetration over nanocarriers^15^, and is actively taken up by rapidly dividing tumor cells to meet their high metabolic demand^16–18^. The utility of albumin as a cancer therapeutic carrier is supported by the clinical success of the albumin-based nanoparticle Abraxane for paclitaxel delivery^19^. Moreover, albumin is rescued from renal clearance, an obstacle for systemically administered oligonucleotides^9^. Association with albumin may therefore improve bioavailability of siRNA.

Here we constructed a library of siRNA-lipid conjugates in an effort to identify structural features that promote *in situ* albumin association and functional tumor delivery *in vivo*. The library systematically varies lipid valency, branch point positioning (in divalent lipid-siRNA conjugates), length and phosphorothioate content of the linker, degradability of the linker, and the structure of the albumin-binding lipid. The optimized construct was used to deliver therapeutic siRNA sequences to models of triple negative breast cancer (TNBC), an aggressive breast cancer subtype with a disproportionately high mortality rate, and few therapeutic options.

Because TNBCs, a heterogenous group of cancers, lack the traditional molecular targets associated with HER2+ and ER+ breast cancers, molecularly targeted therapeutics are relatively unavailable to patients diagnosed with TNBCs, making chemotherapy the primary treatment course. Approximately 70% of TNBCs treated with neoadjuvant chemotherapy (NAC) harbor residual disease (RD), rapid recurrence and high mortality^20^. Molecular and genomic analyses of TNBCs following NAC revealed *MCL1* among the most commonly altered genes in RD. MCL-1, an anti-apoptotic oncogene of the BCL2 family is a known driver of treatment resistance in many cancers, including TNBC^21^. Efforts at targeting MCL-1 are being actively pursued, with multiple small molecules entering clinical trial^22^. Therefore, we sought to demonstrate therapeutic proof of concept for the lead siRNA-lipid conjugate structure by applying it to target MCL1 in TNBC models in vivo.

## Materials and Methods

### Reagents

2’-O-Me and 2’-F phosphoramidites, universal synthesis columns (MM1-2500-1), and all ancillary RNA synthesis reagents were purchased from Bioautomation. Symmetrical branching CED phosphoramidite was obtained from ChemGenes (CLP-5215). Cyanine 5 (Cy5) phosphoramidite (10-5915), stearyl phosphoramidite (10-1979), biotin TEG phosphoramidite (10-1955), hexaethyleneglycol phosphoramidite (10-1918), TEG cholesterol phosphoramidite (10-1976), 5’-Amino-Modifier 5 (10-1905), and desalting columns (60-5010) were all purchased from Glen Research. PE anti-mouse CD19 antibody (Catalog No. 115508) and PC anti-mouse/human CD45R/B220 antibody (Catalog No. 103212) were acquired from BioLegend. MIK665 was from Selleck Chem. All other reagents were purchased from Sigma-Aldrich unless otherwise specified.

### Conjugate Synthesis, Purification, and Validation

Oligonucleotides were synthesized using 2’-F and 2’-O-Me phosphoramidites with standard protecting groups on a MerMade 12 Oligonucleotide Synthesizer (Bioautomation). siRNA sequences can be found in **Supplementary Table 1**. Amidites were dissolved at 0.1M in anhydrous acetonitrile, and 2’OMe U-CE phosphoramidite used 20% anhydrous dimethylformamide as a cosolvent. Stearyl phosphoramidite was dissolved in 3:1 (v:v) dichloromethane:acetonitrile. Coupling was performed under standard conditions, and strands were grown on controlled pore glass with a universal terminus (1 μmol scale, 1000Å pore).

Strands were cleaved and deprotected using 1:1 methylamine:40% ammonium hydroxide (25°C, 2 h). Lipophilic RNAs were purified by reversed-phase high performance liquid chromatography (HPLC) using a Clarity Oligo-RP column (Phenomenex) under a linear gradient [85% mobile phase A (50 mM triethylammonium acetate in water) to 100% mobile phase B (methanol) or 95% mobile phase A to 100% mobile phase B (acetonitrile)]. Oligonucleotide fractions were dried (Savant SpeedVac SPD 120 Vacuum Concentrator, ThermoFisher), resuspended in nuclease free water, sterile filtered, and lyophilized.

Conjugate molecular weight and purity (**Supplementary Figure 1**) was confirmed using liquid chromatography-mass spectrometry (LC-MS, ThermoFisher LTQ Orbitrap XL Linear Ion Trap Mass Spectrometer). Chromatography was performed using a Waters XBridge Oligonucleotide BEH C18 Column under a linear gradient [85% phase A (16.3 mM triethylamine – 400 mM hexafluoroisopropanol) to 100% phase B (methanol)] at 45°C. Control conjugate, si-PEG_45L2_, molecular weight was validated using MALDI-TOF MS as previously reported^23^ using 50 mg/mL 3-hydroxypicolinic acid in 1:1 (v:v) water:acetonitrile with 5 mg/mL ammonium citrate as a matrix. Synthesis of amine-reactive lipids and subsequent modification of oligonucleotides was adapted from methods reported by Prakash et al^24^ and described in detail in the supplementary methods. Purified oligonucleotide was resuspended in 0.9% sterile NaCl and annealed to its complementary strand by heating to 95°C and cooling at a rate of 1.66°C/min until 25°C. Duplexes directly bound to albumin were synthesized in a two-step, one-pot reaction described in detail in the supplementary methods. Fluorophore-labeled duplex was used to cross-validate A260 concentration measurements using fluorescence.

### Cell Culture

Cells were cultured in Dulbecco’s modified eagle’s medium (DMEM, Gibco), containing 4.5 g/L glucose, 10% fetal bovine serum [FBS (Gibco)], and 50 μg/mL gentamicin. *Mycoplasma* contamination testing was done by MycoAlert Myocplasma Detection Kit (Lonza). MDA-MB-231, HCC1187, and HCC70 were purchased from ATCC. MDA-MB-231 cells stably expressing a bicistronic green fluorescent protein (GFP) and firefly luciferase (Luc) expression vector (MDA-MB-231.Luc) were seeded at 4,000/well in 96 well plates. HCC70 and HCC1187 cells were seeded at 10,000/well in 96-well plates. After 24 h, cells were treated with siRNA (25nM) using Lipofectamine 2000 (ThermoFisher) in OptiMEM according to the manufacturer’s protocol or treated carrier-free with si<(EG_18_L)_2_ (31.25 nM - 250 nM dose range), replacing with complete media at 24 h post-transfection. Luciferase activity was measured at 48 h post-transfection in cells treated for 5 min with 150 μg/mL D-Luciferin potassium salt (ThermoFisher) using an IVIS Lumina III imaging system (Caliper Life Sciences). Cells were then assessed for caspase activity using Caspase 3/7-Glo at 24h intervals after adding siRNA conjugate. Caspase-3/7 activity was measured 96 h after treatment using Caspase 3/7-Glo (Promega) according to the manufacturer’s directions. Total RNA was harvested at 96 h after treatment (RNeasy, Qiagen). *MCL1* and *PPIB* (housekeeping) mRNA expression was measured in 0.5 μg purified RNA using Quantigene assay (ThermoFisher).

### In Vitro Uptake

For assessment of albumin dependence of uptake, MDA-MB-231 cells were plated at 10,000/well in a 96-well plate 24 h prior to treatment with 100 nM of Cy5-labeled si<(EG_18_L)_2_ or si<(EG_18_L_diacid_)_2_, in Opti-MEM alone (Gibco) or premixed with human serum albumin (Sigma) at 100 nM or 1 μM final concentration in Opti-MEM. The treatment duration was 4 h at 37°C. Cells were washed with PBS, trypsinized, and suspended in PBS + 2% FBS. Fluorescent uptake was assessed using a Guava easyCyte HT flow cytometer (Luminex) with data analyzed in FlowJo (FlowJo, LLC). For assessment of temperature dependence of uptake, MDA-MB-231 cells were plated as above and treated with 100 nM of Cy5-labeled si<(EG_18_L)_2_ or si<(EG_18_L_diacid_)_2_ in Opti-MEM (Gibco) for 2 h at 37°C vs. 4°C (n=3 each). Cell processing and flow cytometry analysis were performed as above.

### Serum Stability

siRNA (0.1 nmol) in 60% FBS in PBS was incubated at 37° for 0-48 h, then resolved on a 2% agarose gel in 1X TAE Buffer. Gels were stained with GelRed Nucleic Acid Stain (Biotium) according to the manufacturer’s protocol and imaged with UV transillumination.

### Immunofluorescence

Samples were snap frozen in OCT embedding medium. Cryosections (6 μm) at multiple tissue depths were fixed for 10 min in 4% paraformaldehyde (PFA), then stained with rabbit anti-firefly luciferase antibody (anti-Fluc, Abcam, ab185924; 1:500), and goat anti-rabbit Alexa Fluor^®^ 488 (ab150077, Abcam; 1:500), counterstained with DAPI and imaged on a Nikon Eclipse Ti inverted confocal microscope. Imaging settings were kept constant across different treatment groups.

### Biolayer Interferometry

Binding kinetics were measured by biolayer interferometry (BLI) at 30°C, 1000 rpm using an Octet RED 96 (ForteBio). Duplexes were synthesized with antisense strand 5’ terminal TEG-biotin, diluted to 500 nM in Dulbecco’s PBS containing Ca^2+^ and Mg^2+^ (DPBS^+/+^), and loaded for 600 sec on a Streptavidin Dip and Read Biosensor (ForteBio). Biosensor association to human and mouse albumin in DPBS^+/+^ was measured for 300 sec, followed by measurements of dissociation for 300 sec. The binding values were measured using Octet Data Analysis HT Software. Interstep correction was performed by aligning to the dissociation step, and noise filtering was performed. Global analysis was performed to derive constants simultaneously from all tested analyte concentrations.

### Critical Micelle Concentration (CMC)

A duplex serial dilution was prepared in a 96-well plate from 20 μM to 10 nM in 50 uL of Ca^2+^/Mg^2+^ free DPBS with Nile Red (0.25 μg), and the plate was rotated at 37°C in the dark for 2 h. Fluorescence was measured on a plate fluorimeter (Tecan) at excitation 535 ± 10 nm and emission 612 ± 10 nm. The CMC was defined, as previously described ^25^, as the intersection point on the plot of the two linear regions of the Nile Red fluorescence versus duplex concentration.

### Electrophoretic Mobility Shift Assay

siRNA conjugates (0.1 nmol) were incubated with 5X molar excess of human or mouse albumin 30 min at 37°C. Complexes were resolved on 4%-20% polyacrylamide gels (Mini-Protean TGX). Nucleic acid was visualized with GelRed Nucleic Acid Stain (Biotium) under ultraviolet imaging and protein visualized by Coomassie Blue stain under visible light imaging.

Conjugation efficacy of DBCO-modified siRNA duplex with azide-modified albumin was visualized using the Agilent Protein 230 Assay on the Agilent 2100 Bioanalyzer according to the manufacturer’s instructions.

### Circulation Half-life and Biodistribution

Intravital fluorescence microscopy was performed using previously reported methods^26^ on a Nikon Czsi+ system. Briefly, isoflurane-anesthetized, 6-8 week old male CD-1 mice (Charles River) were immobilized on a heated confocal microscope stage. Mouse ears were depilated and immobilized under a glass coverslip with microscope immersion fluid. Using light microscopy to visualize ear vessels, images were focused to the plane of greatest vessel width, where flowing red blood cells were visible. Confocal laser microscopy was used to acquire one image per second, at which point Cy5-labeled siRNA (1 mg/kg) was delivered *via* tail vein. Fluorescent intensity within a circular region of interest (ROI), drawn in the focused vein, was used to measure fluorescence decay. Values were normalized to maximum initial fluorescence and fit to a one-compartment model in PK Solver. For assessment of circulation half-life at time points >30 min, blood was collected at various time points from 5 min to 24h in EDTA-coated tubes, diluted in sterile saline, and Cy5 fluorescence measured in 96-well plates by fluorimetry (Tecan). Cy5 fluorescence was quantified in whole organs (heart, lung, liver, kidney, and spleen) using an IVIS Lumina Imaging system (Xenogen Corporation) at excitation and emission wavelengths of 620 and 670 nm, respectively, using Living Image software version 4.4.

### Size Exclusion Chromatography (SEC)

Murine plasma was collected approximately 45 min after a 1 mg/kg intravenous injection of Cy5-labeled siRNA conjugate. Plasma was filtered (0.22 μm) and then injected into an AKTA Pure Chromatography System (Cytiva) with three inline Superdex 200 Increase columns (10/300 GL). Fractionation was done at 0.3 mL/min using Tris running buffer (10mM Tris-HCl, 0.15M NaCl, 0.2% NaN3) into 1.5 mL fractions with a F9-C 96-well plate fraction collector (Cytiva). Cy5 fluorescence was measured in fractions (100 μL) in black, clear-bottom, 96-well plates (Greiner-Bio-one REF 675096) on a SynergyMx (Biotek) at a gain of 120, excitation 642/9.0, emission 675/9.0. Fraction albumin-bound conjugate was determined by taking the sum of fluorescence intensity for fractions associated with albumin elution divided by the sum of fluorescence intensity for all fractions collected. Albumin-associated fractions were determined by running known protein standards through the SEC system and examining A280 of eluate from each of the fractions.

### Orthotopic Mammary Tumor Studies

All animal experiments were IACUC approved and performed according to AAALAC guidelines and NIH best practices in AAALAC-accredited facilities at Vanderbilt University. MDA-MB-231, MDA-MB-231.Luc, HCC70, or HCC1187 cells (1 x 10^6^) in 100 μL of 50% Matrigel were injected into the inguinal mammary fat pads of 4–6-week-old female athymic (*nu/nu*) mice (Envigo). Mice were randomized into treatment groups when tumor volume reached 50 mm^3^ as measured by calipers using the formula Tvol = (length X width^2^) / 2. Mice were treated by intravenous (i.v.) delivery at the indicated doses, durations, and frequencies. Conjugates were delivered in 0.9% saline. MIK665 treatments were delivered in 2% vitamin E/d-α-tocopheryl polyethylene glycol 1000 succinate (Sigma-Aldrich) in NaCl 0.9% (wt/vol) and delivered i.v. via tail-vein at 12.5 mg/kg at the indicated schedule. Mice were humanely euthanized at treatment day 4, 8, 28, or 35 as indicated, or when tumors exceeded 1000 mm^3^. Tissues and tumors were collected at necropsy, and frozen or processed for analysis.

### Flow Cytometry

For tumor cell uptake of Cy5-labeled siRNA-lipid conjugates, tumors were minced in HBSS (with Ca^2+^ and Mg^2+^), dissociated at 37°C for 1h in collagenase (0.5 mg/mL, Roche Life Sciences) / DNase (0.19 mg/mL, BioRAD) in DMEM, and filtered through a 70 μm cell strainer. Erythrocytes were removed using ACK lysis buffer (Thermo Fisher Scientific) for 2 min. Dissociated tumor cells in PBS^-/-^ were assessed by flow cytometry (Guava easyCyte) using FlowJo software. Cell populations were isolated using forward and side scatter. Gating was done to select for GFP+ tumor cells, and Cy5 fluorescence intensity was measured in this population.

For B cell analyses, spleens were pressed into PBS^-/-^ and 1mM EDTA, passed through a 70 μM cell strainer, and washed. Excised femorae and tibiae were flushed with 10 mL of 1% fetal calf serum in EDTA (5 mM) + PBS^-/-^ using a 25G needle. The exudate was filtered (70 μM) and washed. Erythrocytes were removed from splenocytes and bone marrow cells (BMCs) using ACK lysis buffer for 3 min. 1 x 10^6 cells/mL were blocked with mouse Fc block and then stained with the fluorophore-conjugated antibodies CD19-PE (1:200) and B200-APC (1:400). Samples were run on a BD FACS Diva. Compensations and fluorescence-minus-one controls were used to generate flow parameters. B cells were defined as high intensity staining of CD19 PE and B200 APC.

### Expression Analysis in Tissues

For analysis of luciferase reporter silencing, tumor fragments (200-300 mg) were lysed for 1 h on ice with agitation in 1X Reporter Cell Lysis Buffer (Promega) and then centrifuged at 14,000 x g for 15 min at 4°C. Protein concentration was then quantitated using BCA Assay (Pierce). Lysates (20 mg per well) were assessed in 96-well plates using 90 μl reconstituted Luciferase Assay Substrate (Promega) according to the manufacturer’s directions. Luminescence was measured using IVIS grid quantitation.

*MCL1* mRNA was measured in tumor and liver tissue using QuantiGene SinglePlex assay (ThermoFisher). Tissues were harvested, stored, and homogenized in RNAlater (Thermo Fisher) at 4°C. Samples were washed twice with water and then digested for 6 h at 55°C in Quantigene Diluted Lysis Mixture (DLM) supplemented with proteinase K (0.25 mg/ml). Tissue lysates were diluted 1:2 for Quantigene assessment with manufacturer-designed probe sets directed against human (for tumor analysis) and mouse (for liver analysis) *MCL1* and *PPIB*. Luminescent signals generated from each specific probe set were measured and quantified on a plate luminometer (Tecan). Each sample was assessed in 5 technical replicates. Values shown are the average *MCL1* (corrected for the loading control, *PPIB*), relative to the average *MCL1* value in tumors from saline-treated mice.

### Blood Chemistry and Complete Blood Count

Whole blood was collected in EDTA-coated tubes or spun down at 2,000 x g for 15 min at 4°C to isolate plasma. Samples were then submitted to the Vanderbilt Translational Pathology Shared Resource for CBC and chemistry analyses.

### Statistical Analyses

Data were analyzed using GraphPad Prism 7 software (Graphpad Software, Inc.) Statistical tests used for each dataset are provided in the corresponding figure captions. For all figures, * p≤0.05 ** p≤0.01 *** p≤0.001 **** p≤0.0001. All plots show mean ± standard deviation.

## Results and Discussion

### Divalent Lipid Modifier Improves Bioavailablity of Chemically Stabilized siRNAs

We synthesized siRNAs stabilized with alternating 2’F and 2’OMe modifications in a “zipper” pattern and terminal phosphorothioate (PS) linkages (**Fig. 1a**, **Supplementary Fig. 1**) to confer endonuclease and exonuclease resistance^27^. Delivery of zipper- and PS-modified siRNA sequences against luciferase (si_Luc_) inhibited luciferase expression in MDA-MB-231 cells with similar silencing activity to commercially available Dicer substrate siRNAs that sparingly incorporate 2’ modifications (**Supplementary Fig. 2**). However, agarose gel electrophoresis of siRNA following incubation in serum revealed that the zipper- and PS-modified siRNA remained intact, while the Dicer substrate siRNA was mostly degraded within a few hours (**Supplementary Fig. 2**). Henceforth, all siRNAs harbor the zipper and PS modifications, except studies that focus specifically on variation of PS content.

**Figure 1.**
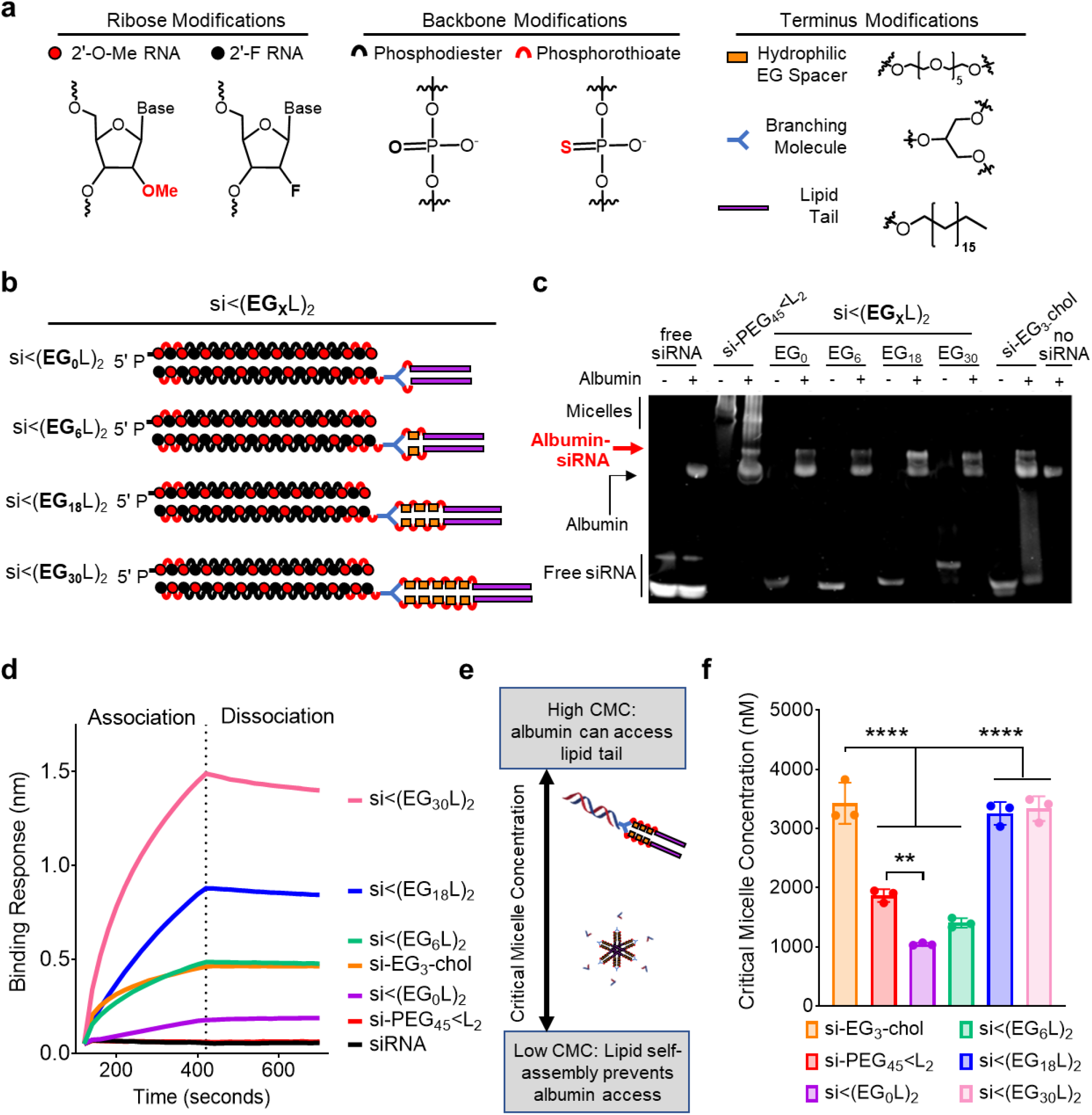
Increased linker EG repeats increases si<(EG_x_L)_2_ albumin binding by reducing self-association. **a-b,** Schematic structural representations of the siRNA and lipid components (**a**) and linker EG repeats (**b**) comprising the novel siRNA-lipid conjugates si<(EG_x_L)_2_. **c,** Representative image of EMSA analysis, used to assess si<(EG_x_L)_2_ interaction with human albumin. **d,** Representative *in vitro* binding responses of siRNA conjugates to human serum albumin (400 nM) measured by biolayer interferometry. **e,** Model illustration of how micellization of siRNA-lipid conjugates, dictated by critical micelle concentration (CMC), influences availability of the albumin-binding lipids. **f,** Critical micelle concentration of siRNA conjugates observed after 2 hours of incubation with Nile Red at 37°C. Each bar represents the average CMC (± S.D.), and each point represents the value measured for an individual experimental replicate, N = 3. ** P < 0.01; **** P < 0.0001

The structure of hydrophobic modifications on siRNA can be tuned to direct siRNA binding to different serum components, such as lipoproteins and albumin, after intravenous (i.v.) administration, which can consequently modify pharmacokinetics and biodistribution^28, 29^. Although there is evidence for active uptake of both albumin^10–14^ and lipoproteins^30–32^ by tumor cells, we sought to preferentially bind to albumin because of its smaller size, lower relative liver tropism, and its capacity for kidney-mediated recycling back into circulation, rather than excretion into urine^29^. The *in vivo* half-life of human albumin is an exceptional 19 days, making it an ideal candidate for increasing the bioavailability of candidate therapeutics^9^.

A library of siRNA-lipid conjugates was generated using solid phase synthesis to maximize yield, purity, and reproducibility, while enabling controlled variation of the linker structure. Reports suggest that valency of lipid end-modified siRNA conjugates affects bioavailability and pharmacodynamics^33–35^. Thus, we compared pharmacokinetics of i.v.-delivered siRNAs conjugated to monovalent (L_1_) or divalent (L_2_) lipid tails comprising 18-carbon stearyls; stearyl was chosen due to its higher affinity to albumin than shorter lipids^24, 36, 37^. Absolute circulation half-life (t_1/2_) of Cy5-labeled siRNA-L_2_, measured in real time using vital fluorescence vascular imaging, was 46 ± 5.9 min; the divalent structure showed higher t_1/2_ than siRNA-L_1_ (28 ± 4.2 min) despite no difference in percent association with albumin in plasma harvested from mice approximately 1h after i.v. injection (**Supplementary Fig. 3**). Whole organ fluorescence was used to assess siRNA-L_1_ and -L_2_ biodistribution approximately 45 min following i.v. delivery, showing decreased renal loss of siRNA-L_2_ as compared to -L_1_. While biodistribution to other organs was similar, siRNA-L_2_ was modestly increased in lungs, a highly vascularized tissue, perhaps reflecting circulating conjugate at this timepoint (**Supplementary Fig. 3**). Based on these findings, the zipper- and PS-stabilized siRNA-L_2_ construct was used as the basis for additional structure-function exploration.

### Hydrophilic Linker Length Increases Albumin Affinity while Reducing Self-Assembly of Lipid-siRNA Conjugates

Variants of siRNA-L_2_ were synthesized with 0 to 30 ethylene glycol (EG) repeats [siRNA<(EG_x_L)_2_] within the hydrophilic linker bridging the lipid and siRNA moieties; structures were created in increments of 6 using a hexaethylene phosphoramidite (**Fig. 1a-b**). For comparative reference, two previously reported siRNA-lipid conjugates were included: cholesterol-TEG-siRNA^38–40^ (si-EG_3_-chol) and si-PEG_45_<L_2_^23^. Electrophoretic mobility shift assay (EMSA) assessing albumin association revealed the shifted mobility of si<(EG_0_L)_2_, si<(EG_6_L)_2_, si<(EG_18_L)_2_, and si<(EG_30_L)_2_ following incubation with human albumin, while mobility of free siRNA was unaltered by albumin (**Fig. 1c**), confirming that all siRNA<(EG_x_L)_2_ conjugates have some level of association with albumin. Similar results were observed using mouse serum albumin (**Supplementary Fig. 4**). The si-EG_3_-chol benchmark compound also displayed albumin-dependent mobility shifts (**Fig. 1c**). However, mobility shifting of si-PEG_45_<L_2_ was seen in both the presence and absence of albumin, suggesting that, in addition to partial binding to albumin, si-PEG_45_<L_2_ self-associates into micelles.

Biolayer interferometry (BLI) was used to measure albumin association and dissociation kinetics of the si<(EG_x_L)_2_ variants (**Fig. 1d**, **Supplementary Fig. 4**). As expected, free siRNA did not exhibit albumin binding. Increasing EG linker length progressively increased the affinity of si<(EG_x_L)_2_ for albumin, with both si<(EG_18_L)_2_ (KD = 30 ± 0.3 nM) and si<(EG_30_L)_2_ (KD = 9.49 ± 0.1 nM) surpassing what was seen with si-EG_3_-chol. These findings are consistent with previous reports showing the influence of linker architecture on the albumin affinity of semaglutide, a therapeutic GLP-1 agonist^41^, and highlight the impact of the EG repeats within the linker region.

Amphiphilic lipid-modified nucleic acids may self-assemble into micellar structures^42–44^, potentially interfering with albumin association if the albumin-binding lipid tails sequester in the micellar core (**Fig. 1e**). To establish the impact of linker length on si<(EG_x_L)_2_ self-assembly, the critical micelle concentration (CMC) was determined (**Figure 1f, Supplementary Fig. 5**). As predicted by cholesterol’s relatively bulky structure that interferes with tight packing, a relatively high CMC was measured for si-EG_3_-chol (3430 ± 350 nM)^45^, while si-PEG_45_<L_2_ exhibited a lower CMC (1860 ± 60 nM), indicating that si-PEG_45_<L_2_ is more likely to self-associate. Interestingly, we observed a correlation between EG repeats and CMC, with si<(EG_0_L)_2_ having the lowest CMC (1040 ± 23 nM), and si<(EG_18_L)_2_ and si<(EG_30_L)_2_ harboring the highest CMCs (3260 ± 190 nM and 3330 ± 210 nM). These results demonstrate that increasing the EG spacer length diminishes the propensity for siRNA-lipid micellization, an important consideration when employing long FA chains for albumin binding *in situ*.

### Hydrophilic Linker Length Determines Pharmacokinetics and *In Vivo* Plasma Disposition of Lipid-siRNA Conjugates

Cy5-labeled si<(EG_x_L)_2_ variants were administered i.v. to assess linker effects on pharmacokinetics. Intravital fluorescence microscopy of mouse ear vasculature demonstrated the rapid and complete diminution of circulating free siRNA within 30 min, while si-EG_3_-chol and si-PEG_45_<L_2_ retained some observable circulating siRNA at 30 min, as did each of the si<(EG_x_L)_2_ (**Fig. 2a**). Real-time measurements of circulating Cy5 fluorescence over time enabled calculation of absolute half-life (t_1/2_), demonstrating the progressively increasing t_1/2_ with increasing EG repeats, with the t_1/2_ of si<(EG_18_L)_2_ nearly 5-fold higher than si<(EG_0_L)_2_. Interestingly, the t_1/2 for_ si<(EG_30_L)_2_ was less than what was seen for si<(EG_18_L)_2_ **(Fig. 2b, Supplementary Table 2**), suggesting a limit to the positive impact of increasing linker length on circulation half-life.

**Figure 2.**
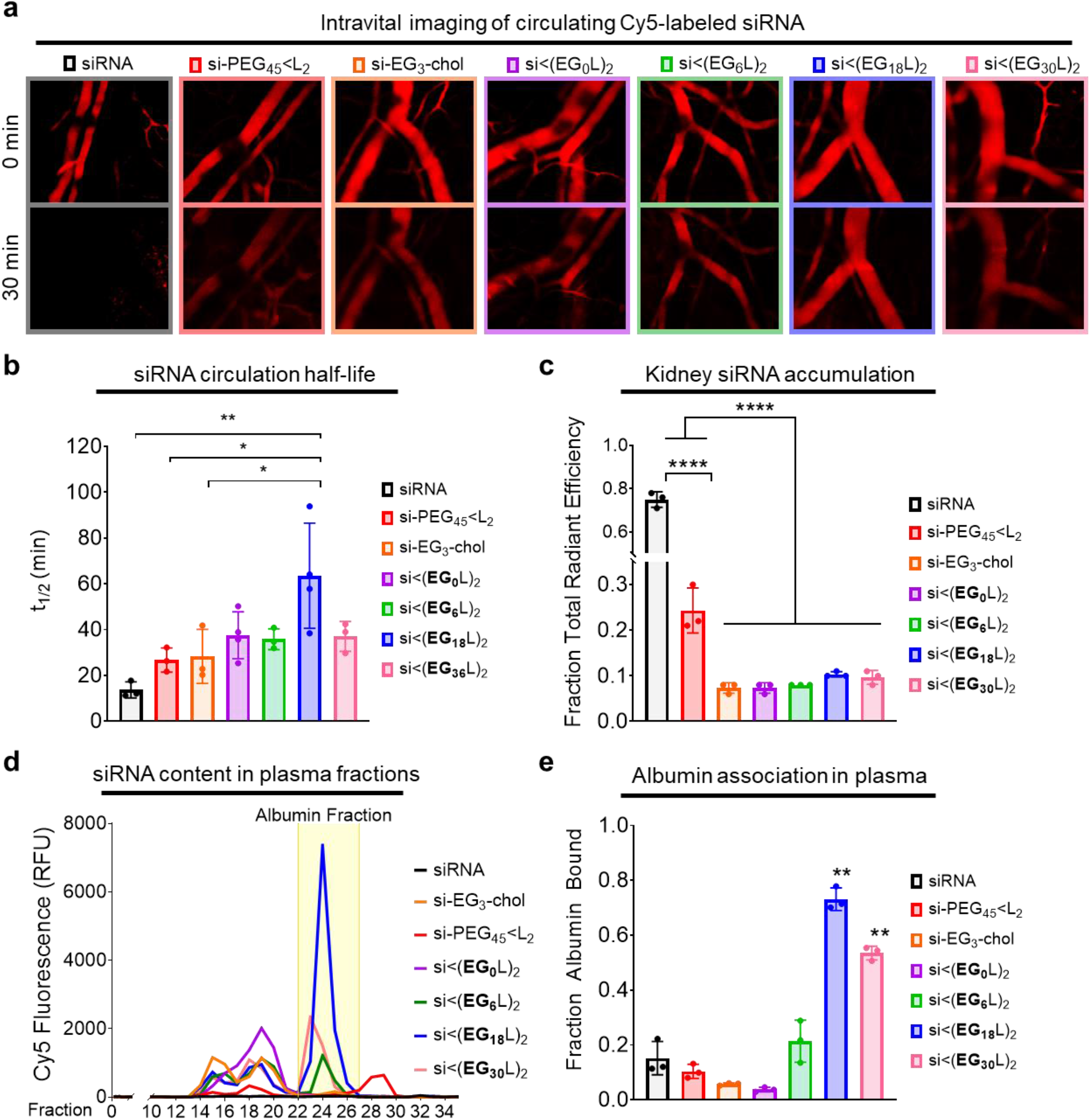
The si<(EG_18_L)_2_ structure achieves the longest circulation half-life and highest albumin association in plasma. Cy5-labeled conjugate or parental siRNA structures were delivered i.v. to mice at 1 mg/kg. **a,** Representative images of intravascular Cy5 fluorescence measured in mouse ear vasculature at 0 min and 30 min after treatment are shown. **b,** Intravascular Cy5 fluorescence recorded over approximately 1h following delivery was used to calculate absolute siRNA circulation half-life (t_1/2_). **c,** Cy5 fluorescence in kidneys, lungs, liver, spleen, and heart was measured approximately 1h after treatment. Fluorescence values in kidneys relative to the total fluorescent values of all organs combined is shown. **d-e,** Plasma samples collected from mice approximately 1h after treatment were fractionated and quantitated for Cy5 fluorescence by size exclusion chromatography. A representative trace is shown of fluorescence signal within each plasma fraction for each siRNA structure; the albumin-containing plasma fractions are highlighted (**d**). Cy5 fluorescence within the albumin-containing fractions was quantitated relative to total plasma fluorescence (**e**). For panels b, c, and e, each bar represents the average value (± S.D.), each point represents values measured for each individual mouse, N = 3-4; * P < 0.05; ** P < 0.01; **** P < 0.0001

Approximately 1h after injection, siRNA biodistribution was measured as Cy5 fluorescence in whole organs, revealing abundant renal accumulation of free siRNA, but substantially reduced renal accumulation of all si<(EG_x_L_2_) variants (**Fig. 2c, Supplementary Fig. 5**). Interestingly, the previously described si-PEG_45_<L_2_ exhibited more renal accumulation than the new si<(EG_x_L_2_) variants. This observation may be due to hydrolytically degradable ester bonds within the DSPE-PEG_2000_ used for si-PEG_45_<L_2_ synthesis. Ester hydrolysis might release siRNA from albumin prematurely in circulation, resulting in renal clearance and shorter circulation time relative to the more stable si<(EG_x_L_2_) variants. Albumin itself harbors intrinsic esterase activity, making the hydrolytic stability of albumin-interacting drugs particularly important^46^. However, si<(EG_x_L_2_) variants likely remain albumin-bound, thus evading renal clearance. This hypothesis was tested by size exclusion chromatography of plasma collected from mice after i.v. delivery of the siRNA conjugates (**Supplementary Fig. 6)**, revealing only ~10% of si-PEG_45_<L_2_ associating with the albumin-containing fraction of plasma, similar to what was seen for free siRNA (**Fig. 2d-e**). In contrast, increasing the EG content of the hydrophilic linker within the si<(EG_x_L_2_) variants increased their association with the albumin-containing plasma fraction, with ~75% of si<(EG_18_L)_2_ within the albumin-containing peak (**Fig. 2e**), consistent with the high albumin affinity of si<(EG_18_L)_2_ measured by biolayer interferometry (**Fig. 1d**) and in support of the idea that albumin-binding in vivo supports circulation half-life (**Fig. 2b**). Interestingly, the percentage of si<(EG_30_L)_2_ associated with the albumin-containing fractions was decreased compared to si<(EG_18_L)_2_. While si<(EG_30_L)_2_ shows some albumin association in vivo, it also partially elutes with earlier fractions associated with large lipoproteins. Therefore, while both si<(EG_18_L)_2_ and si<(EG_30_L)_2_ demonstrate high affinity for purified albumin in vitro, si<(EG_18_L)_2_ shows a greater selectivity for albumin association in the complex, in vivo environment, suggesting that association with circulating albumin in vivo (**Fig. 2d-e**) is a better predictor of circulation half-life (**Fig. 2b**) versus affinity to purified albumin in vitro (**Fig. 1d**).

### Albumin-Binding Increases Carrier Free Tumor Accumulation of siRNA-Lipid Conjugates and Tumor Gene Silencing

To assess tumor accumulation of si<(EG_x_L)_2_ variants, female athymic mice harboring orthotopic Luc-expressing MDA-MB-231 TNBC tumors were treated (2.5 mg/kg, i.v.) on days 1, 3, and 5 with a si_Luc_ sequence synthesized into the different si<(EG_x_L)_2_ variants (**Fig. 3a**). Mice were treated (1 mg/kg) i.v. on day 6 with inactive, Cy5-labeled si<(EG_x_L)_2_ for measuring whole tissue fluorescence in tumors and kidneys approximately 18 h after treatment. These studies revealed abundant kidney fluorescence in mice treated with free siRNA or si-PEG_45_<L_2_, and little fluorescence in tumors (**Fig. 3b**). However, progressively increased tumor fluorescence with conversely decreasing kidney fluorescence was observed in mice treated with si<(EG_0_L)_2_, si<(EG_6_L)_2_, and si<(EG_18_L)_2_, resulting in a tumor:kidney fluorescence ratio of 0.5, 0.9, and 1.0, respectively. These data suggest that improved association with circulating albumin *in vivo* may improve both siRNA circulation time and siRNA tumor accumulation, and that these outcomes can be optimized by tuning of the EG linker repeat length. These observations are in line with findings that increasing circulation time enables higher tumor exposure and interaction with target cells^47, 48^. Consistently, tumor to kidney fluorescence was lower in mice treated with si<(EG_30_L)_2_ compared to si<(EG_18_L)_2_, consistent with its diminished association with the albumin fraction *in vivo*.

**Figure 3.**
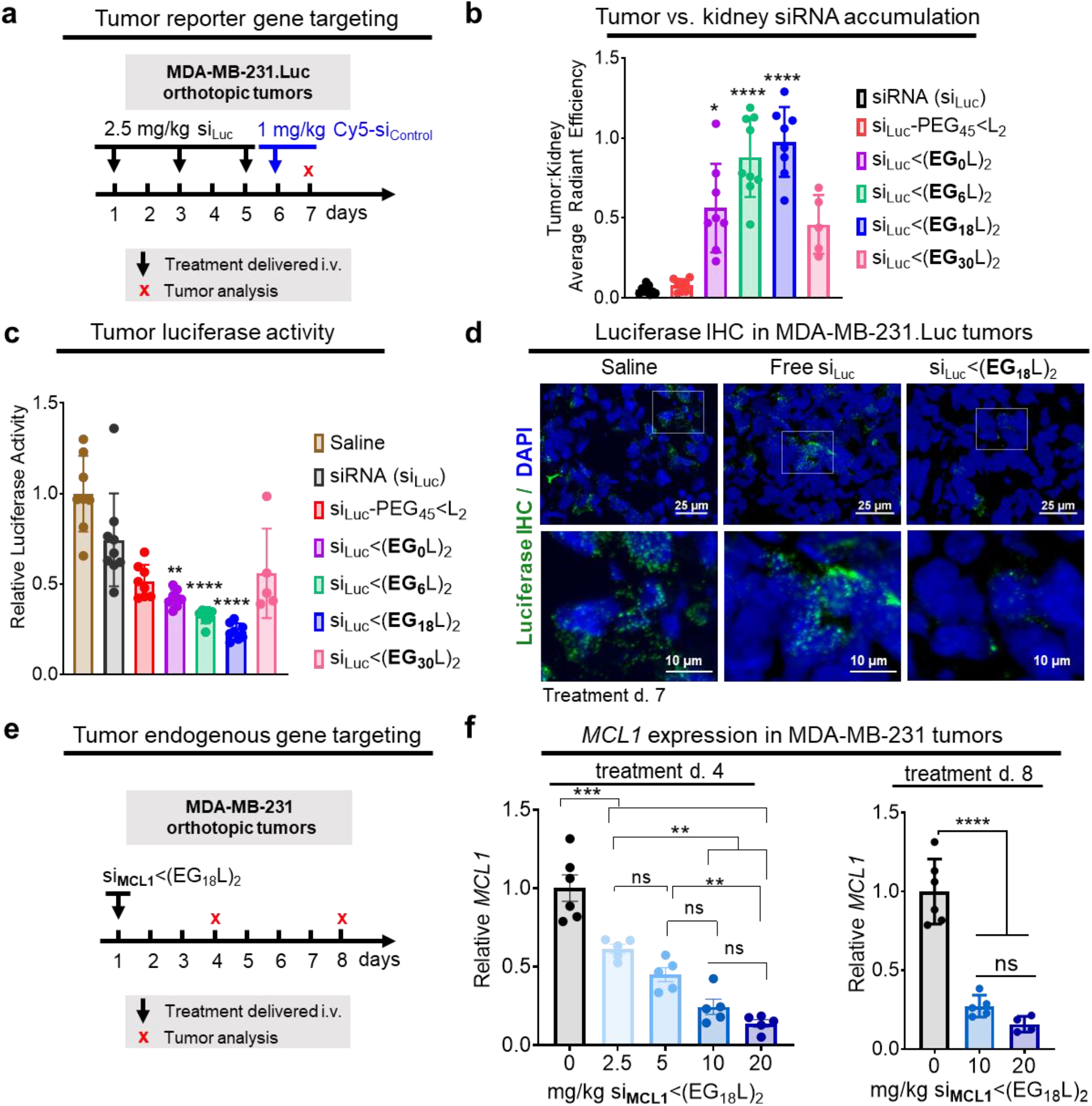
Robust tumor accumulation and tumor gene silencing using si<(EG_18_L)_2_. **a-d,** Mice bearing intra-mammary MDA-MB-231.Luc tumors were treated with siRNA-lipid conjugates, or free siRNA, using siRNA sequences against luciferase (si_Luc_). **a,** Schematic showing treatment schedule for i.v. delivery of siRNA-lipid conjugates, or free siRNA, to tumor-bearing mice. **b,** Cy5 fluorescence tumor:kidney ratio for each mouse as measured on day 7 (18h after injection with fluorescent siRNA). **c,** Luciferase activity was measured in tumor lysates collected on treatment day 7. Bars show the average (± S.D.) tumor luciferase per group, and points are the values for each tumor (assessed in triplicate). All values shown are relative to the average luciferase activity of saline-treated controls, which was set to a value of 1. **d,** Tumors were assessed by IHC analysis for firefly luciferase (green). Nuclei were counterstained with DAPI (blue). Representative images are shown, n = 5-8. **e-f,** Mice bearing intra-mammary MDA-MB-231 tumors were treated one time with si<(EG_18_L)_2_, using siRNA sequences against *MCL1* (si_MCM1_) at increasing doses (2.5 mg/kg - 20 mg/kg). Tumors were assessed on treatment day 4 and day 8. **e,** Schematic showing i.v. delivery of siRNA-lipid conjugates, or free siRNA, to tumor-bearing mice. **f,** Human MCL1 mRNA levels measured in whole tumor RNA collected on treatment day 4 (left panel) and day 8 (right panel). Bars show the average (± S.D.), and points are the values for an individual tumor (assessed in triplicate). All values shown are relative to values measured in non-targeting controls, which was set to a value of 1. * P < 0.05; ** P < 0.01; *** P < 0.001; **** P < 0.0001

Whole tumor luciferase activity was measured as a readout for siRNA silencing activity in tumors collected 18 h after final treatment (day 8), revealing nearly 80% luciferase activity reduction in the tumors of si<(EG_18_L)_2_ treated mice (**Fig. 3c**), more than twice the knockdown seen with the previously reported si-PEG_45_<L_2_. These results were confirmed by immunohistochemical (IHC) staining for luciferase (**Fig. 3d, Supplementary Fig. 7)**. Other reported examples of carrier-free siRNA delivery^47, 49–51^, including cholesterol-siRNA^52^ (10 mg/kg) and receptor-targeted Centyrin-siRNA^47^ (3 x 10 mg/kg), produced target gene knockdown in tumors, but none as potent as si<(EG_18_L)_2_.

To assess knockdown of the endogenous oncogenic driver MCL-1 in tumors, si<(EG_18_L)_2_ was synthesized using *MCL1* siRNA sequences (si_MCM1_) and delivered i.v. as a single bolus to MDA-MB-231 tumor bearing mice (**Fig. 3e**), revealing dose-dependent *MCL1* knockdown at day 4 and day 8 (**Fig. 3f**), with as much as 85% *MCL1* knockdown. Although the *MCL1* siRNA sequence used herein was designed to target both human and mouse *MCL1*, we found less than 20% *MCL1* knockdown in livers of mice treated at the highest dose (**Supplementary Fig. 8**). Given that the si<(EG_18_L)_2_ structure shows the most promise based on a combination of or albumin binding, circulation time, tumor accumulation, and tumor gene silencing over other candidates investigated here, our subsequent analyses further interrogate the structural components of si<(EG_18_L)_2_.

### Phosphorothioate linkages in the 5’ sense and linker region contribute to albumin association and PK

Stabilization of the 5’ sense terminus with phosphorothioate (PS) linkages in lieu of phosphodiester (PO) linkages reportedly confers exonuclease resistance, while enabling extrahepatic, carrier-free gene silencing applications^53, 54^. A variant of si<(EG_18_L)_2_ was synthesized, lacking the PS bond at the 5’ sense (Se) terminus (No 5’Se PS). In another variant, PS bonds were also removed from the bonds within the linker (i.e., between the EG units) between the siRNA and the stearyl groups (No 5’Se or Binder PS) (**Extended Data Fig. 1a**). PS bond removal from the linker significantly increased the CMC, suggesting a lower tendency to self-assemble (**Supplementary Fig. 9)**. Although albumin affinity was unaffected by PS bond removal in either variant, intravital fluorescence microscopy of circulating Cy5-labeled siRNA variants revealed that removal of PS bonds from both the 5’ sense terminus and linker region profoundly diminished t_1/2_ and reduced association with plasma albumin by nearly 10-fold (**Extended Data Fig. 1b**). However, removal of the PS from the 5’ sense terminus alone had only minimal impact on plasma albumin association. Whole tissue Cy5 fluorescence approximately 1h post-delivery showed that loss of 5’ sense terminus PS bond alone modestly but significantly diminished liver siRNA accumulation compared to parental si<(EG_18_L)_2_ (**Extended Data Fig. 1c**). Because kidney accumulation was similar for all variants tested, we speculate that the diminished t_1/2_ of variants lacking PS bonds is most likely due to reduced albumin association rather than increased degradation. This conclusion is also supported by prior reports that PS backbones increase oligonucleotide affinity for plasma proteins^55, 56^. Due to poor circulation half-life of the si<(EG_18_L)_2_ variant lacking PS bonds in both the 5’ sense terminus and linker regions, we compared tumor siRNA accumulation and activity of the parental si<(EG_18_L)_2_ construct and the variant lacking only the 5’ sense PS (**Extended Data Fig. 1d**). These studies revealed no significant differences in tumor accumulation and cell uptake, but si<(EG_18_L)_2_ with no 5’ sense PS bonds exhibited significantly diminished tumor gene silencing (**Extended Data Fig. 1e-f)**. Whereas our lead construct demonstrated nearly 70% knockdown of the target gene, the elimination of the 5’ sense PS bonds resulted in only about 20% knockdown. The comparable biodistribution and uptake of both compounds but large disparity in knockdown suggests that these bonds are stable during circulation and extravasation. However, the PO bonds are likely degraded after biodistribution to the tumor, removing the lipid end groups, and reducing siRNA interactions with and passage through cell and endosomal membranes.

### Proximal positioning of divalent lipid branching point, not relative EG content, dictates tendency to associate with circulating albumin over self-assembly

We next assessed the impact of divalent si<(EG_18_L)_2_ conjugate branchpoint position on albumin binding and micellization. It was hypothesized that branchpoint placement between the EG linker repeats and the stearyl groups, rather than between the siRNA and the EG repeats, will constrain the two stearyls and consequently increase self-assembly. We generated two new analogs of si<(EG_18_L)_2_ with a distal branchpoint location, one matched for overall EG content (si-EG_36_<L_2_) and one matched for the number of EG repeats between the siRNA the lipids (si-EG_18_<L_2_) (**Fig. 4a)**. Interestingly, BLI measurement showed that both new branching point variants had lower albumin binding affinity relative to the parent construct si<(EG_18_L)_2_ (**Fig. 4b**). The CMC, measured by Nile Red encapsulation, was significantly reduced for si-EG_36_<L_2_ (1838 ± 117 nM) and si-EG_18_<L_2_ (2293 ± 132 nM) compared to si<(EG_18_L)_2_ (3255 ± 192 nM) (**Fig. 4c**), suggesting that the proximal branch site of si<(EG_18_L)_2_ increases albumin association due, in part, to a decreased tendency for self-assembly.

**Figure 4.**
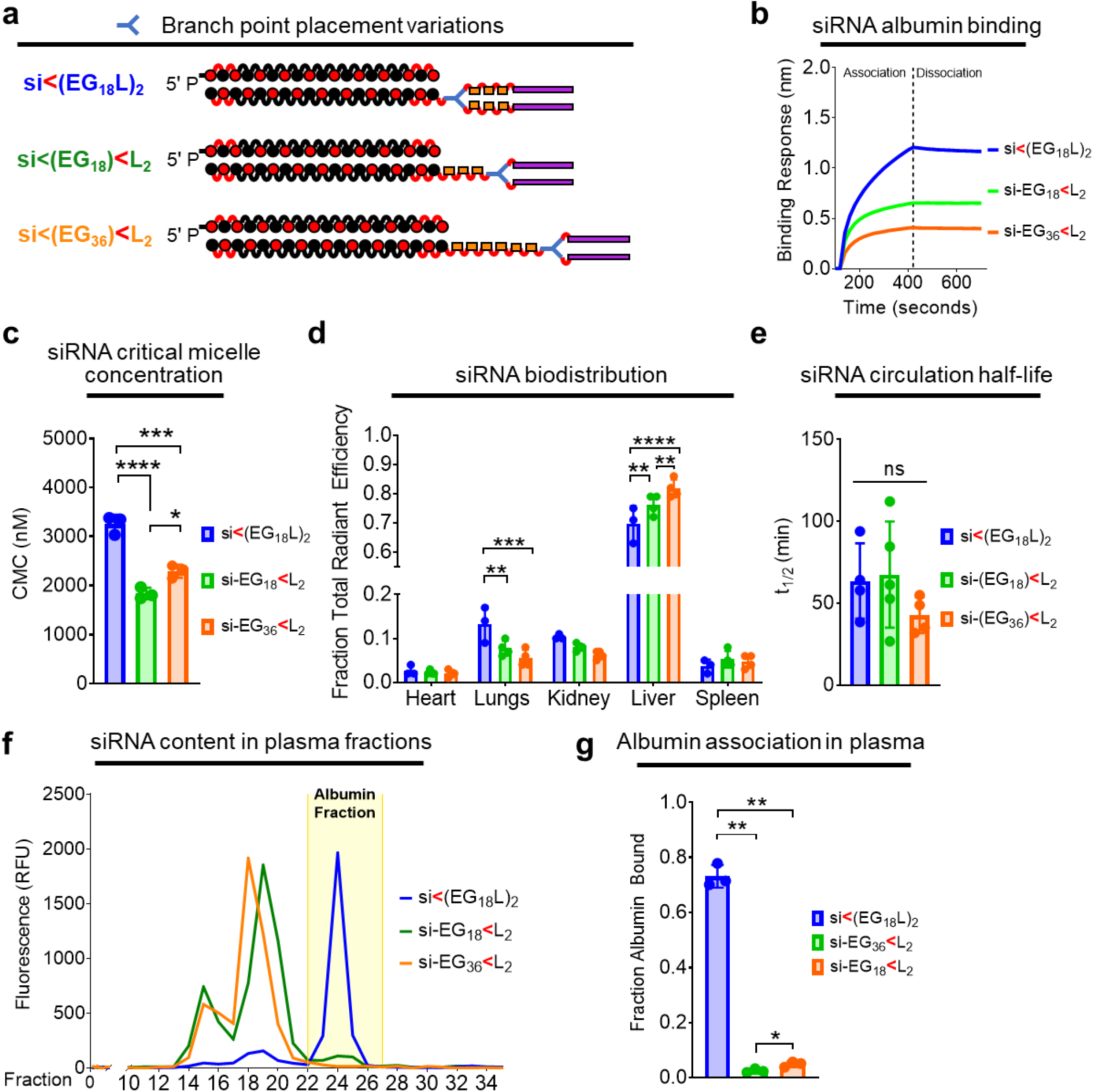
Proximal placement of the divalent lipid branch point immediately adjacent to the siRNA drives albumin binding. **a,** Schematic structural representations of the siRNA-lipid conjugates with varied branchpoint placement and length of the linker. **b,** Representative *in vitro* binding responses of siRNA conjugates to human serum albumin (400 nM) measured by biolayer interferometry. **c,** Critical micelle concentration of siRNA conjugates observed after 2 hours of incubation with Nile Red at 37°C. **d,** Cy5 fluorescence in kidneys, lungs, liver, spleen and heart was measured approximately 1h after i.v. delivery of siRNA-lipid conjugates (1 mg/kg). Fluorescence values in each organ relative to the total fluorescent values of all organs combined is shown. **e,** Intravascular Cy5 fluorescence measured through 1h following i.v. delivery of siRNA-lipid conjugate variants (1 mg/kg) was used to calculate siRNA circulation half-life (t_1/2_). **f,** Plasma samples collected from mice approximately 1h after treatment (1 mg/kg) were fractionated and quantitated for Cy5 fluorescence by size exclusion chromatography. A representative trace is shown of fluorescence signal within each plasma fraction for each siRNA structure; the albumin-containing plasma fractions are highlighted. **g,** Cy5 fluorescence within the albumin-containing fractions was quantitated relative to total plasma fluorescence. For panels C-E and G, each bar represents the average value (± S.D.), and each point represents a value measured for an individual sample. * P < 0.05; ** P < 0.01; *** P < 0.001; **** P < 0.0001

Significantly higher liver accumulation of si-EG_18_<L_2_ and si-EG_36_<L_2_ was seen at 1h post-delivery as compared to si<(EG_18_L)_2_ (**Fig. 4d**) an observation that can be attributed to serum lipoprotein binding, as opposed to albumin binding^28^. Although no significant difference in t_1/2_ was seen among the branching architecture variants (**Fig. 4e**), only about 5% of the delivered si-EG_18_<L_2_ and si-EG_36_<L_2_ conjugates was associated with the plasma albumin fraction *in vivo*, versus ~75% of the parental conjugate (**Fig. 4f-g**). These data show that the novel, proximal branchpoint placement used in the discovery of si-EG_18_<L_2_ substantially diminishes self-assembly and enhances albumin association over commonly used distal branch points that are directly adjacent to the multivalent lipids^23, 33, 35, 36^.

### Hydrophobicity of lipid is more important than binding affinity for conjugate performance

To determine if the nature of the C_18_ lipid affects the performance of si<(EG_18_L)_2_, we synthesized a variant with carboxyls on each terminus [si<(EG_18_L_diacid_)_2_] (**Fig. 5A**). Although the FA carboxyl functionality is usually consumed in conjugation reactions, screening of various peptide structures that led to the development of GLP-1 agonist drug Semaglutide demonstrated that maintaining a free terminal carboxyl increases albumin affinity and circulation half-life^41^. We also generated a variant with a double bond (si<(EG_18_L_unsaturated_)_2_), based on reports of higher affinity between albumin and oleate, an unsaturated variant of stearate; testing of oleate lipids was also motivated for the integration of structural “kinks” caused by FA double bonds that may prevent close packing and thus deter self-assembly^57–60^. First, si<(EG_18_-amine)_2_ was synthesized, to which amine-reactive PFP-modified lipid variants were conjugated (**Supplementary Fig. 11**). Although t_1/2_ and kidney accumulation were similar in si<(EG_18_L_diacid_)_2_ and si<(EG_18_L_unsaturated_)_2_ as compared to si<(EG_18_L)_2_, si<(EG_18_L_unsaturated_)_2_ exhibited significantly diminished association with the albumin-containing plasma fraction *in vivo* compared to its parental (i.e., saturated) and diacid counterparts (~40% bound versus ~75-80% bound) (**Fig. 5b-c**). Strikingly, si<(EG_18_L_diacid_)_2_ showed a substantially increased affinity for human albumin (**Fig. 5c**), with two orders of magnitude lower KD than the parental si<(EG_18_L)_2_ (**Supplementary Fig. 12**). To determine whether the impact of this increased affinity for albumin would extend the circulation time of si<(EG_18_L_diacid_)_2_, we measured circulating Cy5 fluorescence through 24 h following delivery at 5 mg/kg siRNA but found no significant difference between the circulation time of si<(EG_18_L_diacid_)_2_ or si<(EG_18_L)_2_ (**Supplementary Fig. 12**).

**Figure 5.**
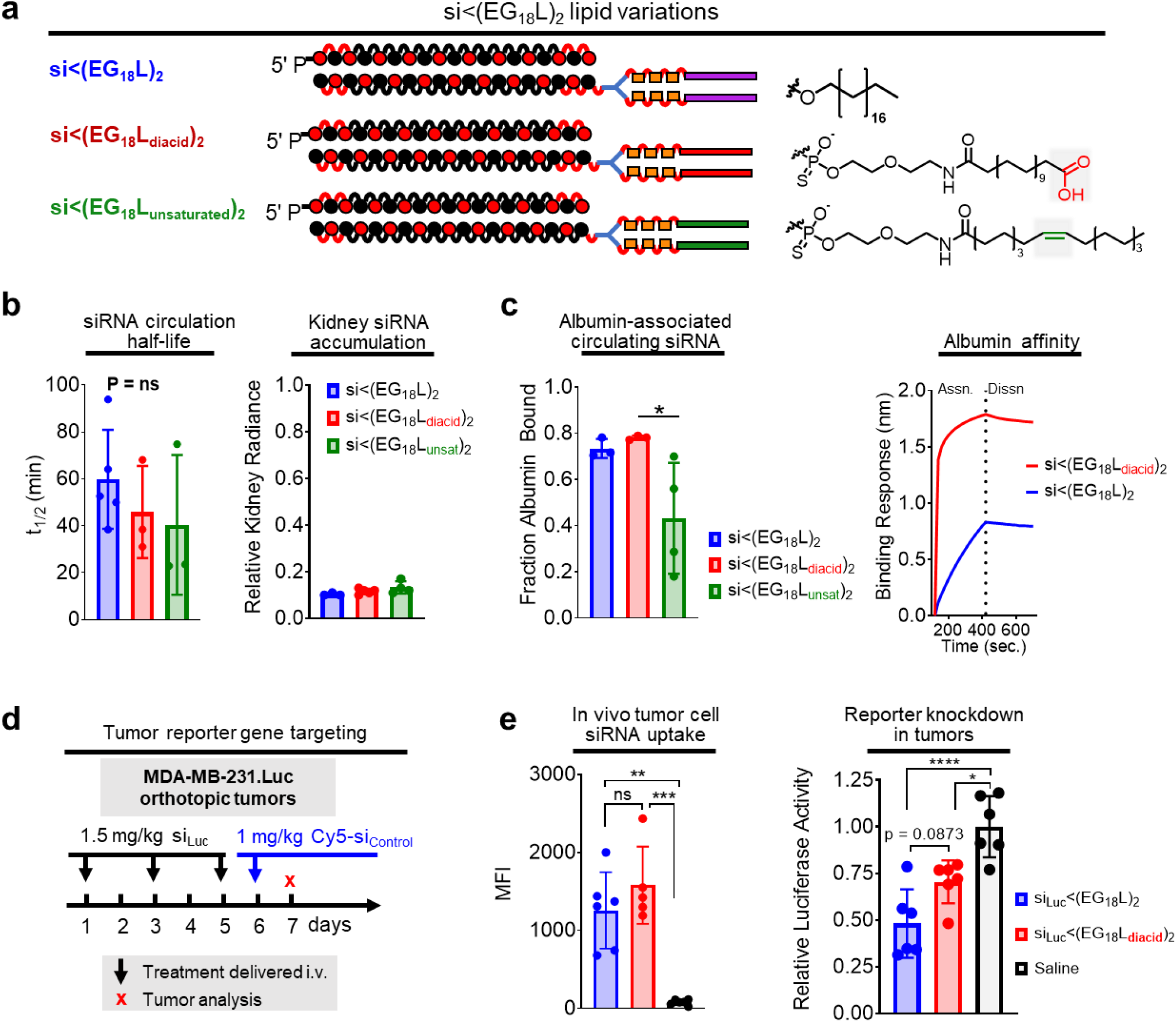
Effects of unsaturated and carboxy-terminated lipid structures. **a,** Schematic structural representations of the siRNA-lipid conjugates with varied lipid chemistries. **b,** Cy5-labeled siRNA-lipid conjugates were delivered (1 mg/kg) i.v. Intravascular Cy5 fluorescence measured over approximately 1h was used to calculate siRNA circulation half-life (left panel). Cy5 fluorescence in kidneys, lungs, liver, spleen, and heart was measured 1h after treatment. Fluorescence values in kidneys relative to the total fluorescence values of all organs combined is shown (right panel). **c,** Size exclusion chromatographic calculation of fraction of Cy5 fluorescence associated with albumin-containing plasma fractions (left panel). Representative *in vitro* binding responses of siRNA conjugates to human serum albumin (400 nM) measured by biolayer interferometry (right panel). **d,** Schematic showing treatment schedule for i.v. delivery of siRNA-lipid conjugates. **e,** Tumors collected on day 7 were dissociated, and tumor cells were assessed for Cy5 fluorescence by flow cytometry (left panel) or lysed and assessed for luciferase activity (right panel). Each bar represents the average value (± S.D.), and each point represents values measured for each independent tumor sample. * P < 0.05; ** P < 0.01; *** P < 0.001; **** P < 0.0001

We further tested si<(EG_18_L)_2_ and si<(EG_18_L_diacid_)_2_ in mice bearing Luc-expressing MDA-MB-231 tumors, using the low dose of 1.5 mg/kg in an attempt to dissect any subtle differences in silencing potency (**Fig. 5d)**. Conjugate si_Luc_ sequences were administered i.v. on days 1, 3, and 5, and again on day 6 using an inactive, Cy5-labeled conjugate. Interestingly, si<(EG_18_L_diacid_)_2_ demonstrated higher renal accumulation and lower liver accumulation compared to si<(EG_18_L)_2_ (**Supplementary Fig. 13)**. Tumors collected on day 7 were assessed by flow cytometry, revealing comparable tumor cell siRNA uptake by si<(EG_18_L)_2_ and si<(EG_18_L_diacid_)_2_ (**Fig. 5e**), and similar levels of Luc knockdown, despite the significantly increased albumin affinity of si<(EG_18_L_diacid_)_2_. These data suggest that, for *in vivo* tumor gene silencing activity, the role of lipid hydrophobicity outweighs the value of increasing albumin affinity beyond the already high-affinity binding of si<(EG_18_L)_2_.

These data led us to pose the hypothesis that the ideal conjugate may selectively hitchhike on albumin in the bloodstream but that this association should be reversible to allow the lipids to participate in cell and endosomal membrane penetration. Further, the conjugate should have sufficient hydrophobicity to promote membrane permeation. We anticipate that albumin-bound conjugate can participate in cell internalization based on high tumor cell albumin uptake and catabolism^9^ but that the efficacy of si<(EG_18_L)_2_ may be at least in part driven by its inherent ability to overcome cell-level barriers. To test this hypothesis, we treated cells with either si<(EG_18_L)_2_ or si<(EG_18_L_diacid_)_2_ in the absence of serum and in the presence of increasing excess of free albumin (**Extended Data Fig. 2a**). This experimental revealed that the si<(EG_18_L)_2_ has inherently higher cell uptake than si<(EG_18_L_diacid_)_2_ likely due to the charged and more hydrophilic diacids reducing membrane insertion. This study also revelated that increasing albumin concentrations reduced si<(EG_18_L)_2_ but not si<(EG_18_L_diacid_)_2_ uptake. This result led us to postulate that albumin-bound siRNAs in general and si<(EG_18_L_diacid_)_2_ (either bound or free form) enter the cell by a form of active endocytosis, while si<(EG_18_L)_2_ does not. To test this hypothesis, we treated cells with both conjugate structures either at 37 or 4°C, with the later condition utilized to inhibit all endocytic processes. Strikingly, cell association of si<(EG_18_L)_2_ was not reduced by the 4°C condition, while it completely abrogated uptake of si<(EG_18_L_diacid_)_2_ (**Extended Data Fig. 2b**). These results indicate that si<(EG_18_L)_2_ has a unique combination of characteristics and balanced level of albumin affinity that contribute to its ability to be optimal for both albumin hitchhiking in circulation and overcoming intracellular barriers.

### Albumin-binding lipophilic siRNA conjugates outperform siRNA directly conjugated to albumin

To further investigate the importance of hydrophobicity and reversibility of albumin binding, we covalently bound the siRNA duplex directly to mouse serum albumin (MSA). The free thiols of MSA were modified with an azido-PEG3-maleimide linker and then reacted with DBCO-modified siRNA duplex (**Extended Data Fig. 3a**). EMSA revealed the expected mobility shift of resulting product relative to the DBCO-duplex precursor (**Extended Data Fig. 3b**). Following i.v. delivery, nearly 80% of the siRNA delivered in the covalent MSA conjugate was found in the albumin-containing plasma fraction (**Extended Data Fig. 3c**). Interestingly, the observed t_1/2_ of the si-covalent-MSA was less than 10 min, far less than the >60 min seen for si<(EG_18_L)_2_ (**Extended Data Fig. 3d**). Kidney accumulation of si-covalent-MSA was similar to what was seen for si<(EG_18_L)_2_ (**Extended Data Fig. 3e**), suggesting that renal clearance is not the primary driver of diminished t_1/2_ for si-covalent-MSA. It is possible that cell surface glycoproteins gp18 and gp30 bind to si-covalent-MSA, trafficking it for lysosomal degradation, as described in previous reports of modified albumin conjugates^61–63^, rather than binding to gp60 which returns unmodified albumin to the circulation. Following delivery to tumor-bearing mice, IVIS and flow cytometry revealed comparable tumor delivery of the two compounds (**Extended Data Fig. 3f-g, Supplementary Fig. 13**). However, the percentage reduction of tumor luciferase activity by si_Luc_-covalent-MSA was only approximately half of that created by si_Luc_<(EG_18_L)_2_ (**Extended Data Fig. 3h**). These data further suggest that reversible albumin binding and presence of the lipid tails of si<(EG_18_L)_2_ contributes to conjugate bioactivity.

The combined results from the diacid and direct albumin conjugate studies suggest that both reversibility of albumin binding and inclusion of a hydrophobic lipid to promote cell uptake are important features of the si<(EG_18_L)_2_ design. Due to the different and somewhat unexpected circulation behavior of the direct conjugate, we sought to further probe this aspect of the design space by using si<(EG18)2 lacking a lipid tail, and also using si<(EG_18_L)_2_ harboring cleavability in the linker that would release the lipid tail; the linkers were designed to be either reducible through presence of a disulfide or to enzymatically degradable by inclusion of a nuclease-susceptible thymidine (**Extended Data Fig. 4a**). Omission of or programmed release of the lipid tail resulted in diminished tumor gene silencing *in vivo* across all conjugates tested, further bolstering the theory that presence of the intact lipid tail is critical for si<(EG_18_L)_2_ activity (**Extended Data Fig. 4b-e**).

### Albumin-Binding siRNA Conjugate Shows Greater Efficacy for TNBC Therapy than a Small Molecule Inhibitor of Mcl-1

To assess the utility of si<(EG_18_L)_2_ for gene targeting in TNBC, HCC70 human TNBC cells were treated in vitro with si<(EG_18_L)_2_ structures based on siRNA sequences directed against *MCL1* (si_MCL1_-L_2_). This system achieved carrier-free, dose-dependent *MCL1* knockdown (**Fig. 6a**) and induced caspase 3/7 activity (**Fig. 6b**), a hallmark of the intrinsic apoptosis pathway and consistent with the phenotypic consequences of MCL-1 loss of function. Similar results in terms of both *MCL1* knockdown and caspase 3/7 induction were found in vitro for si_MCL1_-L_2_-treated HCC1187 and MDA-MB-231 cells (**Supplementary Fig. 14**). Orthotopic HCC70 tumors grown in female athymic mice were treated i.v. with si_MCL1_-L_2_ (10 mg/kg), resulting in potent MCL1 downregulation at treatment day 8, as measured by IHC (**Fig. 6c**), correlating with induction of caspase-3 cleavage and reduction in proliferation (**Fig. 6d, Extended Data Fig. 5a-b**). In parallel, mice bearing HCC70 tumors were treated with doses of 12.5 mg/kg of the MCL-1 small molecule inhibitor MIK665 developed by Servier and Novartis; this doses matches the maximum tolerated dose of this compound^64^ and is more than 18 times the molar quantity of si_MCL1_-L_2_ delivered used. However, MIK665 induced caspase 3 cleavage to a lesser extent and reduced cell proliferation to a lower degree than what was seen for si_MCL1_-L_2_. (**Fig. 6d**). Similar levels of orthotopic tumor *MCL1* silencing and induction of caspase-3 cleavage were also seen in both MDA-MB-231 and HCC1187 tumors at treatment day 8; for the MDA-MB-231 study, comparison to MIK665 treatment was included in the study design and again suggested superiority of si_MCL1_-L_2_ for reducing proliferation and triggering Caspase-3 cleavage within tumors (**Extended Data Fig. 5a-b**).

**Figure 6.**
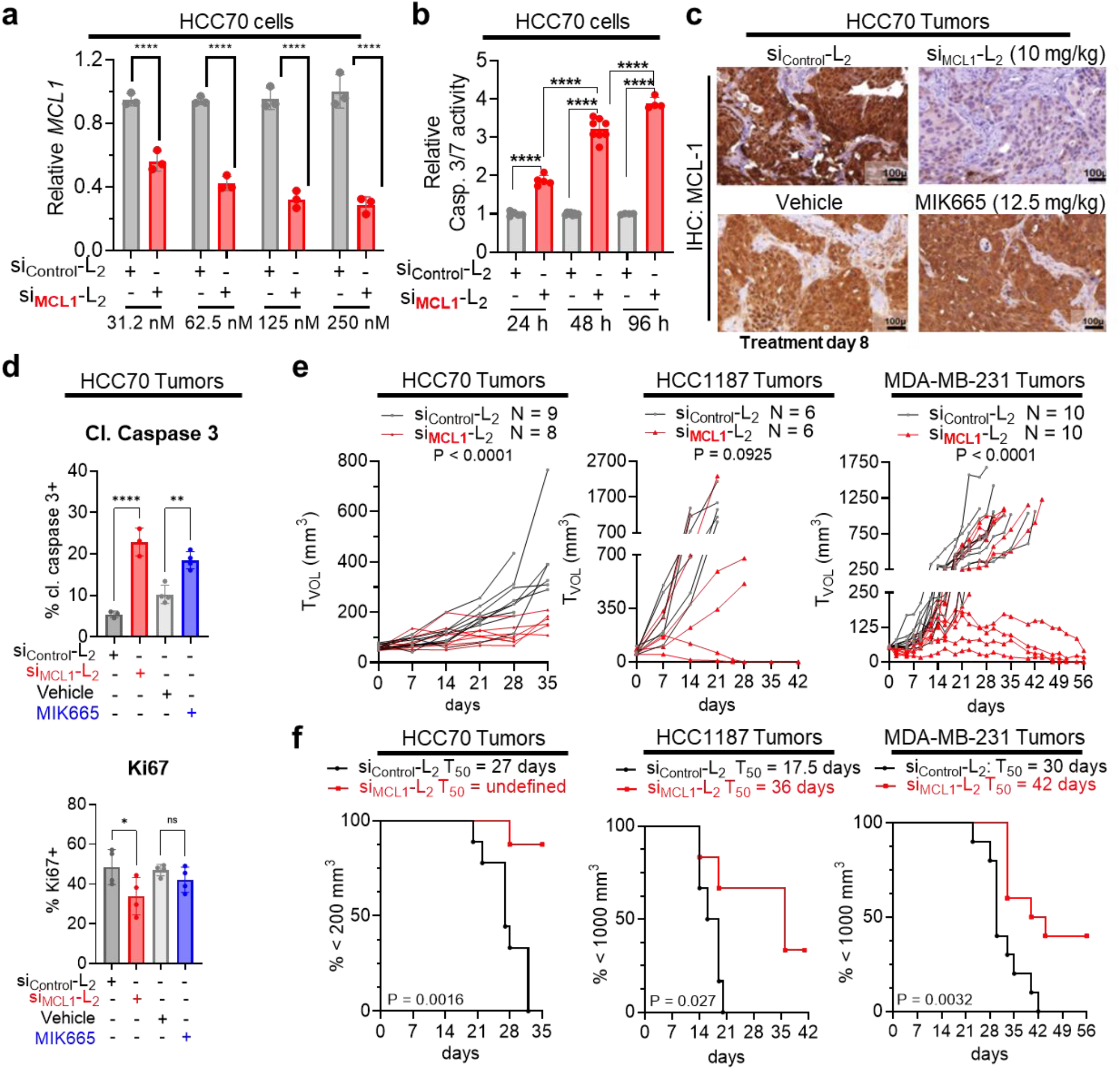
Carrier-free delivery of MCL1-targeting siRNA diminishes MCL-1 expression, tumor cell survival, and tumor growth in TNBC. **a,** HCC70 cells were treated in serum-free OptiMEM with siRNA-lipid conjugates and assessed 48 hours post-treatment for expression of MCL1 transcript levels, corrected for expression of the housekeeping gene PPIB, and are shown relative to MCL1 levels seen in si_Control_-L_2_ (250 nM)-treated cells. **b,** HCC70 cells treated with siRNA-lipid conjugates for 24-96 hours at 100 nM were assessed using Caspase 3/7-Glo. **c-d,** Mice bearing orthotopic HCC70 tumors were treated on day 0 and day 7 with si_Control_-L_2_ or si_MCL1_-L_2_ (10 mg/kg, delivered i.v.). Tumors were collected on day 8. n = 3-4. **c,** Representative images of HCC70 tumors assessed by IHC for MCL-1. **d,** Quantification of the percentage of HCC70 tumor cells staining positive for Ki67 and cleaved (cl.) caspase 3. **e-f,** Mice harboring HCC70 (N = 8-9), HCC1187 (N = 6), and MDA-MB-231 (n = 8-10) orthotopic tumors (50 −100 mm^3^) received once weekly doses of si_Control_-L_2_ or si_MCL1_-L_2_, vehicle (used for delivery of MIK665), or MIK665 through day 28 (HCC70 and HCC1187) and through day 49 (MDA-MB-231) by i.v. injection. Tumors were collected on day 8 (HCC70), day 35 (HCC70), day 42 (HCC1187), and day 56 (MDA-MB-231) or earlier if tumors ulcerated or exceeded humane size limitations. **e,** Tumor volumes were measured throughout treatment. Statistical values were calculated based on area under the curve. **f,** Kaplan-meier analysis of tumor bearing mice, defining survival as tumor volume under 1000 mm^3^ (HCC1187 and MDA-MB-231) or 200 mm^3^ (HCC70, which exhibited slow tumor growth rate). Average time to exceed the defined tumor volume (T_50_) for each group is shown above each panel. P values are calculated using the log-rank (MantelCox) test.

In longer term efficacy studies, weekly si_MCL1_-L_2_ treatment through day 28 inhibited growth in 8/8 HCC70 tumors, 4/6 HCC1187 tumors, and 4/10 MDA-MB-231 as compared to si_Control_-L_2_, leading to overall greater survival (**Fig. 6e-f**). Interestingly, treatment with MIK665 did not alter tumor growth in any HCC70 or MDA-MB-231-bearing mice as compared to those treated with vehicle control (**Supplementary Fig. 15**), revealing a potential therapeutic advantage of MCL-1 knockdown over small molecule MCL-1 inhibition in this setting. Though the reasons underlying these results are uncertain, it is often observed that tumors respond to small molecule MCL-1 inhibition by rapidly upregulating MCL-1 protein^65–69^ translation, potentially increasing resistance to MCL1 inhibition^70^. Importantly, this scenario would be precluded by targeted knockdown of MCL1 transcripts. Further, MCL-1 may have functions unrelated to its BH3-binding domain, which might be carried out unabated by BH3 mimetics like MIK665 but lost upon MCL-1 knockdown^71–73^. Importantly, no significant toxicities were observed in mice treated with si_MCL1_-L_2_, even at the highest dose tested (20 mg/kg), as measured by blood chemistry and CBC, and B cells isolated from the bone marrow and spleen (**Supplementary Figs. 8, 16**). In contrast, toxicities often observed in response to MCL-1 small molecule inhibitors include B-cell and erythrocyte depletion, hemolysis, and weight loss^64^.

## Conclusion

This work shows the optimization of carrier-free, albumin-mediated tumor delivery and bioactivity of siRNA *in vivo*, through systematic variation of lipid-siRNA conjugate valency, linker length, phosphorothioate bonds, lipid chemistry, linker degradability, and linker branching architecture. Modest dosing with this construct achieved nearly 80% carrier-free knockdown in an orthotopic tumor model of triple negative breast cancer, a significant advancement for carrier-free, extra-hepatic RNAi delivery.

In assessing the impact of a hydrophilic linker on albumin binding, we identified an optimal length [si<(EG_18_L)_2_] and branch point placement of the divalent lipid. Notably, designs with the hydrophilic linker before the branching point of the divalent structure, when matched for both overall hydrophobicity or length between the siRNA and lipids, showed inferior albumin association in plasma *in vivo*, which may be attributable to greater lipoprotein association and propensity for self-assembly (lower CMC). Our studies also showed that phosphorothioate, rather than phosphodiester, linkages are beneficial within the linker structure and at the sense strand terminus where the albumin-binding moiety is located, likely contributing to plasma albumin association.

Notably, a combination of selective, but reversible albumin binding in the circulation, combined with an optimal level of hydrophobicity to promote cell uptake whenever the conjugate dissociates from albumin after reaching the tumor site, is critical to the overall performance of si<(EG_18_L)_2_. To further support this point, it was shown that both direct albumin conjugation and conjugates with a degradable linker that can release the C18 moieties have reduced tumor gene silencing function.

Finally, si<(EG_18_L)_2_ was shown to provide significant knockdown and tumor growth inhibition in vivo through silencing of oncogenic MCL-1 in three models of TNBC; the siRNA was also found to outperform a clinically tested MCL-1 small molecule inhibitor given at its MTD on the same treatment schedule. Of critical importance, little MCL-1 knockdown was observed in livers of treated mice, and no toxicities were observed with control or MCL1-directed sequences.

Overall, this work has important implications for the delivery of siRNAs to extrahepatic targets, a goal that has remained clinically elusive. Albumin is known to accumulate at sites of inflammation and vascular leakiness, which are associated with a wide array of diseases. Therefore, insights gleaned from si<(EG_18_L)_2_ development for *in situ* albumin carrier-free delivery will support development and optimization of extrahepatic, carrier-free siRNA therapeutics in cancer treatment and beyond.

## Data Availability

The raw/processed data required to reproduce these findings are available upon request.

## Funding

This work was supported by the National Institutes of Health (R01 CA224241, R01 EB019409, R01 CA260958, R21 AR078636, T32 CA217834, F32 CA268705) and the National Science Foundation (BMAT 1349604).

## Competing Interests Statement

The authors declare that they have no known competing financial interests or personal relationships that could have appeared to influence the work reported in this paper.

---

**Extended Data Figure 1.**
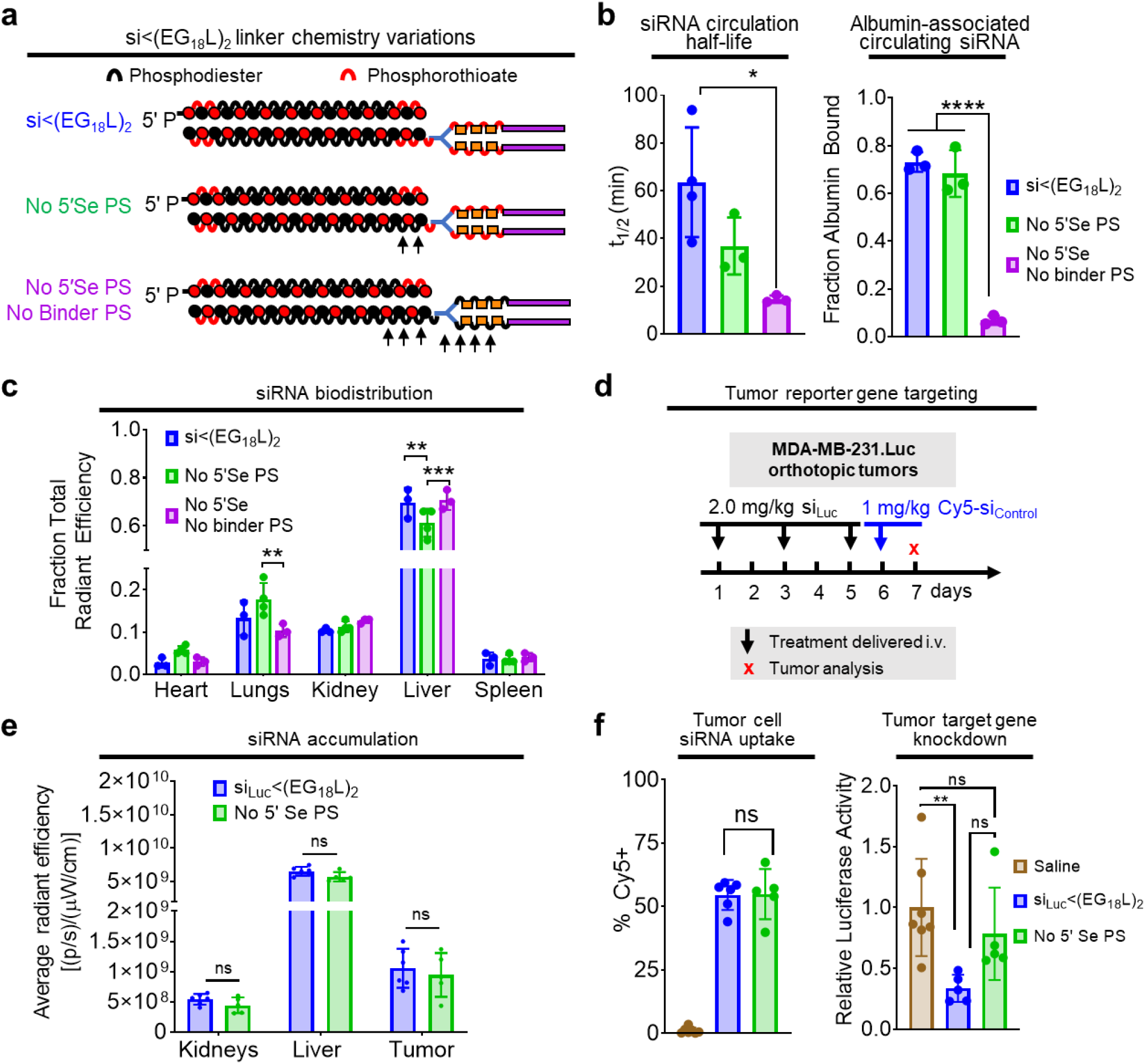
Phosphorothioate linkages on the 5’ end of the sense strand and within the linker region contribute to albumin association in plasma and tumor gene silencing activity. **a,** Schematic structural representations of the siRNA-lipid conjugates with varied inclusion of PS bonds. **b,** Cy5-labeled lipid-siRNA conjugates were delivered i.v. to mice at 1 mg/kg. Intravascular Cy5 fluorescence measured through approximately 1h following delivery was used to calculate siRNA circulation half-life (left panel). Size exclusion chromatographic calculation of fraction of Cy5 fluorescence associated with albumin-containing plasma fractions (right panel). **c,** Cy5 fluorescence in kidneys, lungs, liver, spleen and heart was measured 1h after treatment. Fluorescence values in each organ relative to the total fluorescent values of all organs combined is shown. **d-f,** Mice bearing intra-mammary MDA-MB-231.Luc tumors were treated i.v. with siRNA-lipid conjugates, using siRNA sequences against luciferase (si_Luc_). **d,** Schematic showing treatment schedule for i.v. delivery of siRNA-lipid conjugates. **e,** Cy5 fluorescence in kidneys, liver and tumor for each mouse was measured 18h after treatment. **f,** Tumors were collected on day 7 and either dissociated to assess for Cy5 fluorescence by flow cytometry (left panel) or lysed to assess for luciferase activity (right panel). Bars show the average (± S.D.) value per group, and points are the values for each tumor analyzed. All relative values shown are calculated relative to the average value of saline-treated controls, which was set to a value of 1. Each bar represents the average value (± S.D.), and each point represents values measured for each individual sample, For B and C, N = 3-4. For E and F, N=5-6. * P < 0.05; ** P < 0.01; **** P < 0.0001

**Extended Data Figure 2:**
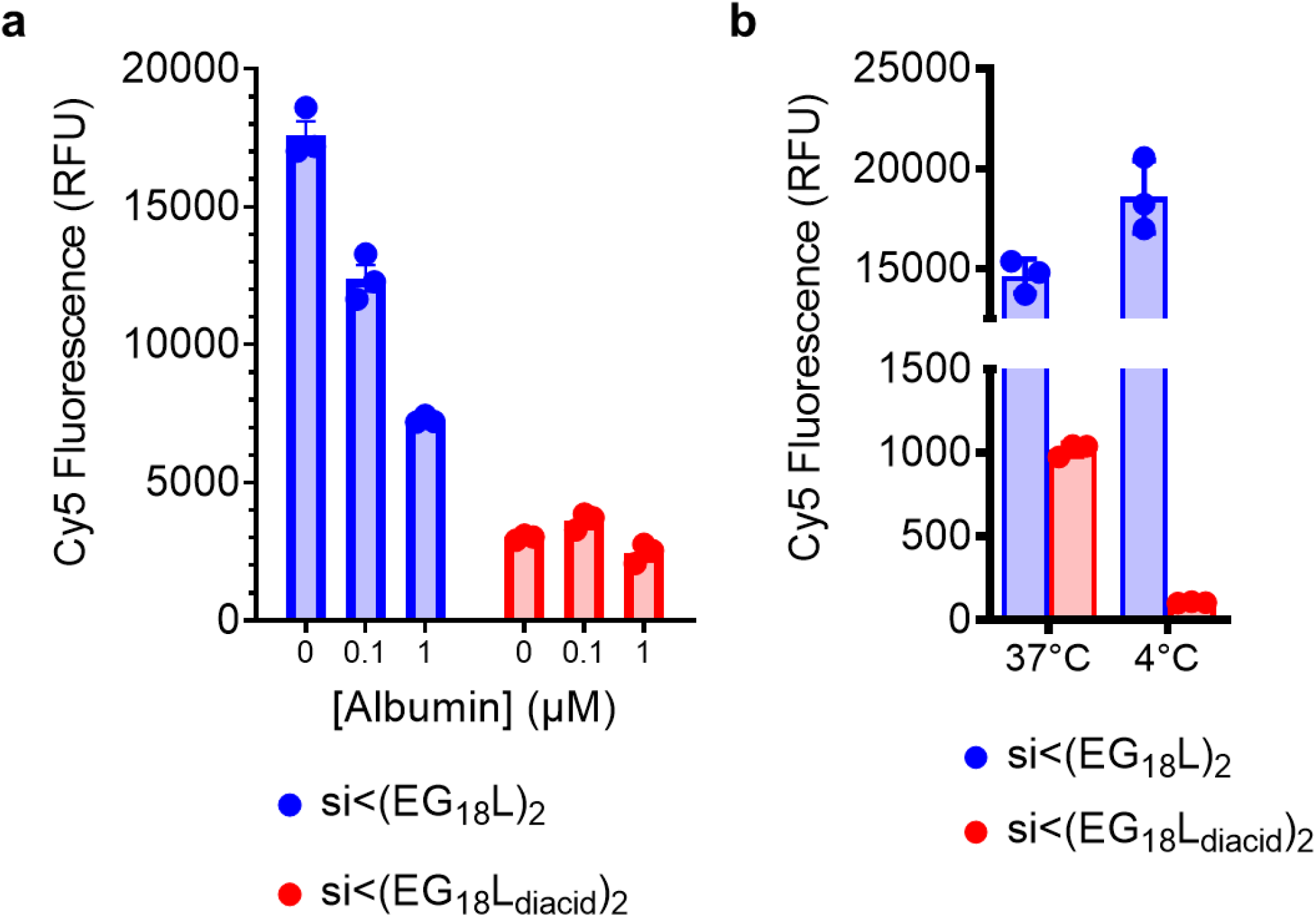
Cellular association of si<(EG_18_L)_2_ is driven by hydrophobicity and does not depend on active endocytosis. **a,** Cy5 fluorescence measured by flow cytometry of MDA-MB-231 cells incubated with 100 nM Cy5-labeled si<(EG_18_L)_2_ or si<(EG_18_L_diacid_)_2_ in serum-free media, with or without albumin, for 4h at 37°C. **b,** Cy5 fluorescence measured by flow cytometry of MDA-MB-231 cells incubated with 100 nM Cy5-labeled si<(EG_18_L)_2_ or si<(EG_18_L_diacid_)_2_ in Opti-MEM for 2 h at 37°C vs. 4°C.

**Extended Data Figure 3.**
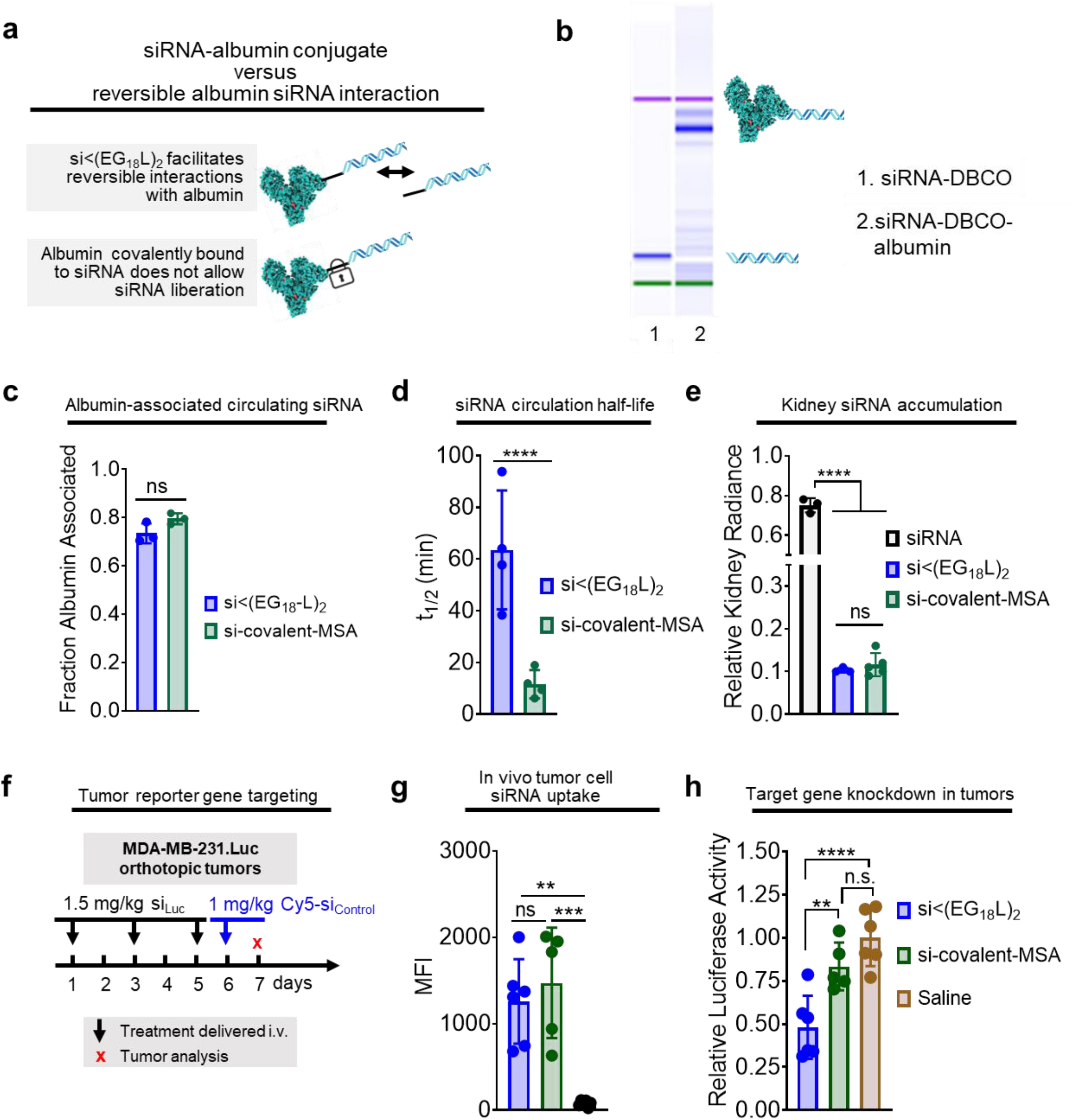
Reversible siRNA interaction with albumin, enabled by siRNA-lipid conjugation, enhances siRNA circulation and in vivo bioactivity. **a,** Schematic model describing how siRNA conjugated to lipid enables reversible albumin interactions, as opposed to the irreversible nature of siRNA covalently linked with albumin. **b,** Confirmation of conjugation of siRNA to albumin visualized using an Agilent 2100 Bioanalyzer. **c,** Cy5-labeled siRNA-lipid conjugates or siRNA-albumin were delivered (1 mg/kg). Size exclusion chromatographic measurements of Cy5 fluorescence in albumin-containing plasma fraction. **d,** Intravascular Cy5 fluorescence measured through approximately 1h was used to calculate siRNA circulation half-life. **e,** Cy5 fluorescence in kidneys, lungs, liver, spleen and heart was measured 1h after treatment. Fluorescence values in kidneys relative to the total fluorescent values of all organs combined is shown (left panel). **f,** Schematic showing treatment schedule for i.v. delivery of siRNA-lipid conjugates to tumor-bearing mice. **g,** Dissociated tumors collected on treatment day 7 were assessed for Cy5 fluorescence by flow cytometry or H. lysed and assessed for luciferase activity. For panels C-E,G, and H, each bar represents the average value (± S.D.), each point represents values measured for each individual sample. * P < 0.05; ** P < 0.01; *** P < 0.001; **** P < 0.0001

**Extended Data Figure 4:**
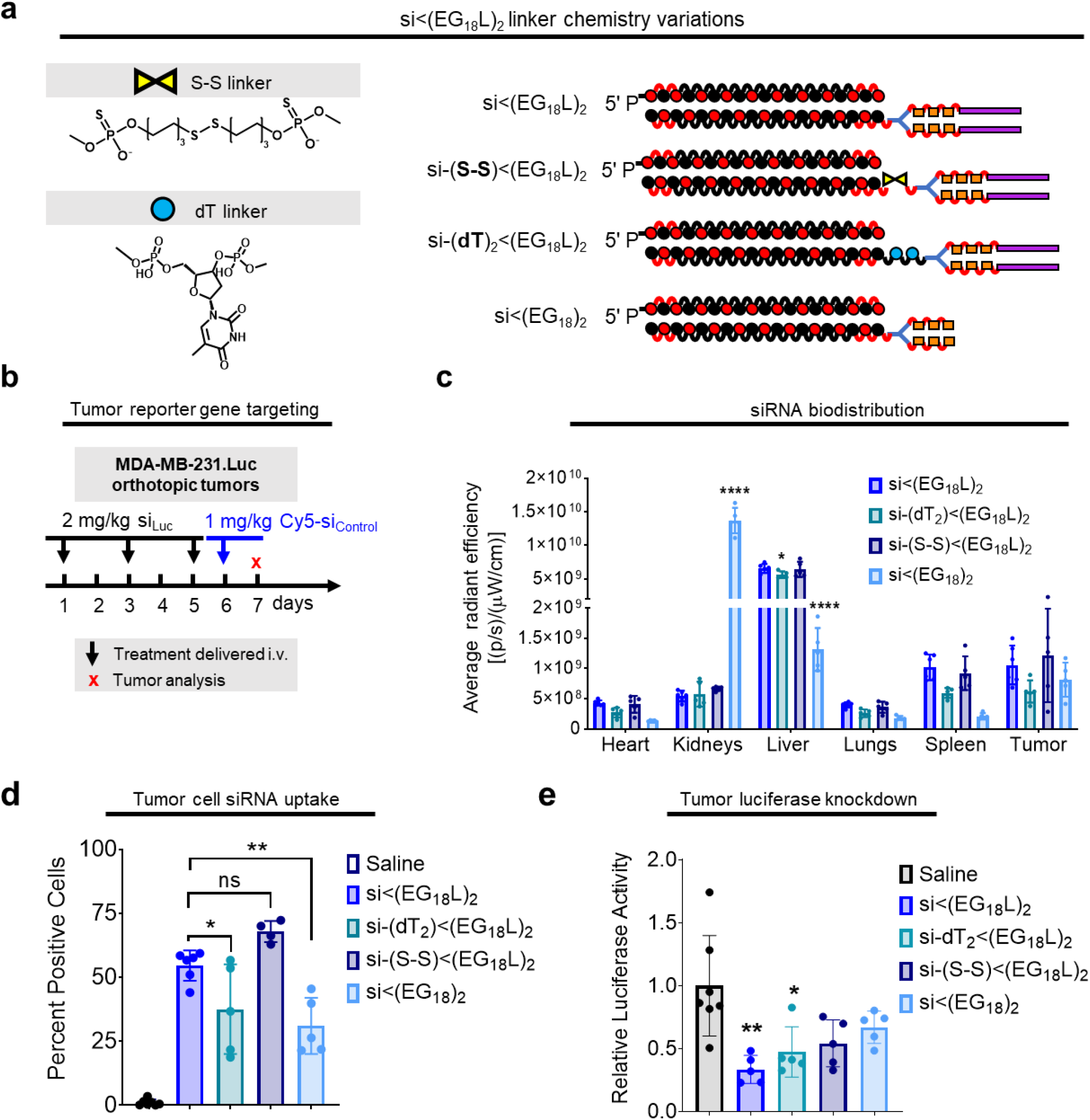
Interrogation of siRNA conjugate lipid tail presence. **a,** Schematic structural representations of the si<(EG_18_L_2_) variants. **b,** Treatment schematic showing i.v. dosing of conjugates to tumor bearing mice. **c,** Biodistribution of Cy5-labeled siRNA conjugates measured by epifluorescence 18 h after 1 mg/kg intravenous injection. Significance was assessed by 2-way ANOVA with Tukey’s multiple comparisons (n=4-6). Significance compared to si<(EG_18_L)_2_ displayed. **d,** Percent Cy5 positive tumor cells determined by flow cytometry **e,** Relative luciferase activity of tumor lysates after 3 x 2 mg/kg intravenous injections of siRNA conjugates targeting firefly luciferase over the course of one week. Significance assessed by 1-way ANOVA with Tukey’s multiple comparisons (n=4-7). Significance compared to saline control displayed.

**Extended Data Figure 5.**
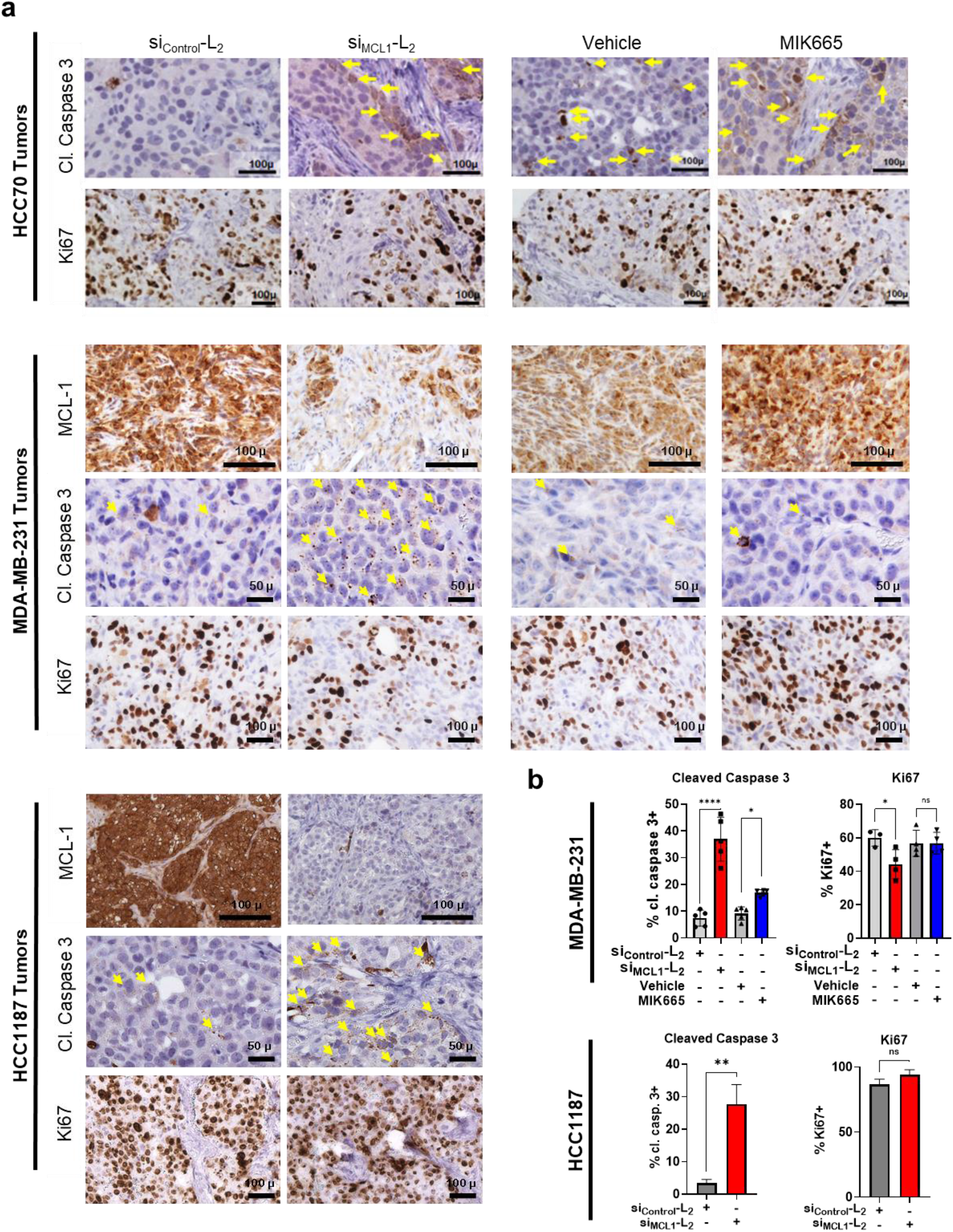
Carrier-free delivery of MCL1-targeting siRNA diminishes MCL-1 expression and tumor cell survival in TNBC. **a,** Mice bearing orthotopic HCC70, HCC1187, or MDA-MB-231 tumors were treated on day 0 and day 7 with si_Control_-L_2_ or si_MCL1_-L_2_ (10 mg/kg, delivered i.v.). Tumors were collected on day 8. N = 3-4. Representative images of immunohistochemical staining for cleaved caspase 3, Ki67, and MCL-1 are shown. **b,** Quantification of the percentage of HCC1187 and MDA-MB-231 tumor cells staining positive for Ki67 and cleaved (cl.) caspase 3 is shown. (HCC70 quantitation is shown in Figure 6.)

## Notes

### Competing Interest Statement

The authors have declared no competing interest.

## References

1. Zhang, X. et al. Patisiran Pharmacokinetics, Pharmacodynamics, and Exposure-Response Analyses in the Phase 3 APOLLO Trial in Patients With Hereditary Transthyretin-Mediated (hATTR) Amyloidosis. J Clin Pharmacol 60, 37–49 (2020).

2. Jackson, M.A. et al. Kupffer cell release of platelet activating factor drives dose limiting toxicities of nucleic acid nanocarriers. Biomaterials 268, 120528 (2021).

3. Kedmi, R., Ben-Arie, N. & Peer, D. The systemic toxicity of positively charged lipid nanoparticles and the role of Toll-like receptor 4 in immune activation. Biomaterials 31, 6867–6875 (2010).

4. Urits, I. et al. A Review of Patisiran (ONPATTRO^®^) for the Treatment of Polyneuropathy in People with Hereditary Transthyretin Amyloidosis. Neurol Ther 9, 301–315 (2020).

5. He, H., Liu, L., Morin, E.E., Liu, M. & Schwendeman, A. Survey of Clinical Translation of Cancer Nanomedicines-Lessons Learned from Successes and Failures. Acc Chem Res 52, 2445–2461 (2019).

6. Nichols, J.W. & Bae, Y.H. EPR: Evidence and fallacy. J Control Release 190, 451–464 (2014).

7. Danhier, F. To exploit the tumor microenvironment: Since the EPR effect fails in the clinic, what is the future of nanomedicine? Journal of Controlled Release 244, 108–121 (2016).

8. Nakamura, Y., Mochida, A., Choyke, P.L. & Kobayashi, H. Nanodrug Delivery: Is the Enhanced Permeability and Retention Effect Sufficient for Curing Cancer? Bioconjug Chem 27, 2225–2238 (2016).

9. Hoogenboezem, E.N. & Duvall, C.L. Harnessing albumin as a carrier for cancer therapies. Adv Drug Deliv Rev 130, 73–89 (2018).

10. Matsumura, Y. & Maeda, H. A new concept for macromolecular therapeutics in cancer chemotherapy: mechanism of tumoritropic accumulation of proteins and the antitumor agent smancs. Cancer research 46, 6387–6392 (1986).

11. Stehle, G. et al. Plasma protein (albumin) catabolism by the tumor itself--implications for tumor metabolism and the genesis of cachexia. Crit Rev Oncol Hematol 26, 77–100 (1997).

12. Wunder, A. et al. Enhanced albumin uptake by rat tumors. Int J Oncol 11, 497–507 (1997).

13. Schilling, U. et al. Design of compounds having enhanced tumour uptake, using serum albumin as a carrier--Part II. In vivo studies. Int J Rad Appl Instrum B 19, 685–695 (1992).

14. Andersson, C., Iresjö, B.M. & Lundholm, K. Identification of tissue sites for increased albumin degradation in sarcoma-bearing mice. J Surg Res 50, 156–162 (1991).

15. Qin, S. et al. A physiological perspective on the use of imaging to assess the in vivo delivery of therapeutics. Ann Biomed Eng 42, 280–298 (2014).

16. Campbell, P.M. & Der, C.J. Oncogenic Ras and its role in tumor cell invasion and metastasis. Seminars in Cancer Biology 14, 105–114 (2004).

17. Commisso, C. et al. Macropinocytosis of protein is an amino acid supply route in Ras-transformed cells. Nature 497, 633–637 (2013).

18. Kamphorst, J.J. et al. Human pancreatic cancer tumors are nutrient poor and tumor cells actively scavenge extracellular protein. Cancer Research 75, 544–553 (2015).

19. Kratz, F. Albumin as a drug carrier: design of prodrugs, drug conjugates and nanoparticles. J Control Release 132, 171–183 (2008).

20. Yu, K.D. et al. Identification of prognosis-relevant subgroups in patients with chemoresistant triple-negative breast cancer. Clin Cancer Res 19, 2723–2733 (2013).

21. Bolomsky, A. et al. MCL-1 inhibitors, fast-lane development of a new class of anti-cancer agents. J Hematol Oncol 13, 173 (2020).

22. Wang, H., Guo, M., Wei, H. & Chen, Y. Targeting MCL-1 in cancer: current status and perspectives. J Hematol Oncol 14, 67 (2021).

23. Sarett, S.M. et al. Lipophilic siRNA targets albumin in situ and promotes bioavailability, tumor penetration, and carrier-free gene silencing. Proceedings of the National Academy of Sciences of the United States of America 114, e6490–E6497 (2017).

24. Prakash, T.P. et al. Fatty acid conjugation enhances potency of antisense oligonucleotides in muscle. Nucleic Acids Res 47, 6029–6044 (2019).

25. Coutinho, P.J.G., Castanheira, E.M.S., Rei, M.C. & Oliveira, M.E.C.D.R. Nile Red and DCM Fluorescence Anisotropy Studies in C12E7/DPPC Mixed Systems. The Journal of Physical Chemistry B 106, 12841–12846 (2002).

26. Jackson, M.A. et al. Dual carrier-cargo hydrophobization and charge ratio optimization improve the systemic circulation and safety of zwitterionic nano-polyplexes. Biomaterials 192, 245–259 (2019).

27. Allerson, C.R. et al. Fully 2’-modified oligonucleotide duplexes with improved in vitro potency and stability compared to unmodified small interfering RNA. J Med Chem 48, 901–904 (2005).

28. Osborn, M.F. et al. Hydrophobicity drives the systemic distribution of lipid-conjugated siRNAs via lipid transport pathways. Nucleic Acids Res 47, 1070–1081 (2019).

29. Wolfrum, C. et al. Mechanisms and optimization of in vivo delivery of lipophilic siRNAs. Nat Biotechnol 25, 1149–1157 (2007).

30. Pires, L.A. et al. Use of cholesterol-rich nanoparticles that bind to lipoprotein receptors as a vehicle to paclitaxel in the treatment of breast cancer: pharmacokinetics, tumor uptake and a pilot clinical study. Cancer Chemother Pharmacol 63, 281–287 (2009).

31. Vitols, S., Peterson, C., Larsson, O., Holm, P. & Aberg, B. Elevated uptake of low density lipoproteins by human lung cancer tissue in vivo. Cancer Res 52, 6244–6247 (1992).

32. Mooberry, L.K., Nair, M., Paranjape, S., McConathy, W.J. & Lacko, A.G. Receptor mediated uptake of paclitaxel from a synthetic high density lipoprotein nanocarrier. J Drug Target 18, 53–58 (2010).

33. Biscans, A., Coles, A., Echeverria, D. & Khvorova, A. The valency of fatty acid conjugates impacts siRNA pharmacokinetics, distribution, and efficacy in vivo. J Control Release 302, 116–125 (2019).

34. Hvam, M.L. et al. Fatty Acid-Modified Gapmer Antisense Oligonucleotide and Serum Albumin Constructs for Pharmacokinetic Modulation. Molecular Therapy 25, 1710–1717 (2017).

35. Lacroix, A., Edwardson, T.G.W., Hancock, M.A., Dore, M.D. & Sleiman, H.F. Development of DNA Nanostructures for High-Affinity Binding to Human Serum Albumin. J Am Chem Soc 139, 7355–7362 (2017).

36. Liu, H. et al. Structure-based programming of lymph-node targeting in molecular vaccines. Nature 507, 519–522 (2014).

37. Madsen, K. et al. Structure-Activity and Protraction Relationship of Long-Acting Glucagon-like Peptide-1 Derivatives: Importance of Fatty Acid Length, Polarity, and Bulkiness. Journal of Medicinal Chemistry 50, 6126–6132 (2007).

38. Jeong, J.H., Mok, H., Oh, Y.K. & Park, T.G. siRNA conjugate delivery systems. Bioconjug Chem 20, 5–14 (2009).

39. Shmushkovich, T. et al. Functional features defining the efficacy of cholesterol-conjugated, self-deliverable, chemically modified siRNAs. Nucleic Acids Res 46, 10905–10916 (2018).

40. Osborn, M.F. & Khvorova, A. Improving siRNA Delivery In Vivo Through Lipid Conjugation. Nucleic Acid Ther 28, 128–136 (2018).

41. Lau, J. et al. Discovery of the Once-Weekly Glucagon-Like Peptide-1 (GLP-1) Analogue Semaglutide. Journal of Medicinal Chemistry 58, 7370–7380 (2015).

42. Liu, H. et al. DNA-based micelles: synthesis, micellar properties and size-dependent cell permeability. Chemistry 16, 3791–3797 (2010).

43. Chen, T. et al. DNA micelle flares for intracellular mRNA imaging and gene therapy. Angew Chem Int Ed Engl 52, 2012–2016 (2013).

44. Patwa, A., Gissot, A., Bestel, I. & Barthélémy, P. Hybrid lipid oligonucleotide conjugates: synthesis, self-assemblies and biomedical applications. Chem Soc Rev 40, 5844–5854 (2011).

45. Ercole, F., Whittaker, M.R., Quinn, J.F. & Davis, T.P. Cholesterol Modified Self-Assemblies and Their Application to Nanomedicine. Biomacromolecules 16, 1886–1914 (2015).

46. Yang, F. et al. Effect of human serum albumin on drug metabolism: structural evidence of esterase activity of human serum albumin. J Struct Biol 157, 348–355 (2007).

47. Klein, D. et al. Centyrin ligands for extrahepatic delivery of siRNA. Mol Ther 29, 2053–2066 (2021).

48. Dassie, J., McNamara Ii, J. & Giangrande, P. Systemic administration of optimize aptamer-siRNA chimeras promotes regression of PSMA-expressing tumors. nature biotechnology 27(2009).

49. Yu, Z. et al. Antibody-siRNA conjugates (ARCs) using multifunctional peptide as a tumor enzyme cleavable linker mediated effective intracellular delivery of siRNA. Int J Pharm 606, 120940 (2021).

50. Cen, B. et al. An Efficient Bivalent Cyclic RGD-PIK3CB siRNA Conjugate for Specific Targeted Therapy against Glioblastoma In Vitro and In Vivo. Mol Ther Nucleic Acids 13, 220–232 (2018).

51. He, S. et al. A tumor-targeting cRGD-EGFR siRNA conjugate and its anti-tumor effect on glioblastoma in vitro and in vivo. Drug Deliv 24, 471–481 (2017).

52. Chernikov, I.V. et al. Cholesterol-Containing Nuclease-Resistant siRNA Accumulates in Tumors in a Carrier-free Mode and Silences MDR1 Gene. Molecular Therapy - Nucleic Acids 6, 209–220 (2017).

53. Nair, J.K. et al. Impact of enhanced metabolic stability on pharmacokinetics and pharmacodynamics of GalNAc-siRNA conjugates. Nucleic Acids Res 45, 10969–10977 (2017).

54. Biscans, A. et al. The chemical structure and phosphorothioate content of hydrophobically modified siRNAs impact extrahepatic distribution and efficacy. Nucleic Acids Res 48, 7665–7680 (2020).

55. Hyjek-Składanowska, M. et al. Origins of the Increased Affinity of Phosphorothioate-Modified Therapeutic Nucleic Acids for Proteins. J Am Chem Soc 142, 7456–7468 (2020).

56. Geary, R.S., Norris, D., Yu, R. & Bennett, C.F. Pharmacokinetics, biodistribution and cell uptake of antisense oligonucleotides. Adv Drug Deliv Rev 87, 46–51 (2015).

57. Vollhardt, D. Effect of unsaturation in fatty acids on the main characteristics of Langmuir monolayers. Journal of Physical Chemistry C 111, 6805–6812 (2007).

58. Alberts, B. et al. in Molecular Biology of the Cell. 4th edition (Garland Science, 2002).

59. Ashbrook, J.D., Spector, A.A., Santos, E.C. & Fletcher, J.E. Long chain fatty acid binding to human plasma albumin. J Biol Chem 250, 2333–2338 (1975).

60. Birkett, D.J., Myers, S.P. & Sudlow, G. Effects of Fatty Acids on Two Specific Drug Binding Sites on Human Serum Albumin. Molecular Pharmacology 13, 987–992 (1977).

61. Schnitzer, J.E., Sung, A., Horvat, R. & Bravo, J. Preferential interaction of albumin-binding proteins, gp30 and gp18, with conformationally modified albumins. Presence in many cells and tissues with a possible role in catabolism. J Biol Chem 267, 24544–24553 (1992).

62. Schnitzer, J.E. & Bravo, J. High affinity binding, endocytosis, and degradation of conformationally modified albumins. Potential role of gp30 and gp18 as novel scavenger receptors. J Biol Chem 268, 7562–7570 (1993).

63. Schnitzer, J.E. & Oh, P. Albondin-mediated capillary permeability to albumin. Differential role of receptors in endothelial transcytosis and endocytosis of native and modified albumins. J Biol Chem 269, 6072–6082 (1994).

64. Brennan, M.S. et al. Humanized. Blood 132, 1573–1583 (2018).

65. Xiao, Y. et al. MCL-1 Is a Key Determinant of Breast Cancer Cell Survival: Validation of MCL-1 Dependency Utilizing a Highly Selective Small Molecule Inhibitor. Mol Cancer Ther 14, 1837–1847 (2015).

66. Leverson, J.D. et al. Potent and selective small-molecule MCL-1 inhibitors demonstrate on-target cancer cell killing activity as single agents and in combination with ABT-263 (navitoclax). Cell Death Dis 6, e1590 (2015).

67. Kotschy, A. et al. The MCL1 inhibitor S63845 is tolerable and effective in diverse cancer models. Nature 538, 477–482 (2016).

68. Rysanek, D. et al. Synergism of BCL-2 family inhibitors facilitates selective elimination of senescent cells. Aging (Albany NY) 14, 6381–6414 (2022).

69. Anderson, G.R. et al. PIK3CA mutations enable targeting of a breast tumor dependency through mTOR-mediated MCL-1 translation. Sci Transl Med 8, 369ra175 (2016).

70. Widden, H. & Placzek, W.J. The multiple mechanisms of MCL1 in the regulation of cell fate. Commun Biol 4, 1029 (2021).

71. Rasmussen, M.L. et al. A Non-apoptotic Function of MCL-1 in Promoting Pluripotency and Modulating Mitochondrial Dynamics in Stem Cells. Stem Cell Reports 10, 684–692 (2018).

72. Perciavalle, R.M. et al. Anti-apoptotic MCL-1 localizes to the mitochondrial matrix and couples mitochondrial fusion to respiration. Nat Cell Biol 14, 575–583 (2012).

73. Morciano, G. et al. Mcl-1 involvement in mitochondrial dynamics is associated with apoptotic cell death. Mol Biol Cell 27, 20–34 (2016).

