## Supplemental Information for "Structural Optimization of siRNA Conjugates for Albumin Binding Achieves Effective MCL1-Targeted Cancer Therapy"

### Supplementary Information:

**Supplementary Table 1: siRNA Sequence Details**

| | **Sequence (5' to 3')** |
|---|---|
| Luciferase (Luc) Antisense | (PHO) (OMeU)*(fU)*(OMeC) (fA) (OMeU) (fU) (OMeA) (fU) (OMeC) (fA) (OMeG) (fU) (OMeG) (fC) (OMeA) (fA) (OMeU)*(fU)*(OMeG) |
| Luc Sense | (fC)*(OMeA)*(fA) (OMeU) (fU) (OMeG) (fC) (OMeA) (fC) (OMeU) (fG) (OMeA) (fU) (OMeA) (fA) (OMeU) (fG)*(OMeA)*(fA) |
| Scrambled (Scr) Antisense | (PHO) (OMeG)*(fU)*(OMeA) (fU) (OMeU) (fA) (OMeU) (fA) (OMeC) (fG) (OMeC) (fG) (OMeA) (fU) (OMeU) (fA) (OMeA) (fC) (OMeG)*(fA)*(OMeC) |
| Scr Sense | (fC)*(OMeG)*(fU) (OMeU) (fA) (OMeA) (fU) (OMeC) (fG) (OMeC) (fG) (OMeU) (fA) (OMeU) (fA) (OMeA) (fU)*(OMeA)*(fC) |
| Cy5-labeled Luc Antisense | (Cy5) (OMeU)*(fU)*(OMeC) (fA) (OMeU) (fU) (OMeA) (fU) (OMeC) (fA) (OMeG) (fU) (OMeG) (fC) (OMeA) (fA) (OMeU)*(fU)*(OMeG) |
| Biotin-TEG-labeled Luc Antisense | (BTN-TEG)*(OMeU)*(fU)*(OMeC) (fA) (OMeU) (fU) (OMeA) (fU) (OMeC) (fA) (OMeG) (fU) (OMeG) (fC) (OMeA) (fA) (OMeU)*(fU)*(OMeG) |
| Mcl-1 Antisense | (fC)*(OMeA)*(fU)(OMeC)(fG)(OMeA)(fA)(OMeC)(fC)(OMeA)(fU)(OMeU)(fA)(OMeG)(fC)(OMeA)(fG)*(OMeA)*(fA) |
| Mcl-1 Sense | (PHO)(OMeU)*(fU)*(OMeC) (fU)(OMeG)(fC) (OMeU)(fA)(OMeA) (fU)(OMeG)(fG) (OMeU)(fU)(OMeC) (fG)(OMeA)*(fU)*(OMeG) |
| *Phosphorothioate bond (X)*(X)*<br>*Phosphodiester bond (X) (X)*<br>*2'F substituted base (fX)*<br>*2'OMe substituted base (OMeX)* | |

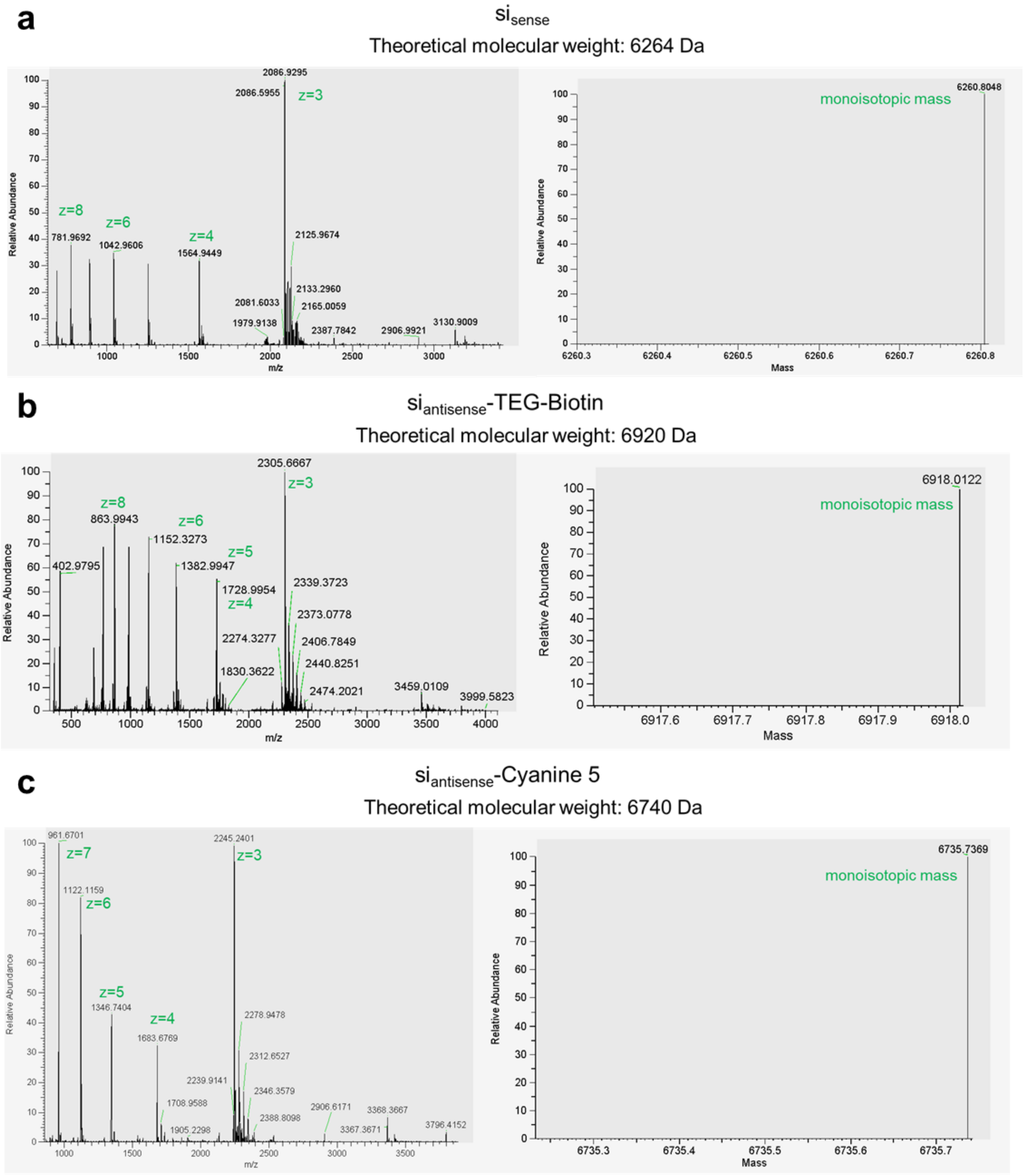
a
$si_{sense}$
Theoretical molecular weight: 6264 Da
2086.9295
2086.5955
z=3
z=8
z=6
z=4
781.9692
1042.9606
1564.9449
2125.9674
2133.2960
2081.6033
2165.0059
1979.9138
2387.7842
2906.9921
3130.9009
Relative Abundance
m/z
6260.8048
monoisotopic mass
Mass
b
$si_{antisense}$-TEG-Biotin
Theoretical molecular weight: 6920 Da
2305.6667
z=3
z=8
863.9943
z=6
1152.3273
z=5
402.9795
1382.9947
1728.9954
z=4
2339.3723
2373.0778
2274.3277
2406.7849
2440.8251
1830.3622
2474.2021
3459.0109
3999.5823
Relative Abundance
m/z
6918.0122
monoisotopic mass
Mass
c
$si_{antisense}$-Cyanine 5
Theoretical molecular weight: 6740 Da
961.6701
z=7
2245.2401
z=3
1122.1159
z=6
z=5
1346.7404
z=4
1683.6769
2278.9478
2312.6527
2239.9141
2346.3579
1708.9588
2906.6171
3368.3667
2388.8098
3367.3671
1905.2298
3796.4152
Relative Abundance
m/z
6735.7369
monoisotopic mass
Mass

**d**

si-$L_1$

Theoretical molecular weight: 6612 Da

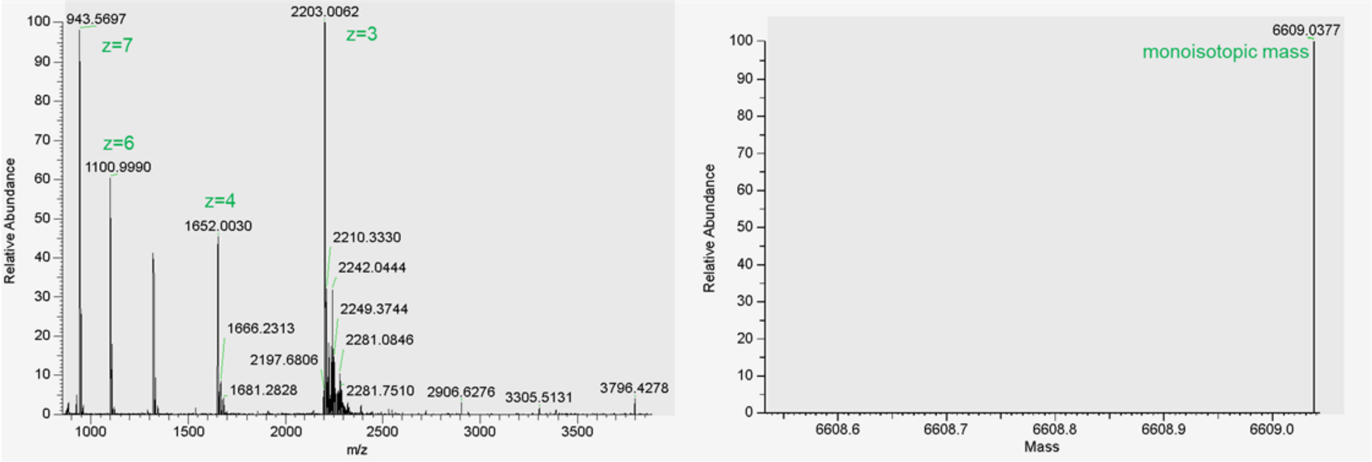


**e**

si-$EG_3$-chol

Theoretical molecular weight: 6963 Da

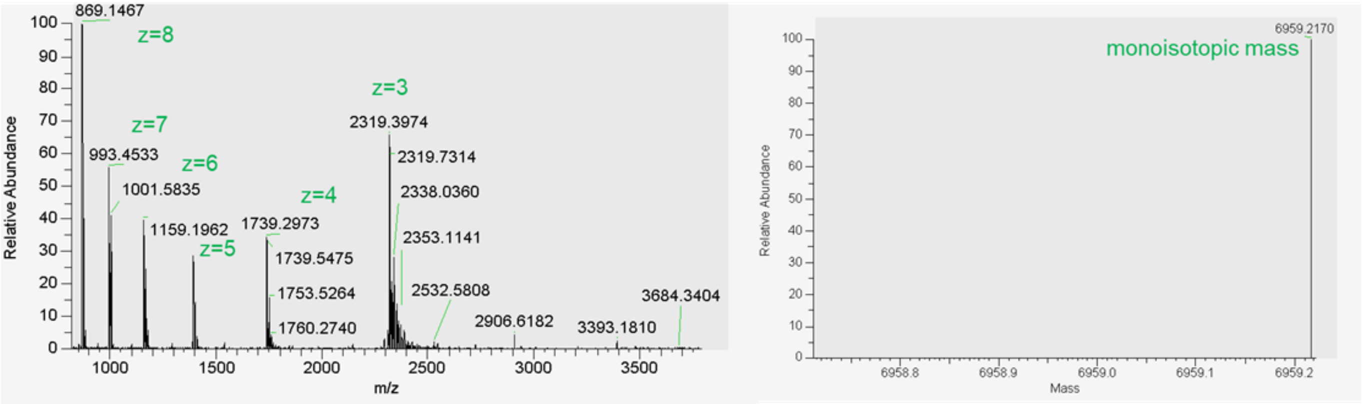


**f**

si<($EG_0L$)$_2$

Theoretical molecular weight: 7126 Da

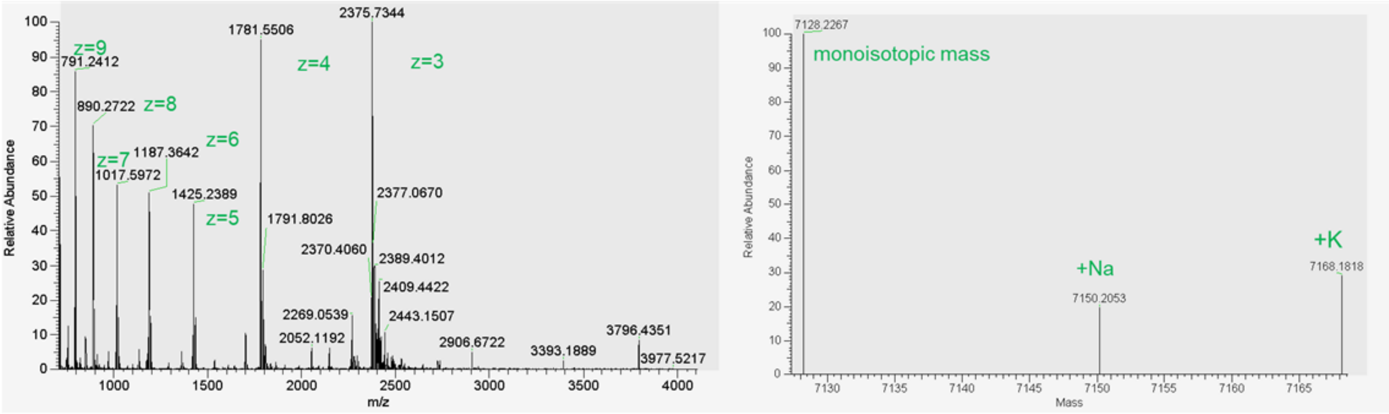

**g**

si<$(EG_6L)_2$

Theoretical molecular weight: 7846 Da

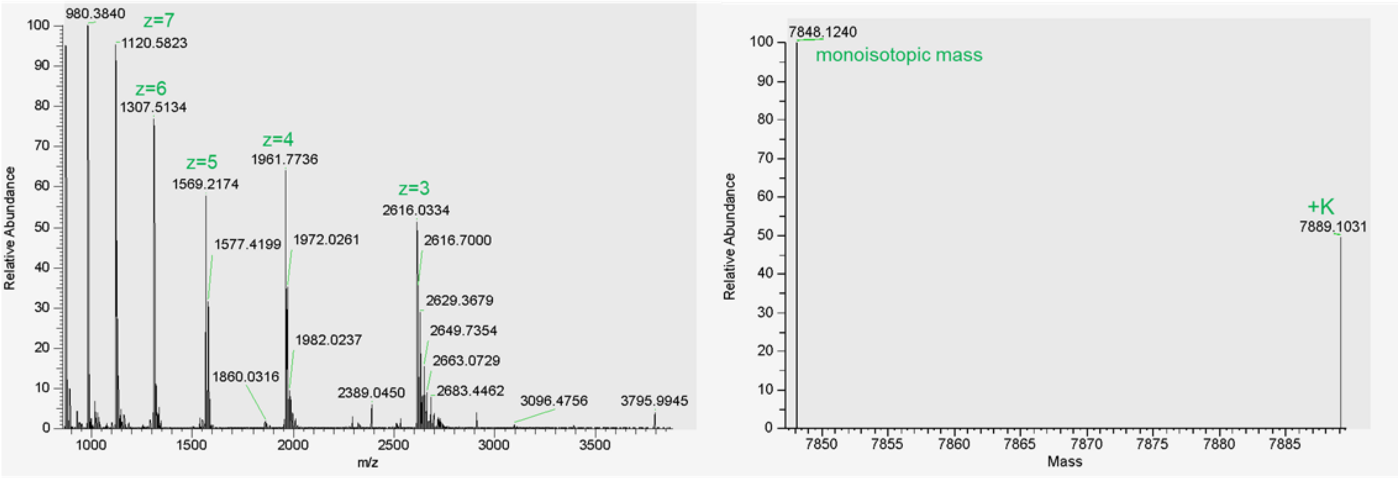


**h**

si<$(EG_6L)_2$

Theoretical molecular weight: 7846 Da

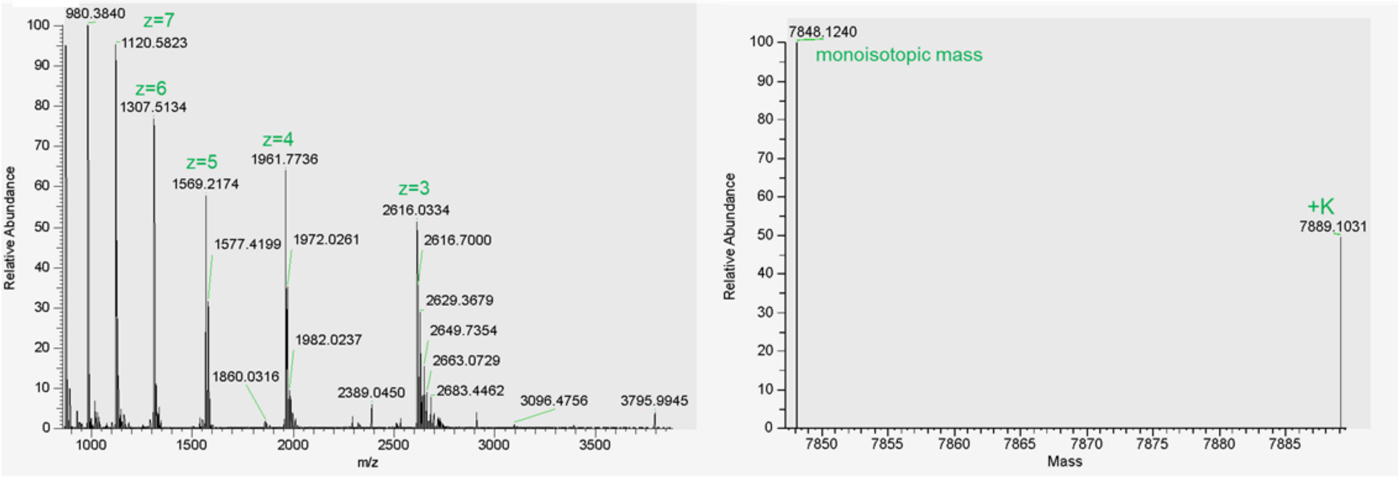


**i**

si<$(EG_{30}L)_2$

Theoretical molecular weight: 10,735 Da

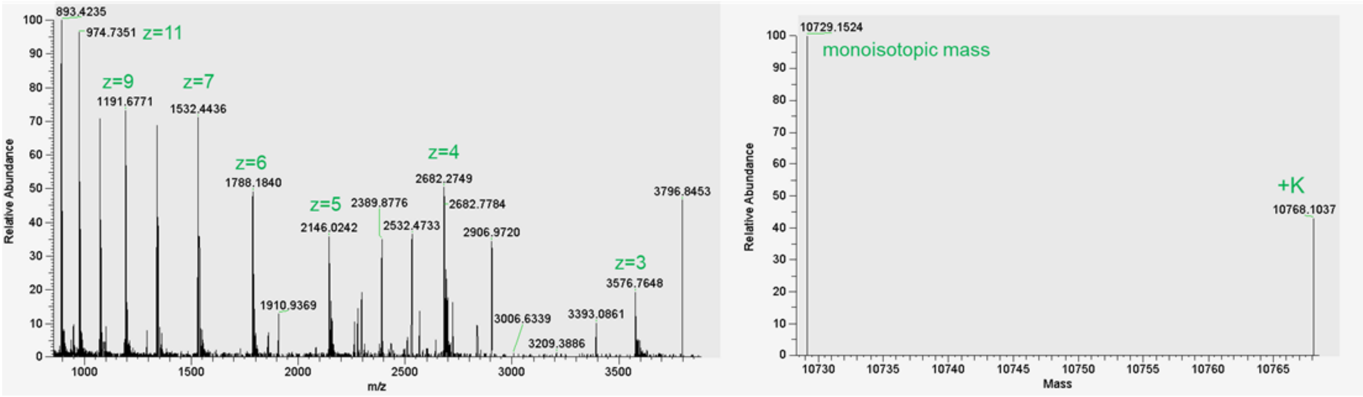

**j**

si<($EG_{18}L)_2$ No 5' PS or Binder PS

Theoretical molecular weight: 9115 Da

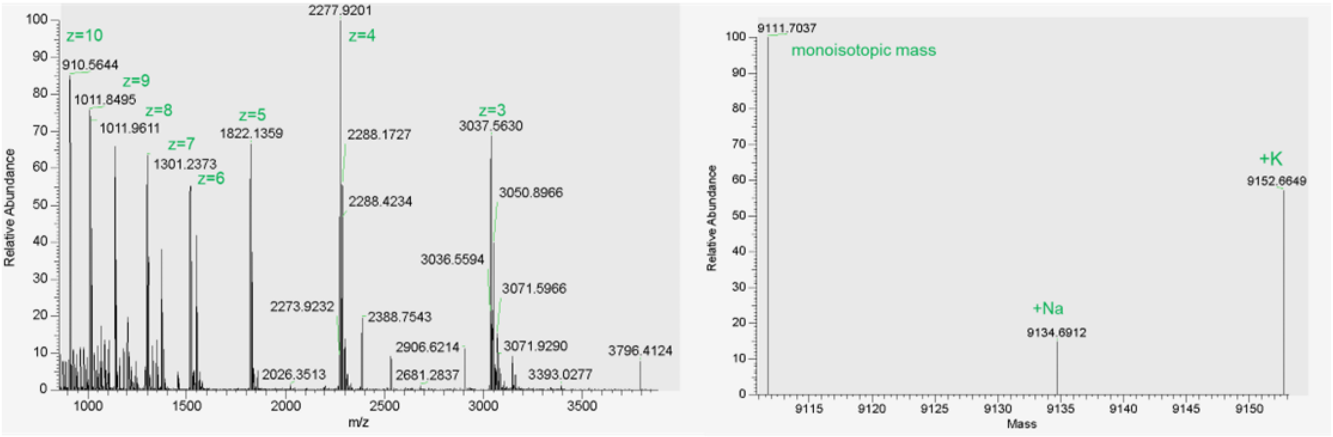


**k**

si<($EG_{18}L)_2$ No 5' PS

Theoretical molecular weight: 9260 Da

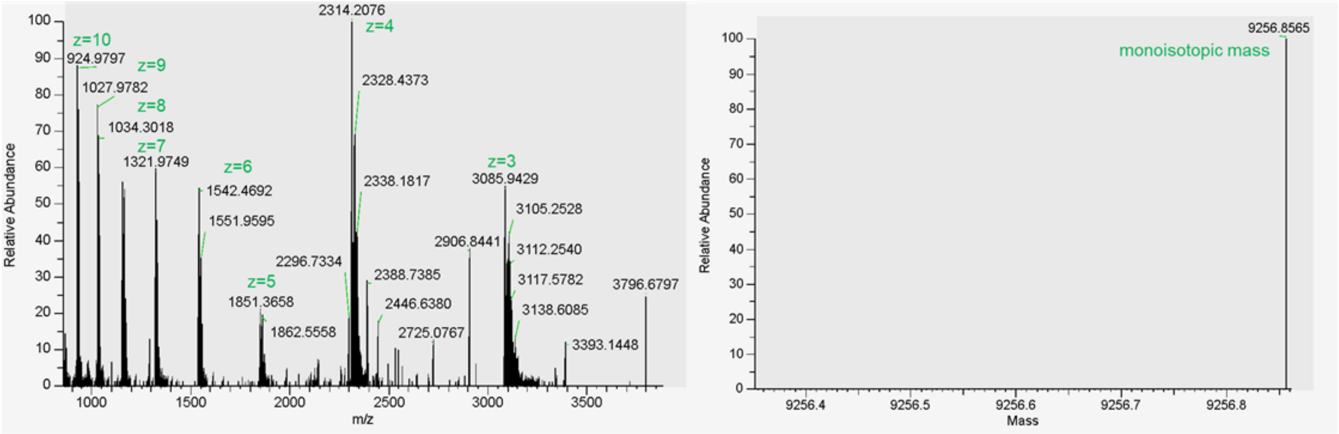


**l**

si-$EG_{18}$<$L_2$

Theoretical molecular weight: 8211 Da

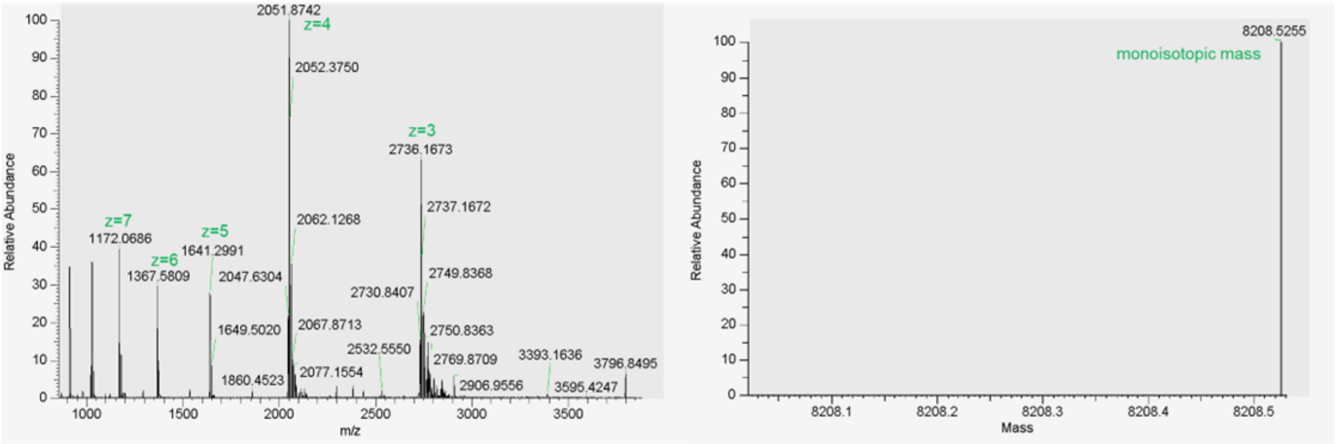

**m**

$\text{si-EG}_{36}<\text{L}_2$

Theoretical molecular weight: 9292 Da

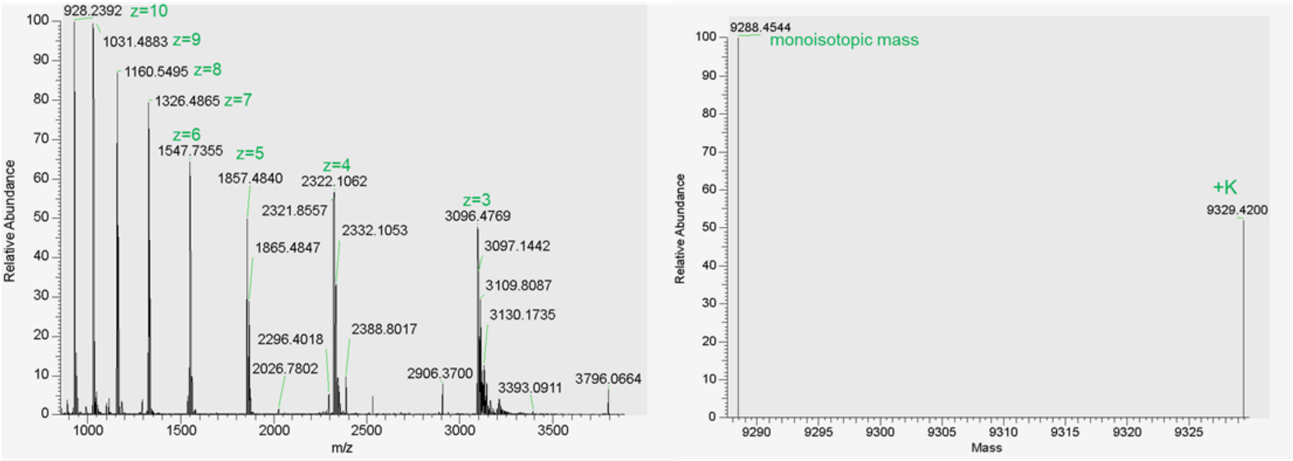


**n**

$\text{si}<(\text{EG}_{18}\text{L}_{\text{unsaturated}})_2$

Target molecular weight 9490

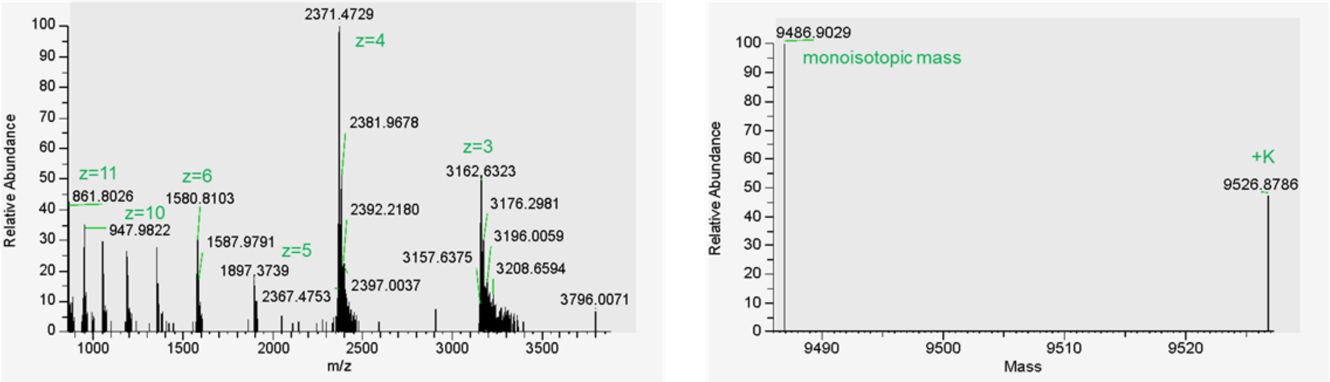


**o**

$\text{si}<(\text{EG}_{18}\text{L}_{\text{diacid}})_2$

Target molecular weight 9554

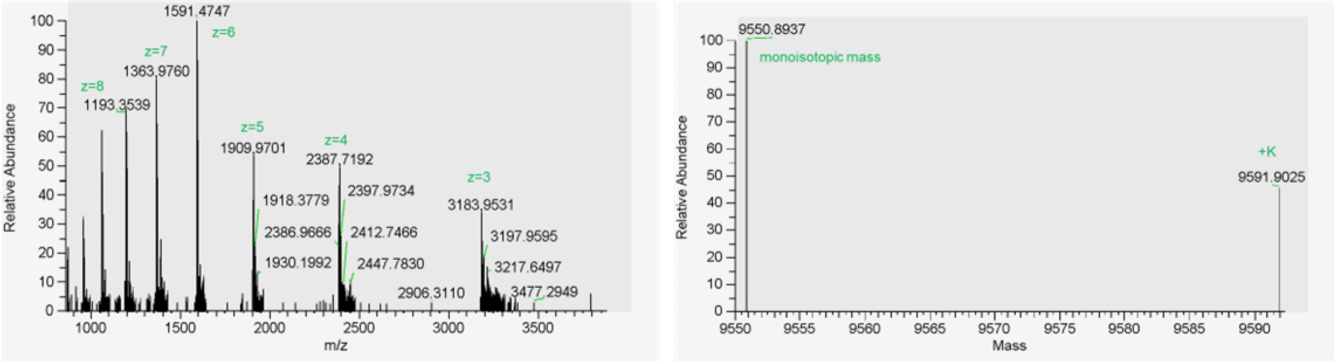

p

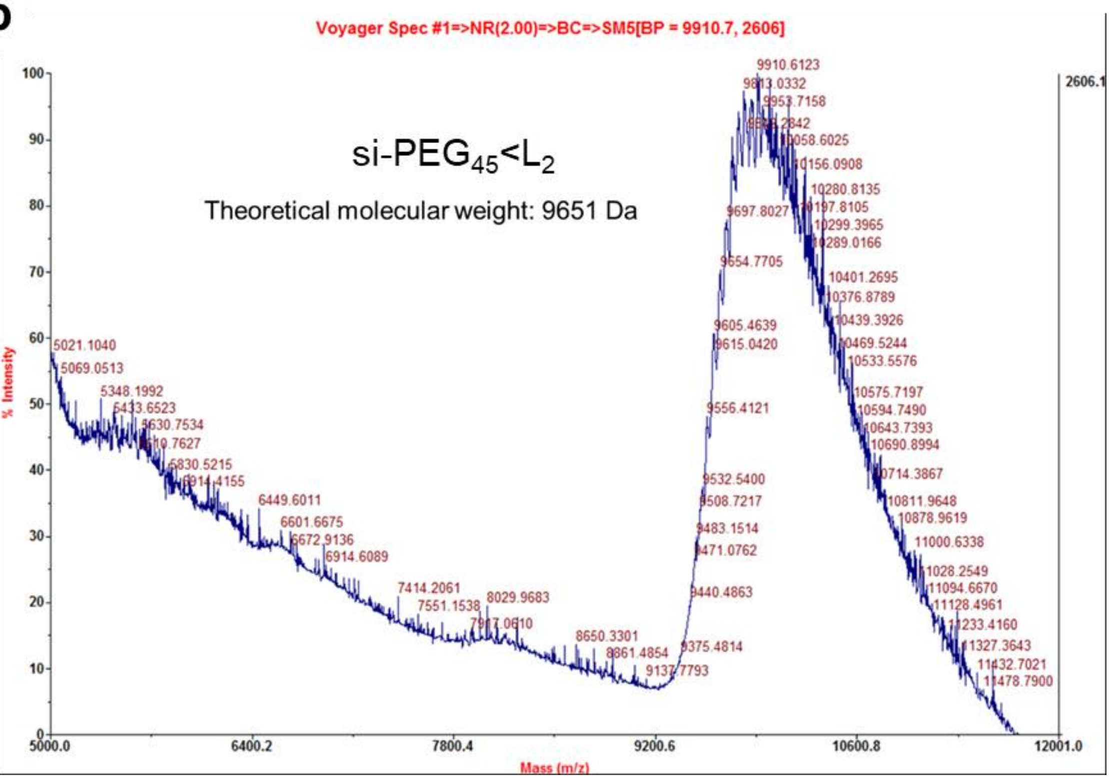

Voyager Spec #1=>NR(2.00)=>BC=>SM5[BP = 9910.7, 2606]
si-PEG$_{45}$<L$_2$
Theoretical molecular weight: 9651 Da
% Intensity
Mass (m/z)
2606.1
9910.6123

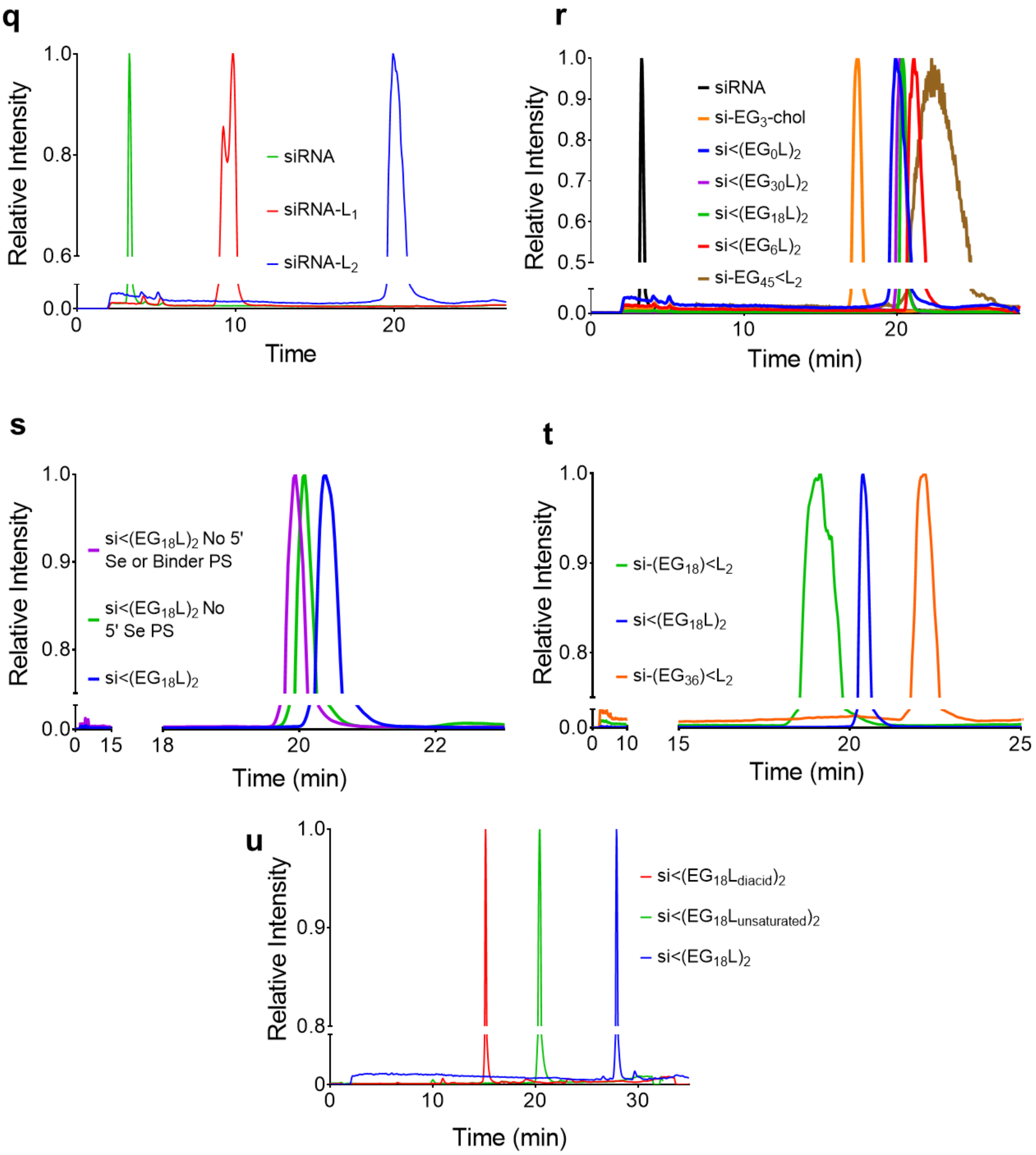


**Supplementary Figure 1: LC-MS characterization of siRNA and siRNA conjugates.** Conjugate purity and mass fidelity were confirmed by LCMS ESI$^-$. LC was performed using a Waters XBridge Oligonucleotide BEH C18 Column under a linear gradient from 85% A (16.3 mM triethylamine – 400 mM hexafluoroisopropanol) to 100% B (methanol) at 45°. (A-P) Full spectrum presented on left and deconvoluted mass presented on right as calculated by ThermoFisher FreeStyle Software. Molecular weight of si<PEG$_{45}$L$_2$ was validated using MALDI-TOF mass spectrometry. HLPC chromatograms for siRNAs with variations in valency (Q), number of ethylene glycol linker repeats (R), phosporothioate content (S), branching architecture (T), and lipid chemistry (U). Axes are adjusted to enable visualization of differences in elution time.

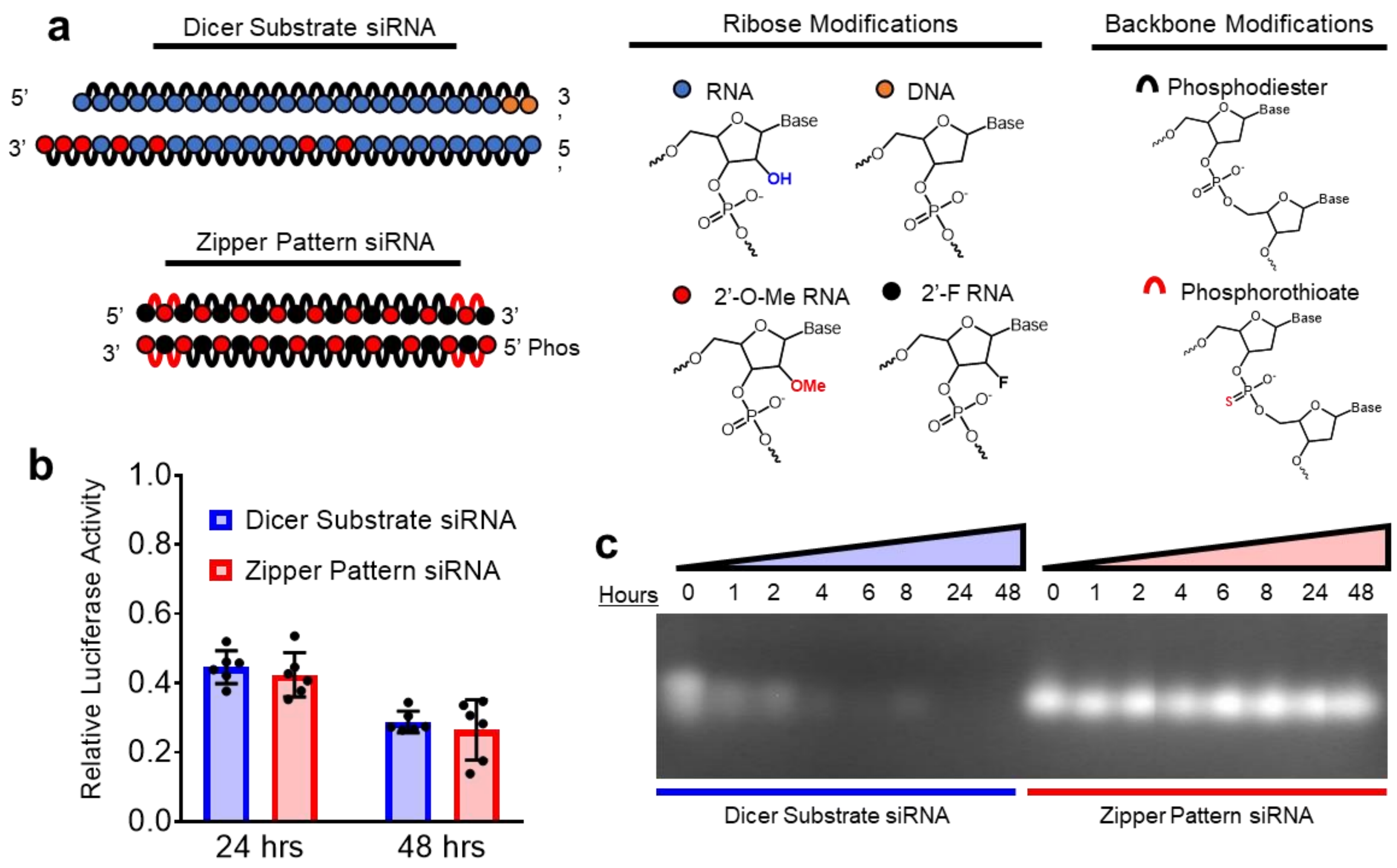


**Supplementary Figure 2**: **siRNA modification pattern validation and stability.** (A) Graphical schematic of structural differences between synthetic Dicer Substrate siRNA and alternating 2'F-, 2'OMe-modified "Zipper" siRNA used throughout this study. (B) Lipofection-mediated knockdown of 25 nM siRNA in Luciferase-expressing MDA-MB-231s. Luminescent signal was normalized to scrambled control siRNAs to account for any nonspecific toxicity effects. (C) Serum stability of siRNA challenged with 60% fetal bovine serum at 37°C visualized by agarose gel electrophoresis.

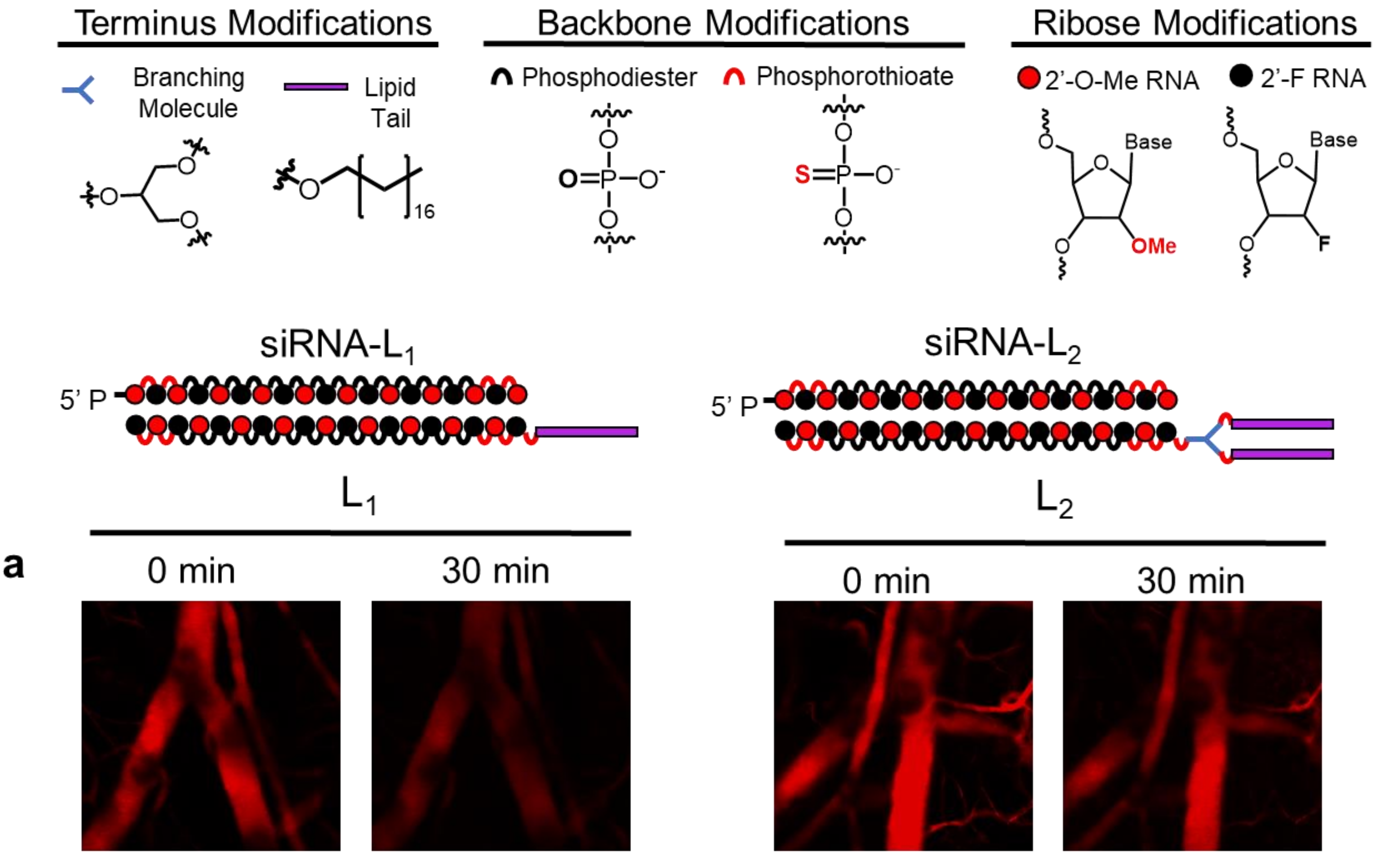


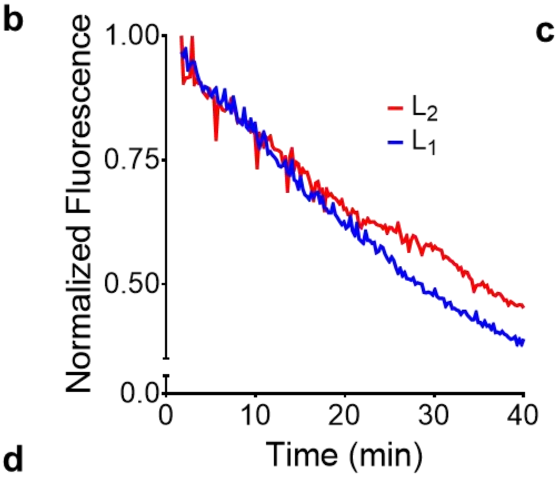


c

| | $t_{1/2}$ (min) | $AUC_{0\text{-}40m}$ (µg•min)/(mL) | CL (mL)/(min) |
|---|---|---|---|
| siRNA-$L_1$ | 28 ± 4.2 | 430 ± 56 | 0.026 ± 0.0058 |
| siRNA-$L_2$ | 46 ± 5.9** | 440 ± 51 | 0.018 ± 0.0040 |

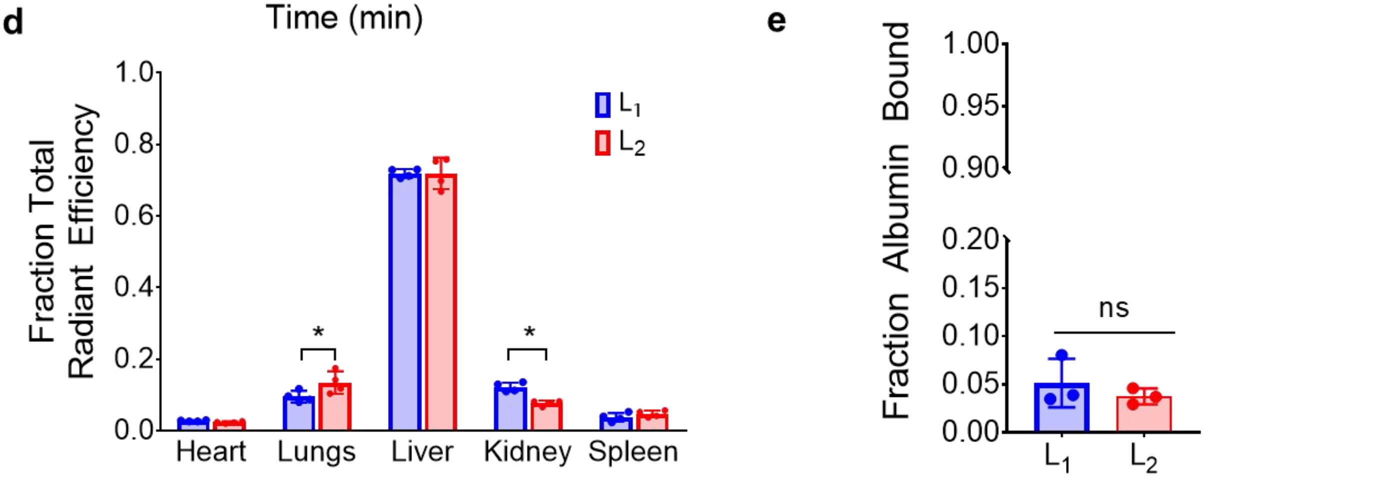


**Supplementary Figure 3: Divalent lipid-siRNA conjugate exhibits improved bioavailability compared to monovalent conjugate** A) Representative intravital microscopy images of mouse ear vasculature monitored for fluorescence decay after i.v. injection of siRNA conjugate. B) Average of (n=4) intravital fluorescent signals from individual mice displayed. C) Pharmacokinetic parameters of siRNA conjugates determined by intravital microscopy (D) Biodistribution of fluorescent conjugates approximately 45 minutes after 1 mg/kg intravenous injection as measured by IVIS (n=4). Significance assessed by 2-way ANOVA with Sidak's multiple comparisons tests (E) fraction of conjugate bound to albumin in mouse plasma after 1 mg/kg intravenous injection as calculated from known standards and sum of fractions' fluorescent intensities (n=3).Significance assessed using Welch's t-test.

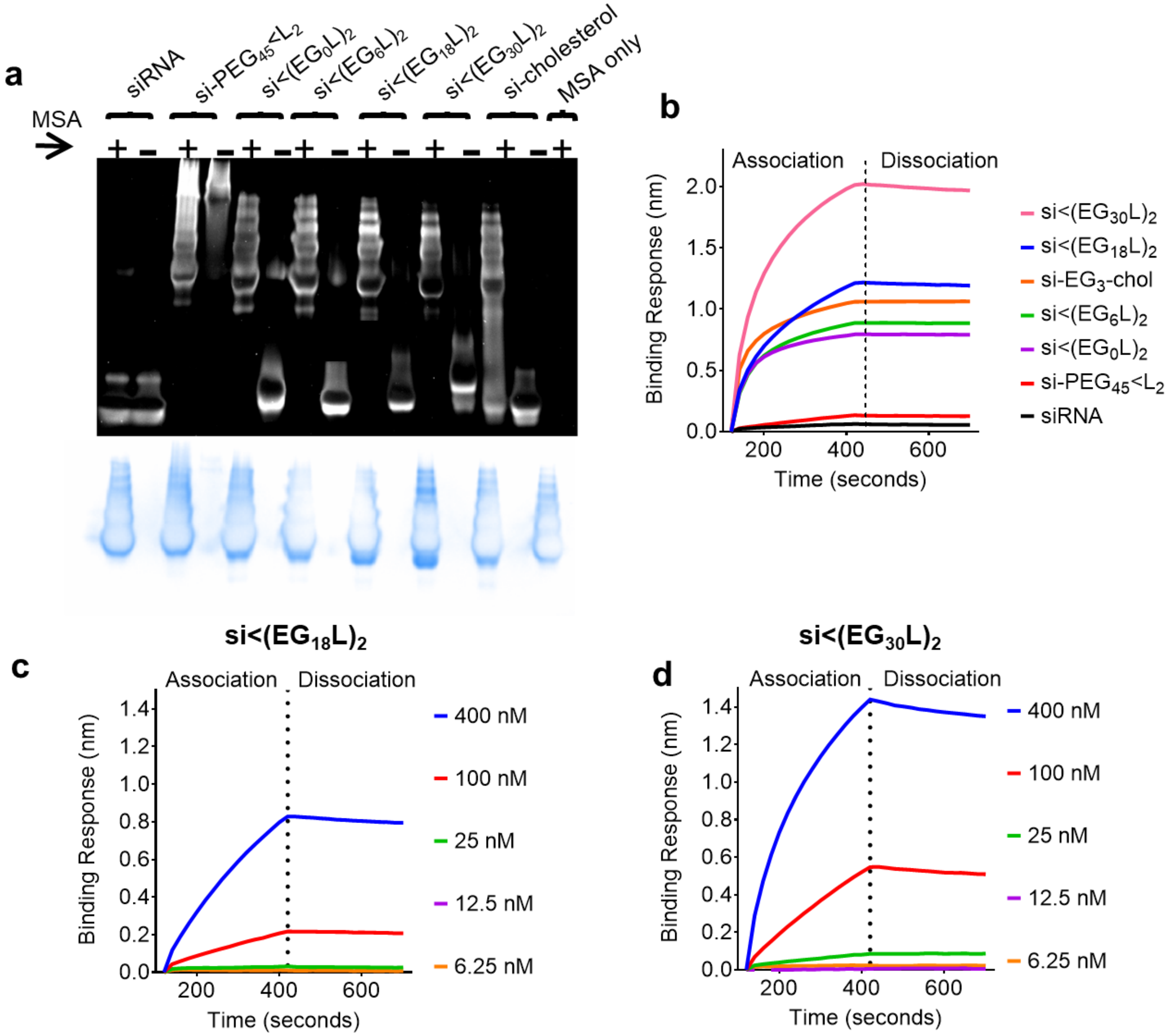


**e**

| | KD (nM) | $k_{on}$ (1/Ms) | $k_{off}$ (1/s) |
|---|---|---|---|
| **si<$(EG_{18}L)_2$** | 30 ± 0.3 | $6.97x10^3$ ±152 | $2.09x10^{-4}$ ± $1.75x10^{-6}$ |
| **si<$(EG_{30}L)_2$** | 9.49 ± 0.1 | $1.85x10^4$ ± 340 | $1.75x10^{-4}$ ± $2.09x10^{-6}$ |

**Supplementary Figure 4:** (A) Native PAGE Gel of siRNA conjugates run in the presence or absence of mouse serum albumin (MSA). Albumin colocalization can be seen by an upwards shift of nucleic acid staining to coincide with albumin (indicated in bottom panel by staining of the same gel with Coomassie blue). (B) Binding response of siRNA conjugates to 400 nM mouse serum albumin as measured by biolayer interferometry. Full panel of C) si<$(EG_{18}L)_2$ and D) si<$(EG_{36}L)_2$ binding responses to HSA used to determine E) binding kinetic parameters.

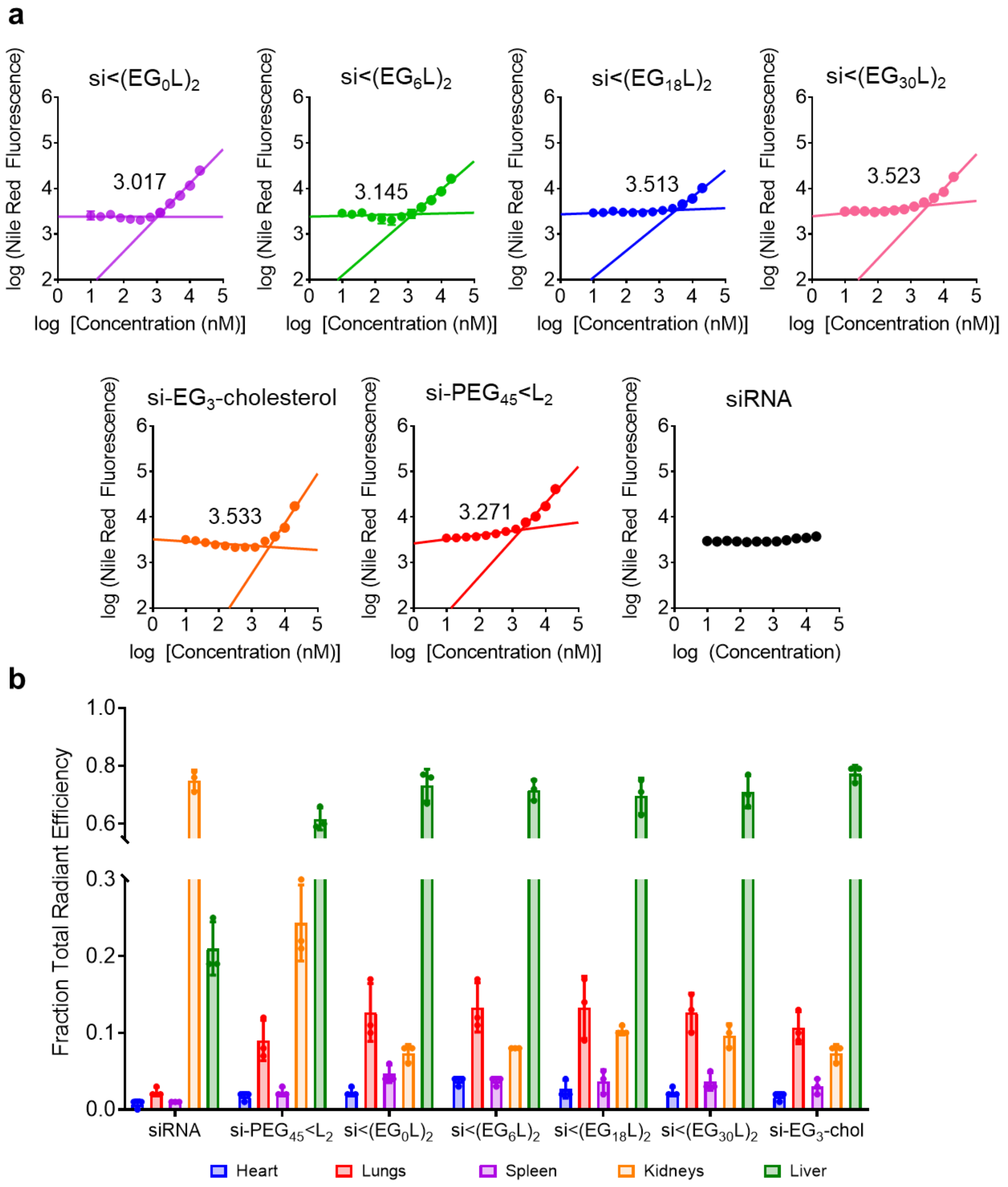


**Supplementary Figure 5**: (A) Critical micelle concentration (CMC) plots for siRNA conjugate library. (B) Biodistribution of Cy5-labeled siRNA conjugates in non-tumor bearing mice. Organs harvested approximately 1 h after injection at 1 mg/kg intravenously *via* tail vein (n=3).

| | | $t_{1/2}$ (min) | $AUC_{0\text{-}35m}$ (µg·min)/(mL) | CL (mL)/(min) |
|---|---|---|---|---|
| | siRNA | 14 ± 3.5 | 210 ± 25 | 0.069 ± 0.012 |
| | si-EG$_3$-cholesterol | 28 ± 12 | 410 ± 42 | 0.025 ± 0.0079 |
| | si-PEG$_{45}$<L$_2$ | 27 ± 5.3 | 390 ± 25 | 0.026 ± 0.0046 |
| EG Variants | si<(EG$_0$L)$_2$ | 33 ± 6.6 | 416 ± 55 | 0.028 ± 0.0059 |
| EG Variants | si<(EG$_6$L)$_2$ | 36 ± 4.6 | 450 ± 15 | 0.019 ± 0.0022 |
| EG Variants | si<(EG$_{18}$L)$_2$ | 64 ± 23 | 470 ± 52 | 0.012 ± 0.0032 |
| EG Variants | si<(EG$_{30}$L)$_2$ | 37 ± 6.6 | 440 ± 36 | 0.019 ± 0.0043 |
| PS Variants | si<(EG$_{18}$L)$_2$ No 5'Se PS | 37 ± 12 | 430 ± 50 | 0.020 ± 0.0060 |
| PS Variants | si<(EG$_{18}$L)$_2$ No 5'Se or Binder PS | 15 ± 1.5 | 300 ± 36 | 0.047 ± 0.0070 |
| Branching Variants | si-(EG$_{18}$)<L$_2$ | 67 ± 32 | 490 ± 35 | 0.012 ± 0.0070 |
| Branching Variants | si-(EG$_{36}$)<L$_2$ | 41 ± 13 | 450 ± 27 | 0.018 ± 0.0050 |
| Lipid Variants | si<(EG$_{18}$L$_{diacid}$)$_2$ | 47 ± 21 | 469 ± 64 | 0.017 ± 0.0073 |
| Lipid Variants | si<(EG$_{18}$L$_{unsaturated}$)$_2$ | 34 ± 26 | 391 ± 121 | 0.029 ± 0.016 |

**Supplementary Table 2: Pharmacokinetic parameters for full siRNA conjugate library determined from intravital microscopy**

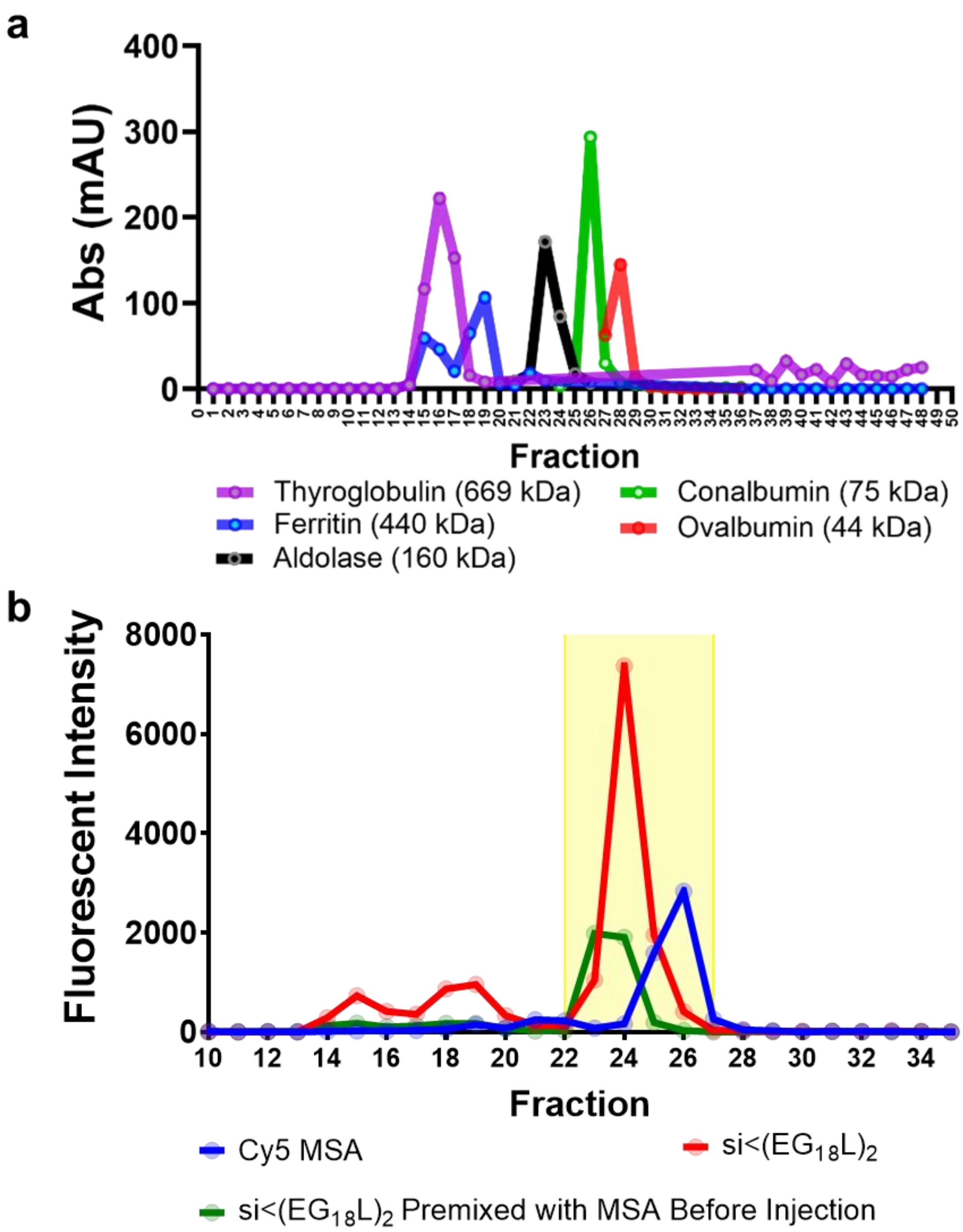


**Supplementary Figure 6:** (A) Size exclusion chromatography (SEC) of protein standards. Representative protein standards run on SEC system to demonstrate resolution by molecular weight. (B) Size exclusion chromatography (SEC) of mouse plasma after injection with siRNA conjugate +/- MSA versus MSA alone. Mice were injected at 1 mg/kg intravenously via tail vein and plasma was harvested approximately 1h after injection**.** Fractions highlighted in yellow were used for calculating albumin-bound fraction.

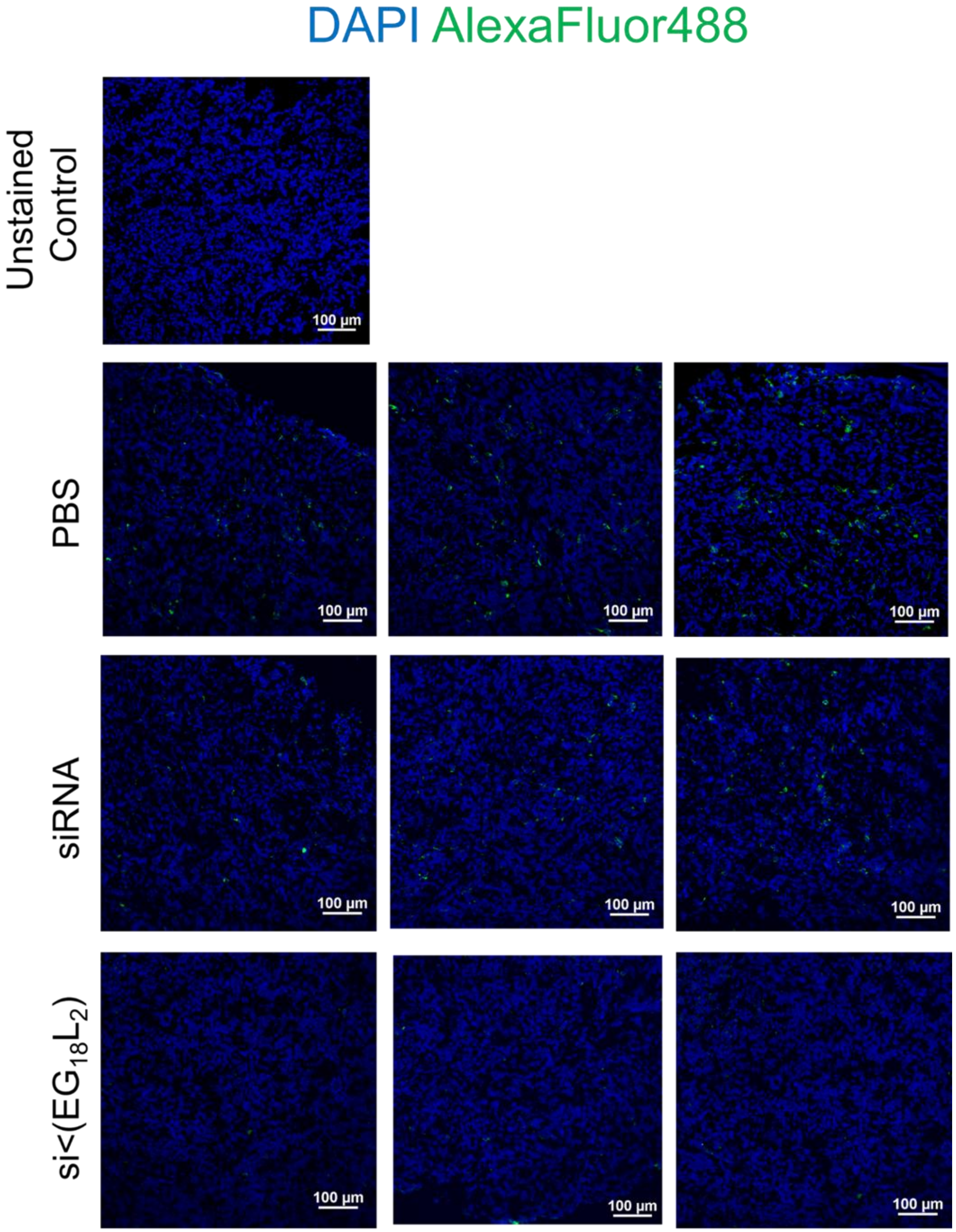


**Supplementary Figure 7: Extended images of tumors sectioned and immunohistochemically stained for Firefly Luciferase protein (green, Alexafluor488) and cell nuclei (blue, DAPI).**

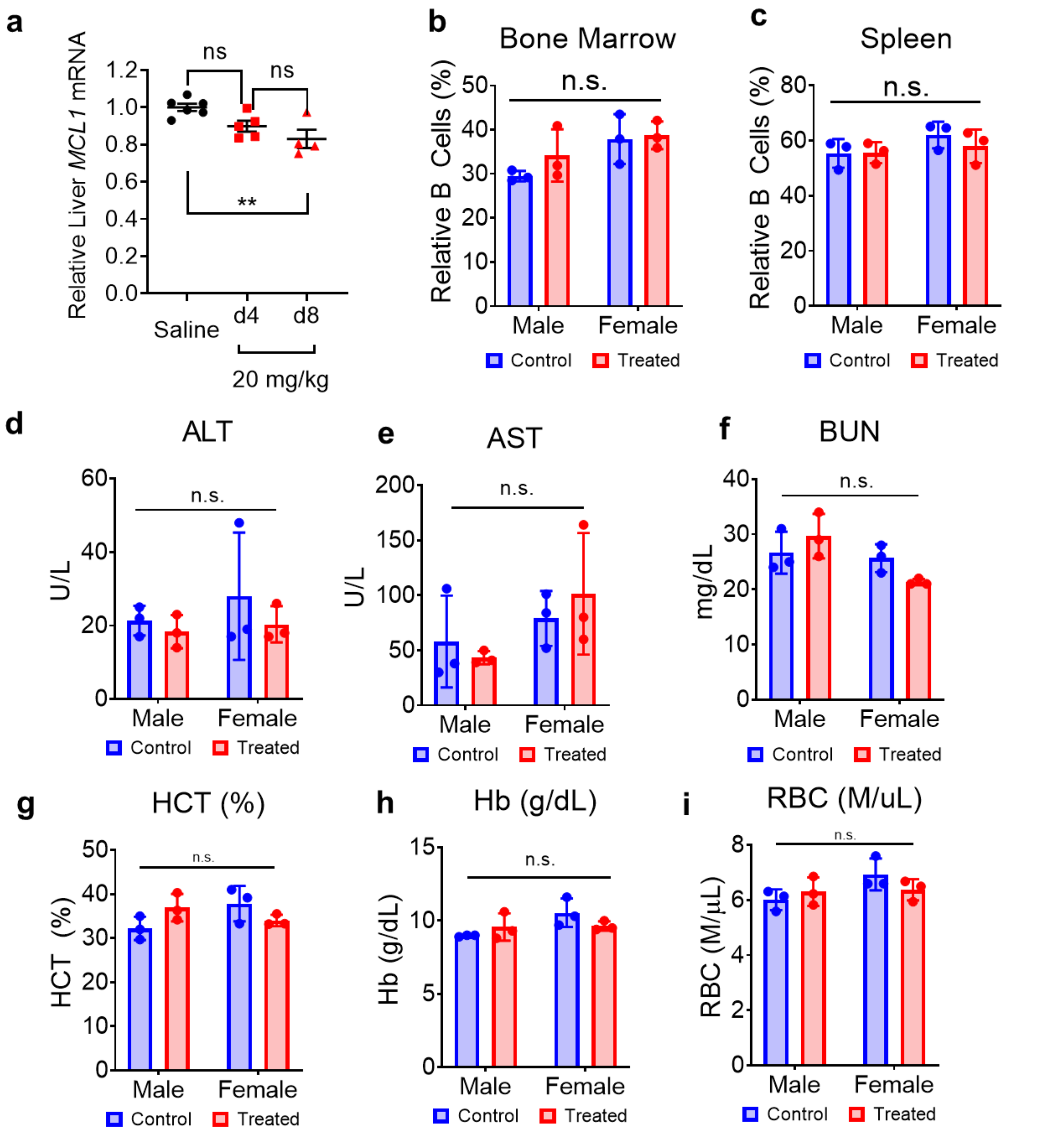


**Supplementary Figure 8:** (A) Liver knockdown of Mcl-1 mRNA after bolus i.v. injection of si$_{Mcl-1}$<(EG$_{18}$L)$_{2}$. Significance assessed by 1-way ANOVA with Tukey's multiple comparisons test (n=4-6). B cells relative to total viable cells isolated from (B) bone marrow and (C) spleen isolated from wild type mice three days after injection with 20 mg/kg siRNA conjugate targeting Mcl-1 compared to untreated controls. Levels of blood chemistry markers (D) alanine aminotransferase (E) aspartate aminotransferase and (F) blood urea nitrogen from sera of treated and untreated mice. Significance assessed by 2-way ANOVA with Tukey's multiple comparisons test (n=3). (G-I) Complete blood count values collected from mice 3 days after injection with 20 mg/kg of siRNA conjugate targeting Mcl-1. Significance assessed by 2-way ANOVA (n=3).

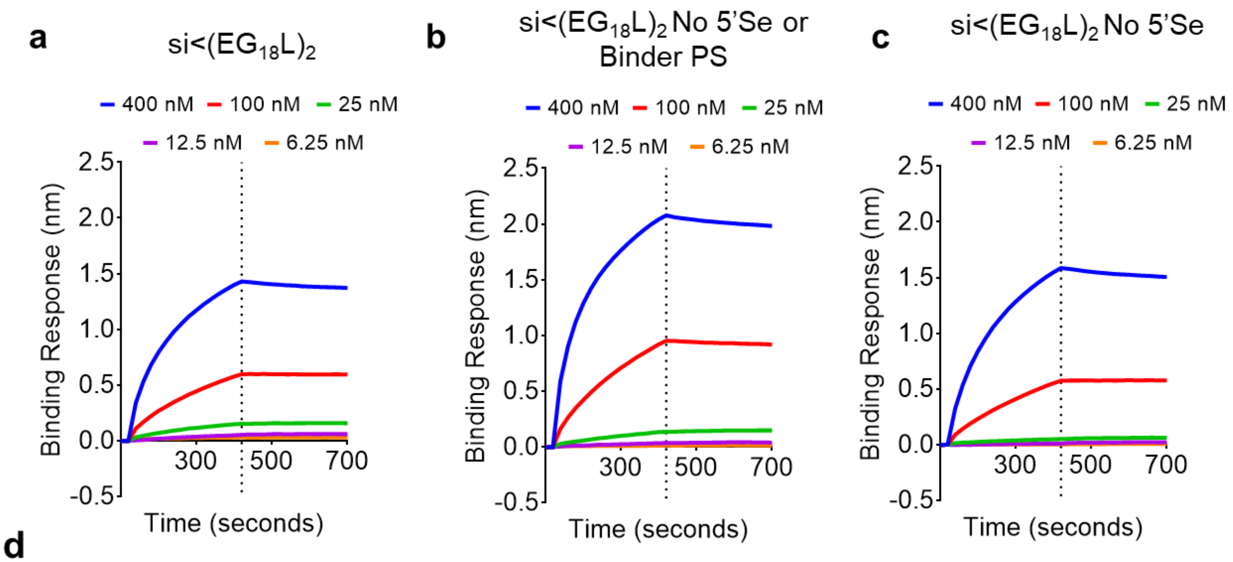


d

| | KD (nM) | $k_{on}$ (1/Ms) | $k_{off}$ (1/s) |
|---|---|---|---|
| **si<($EG_{18}L$)$_2$** | 3.89 ± 0.13 | $2.35x10^4$ ± 6.37 | $9.14x10^{-5}$ ± $3.09x10^{-6}$ |
| **si<($EG_{18}L$)$_2$ No 5'Se or Binder PS** | 2.81 ± 0.10 | $3.01x10^4$ ± 7.47 | $8.46x10^{-5}$ ± $3.05x10^{-6}$ |
| **si<($EG_{18}L$)$_2$ No 5'Se** | 5.58 ± 0.10 | $2.08x10^4$ ± 3.91 | $1.16x10^{-4}$ ± $2.07x10^{-6}$ |

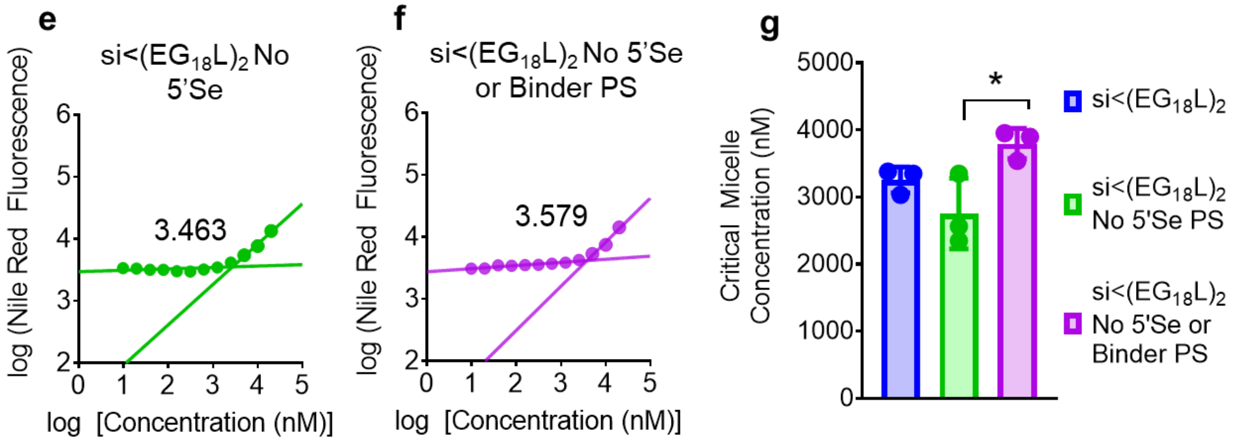


**Supplementary Figure 9: Extended characterization of si<($EG_{18}L$)$_2$ phosphorothioate variants.** (A-C) Full panel of binding responses to HSA at varied concentrations measured by biolayer interferometry used to determine D) binding kinetic parameters (E-G) Critical micelle concentration observed after 2-hour incubation with Nile Red at 37°C (n=3). Significance assessed by 1-way ANOVA with Tukey's multiple comparisons test.

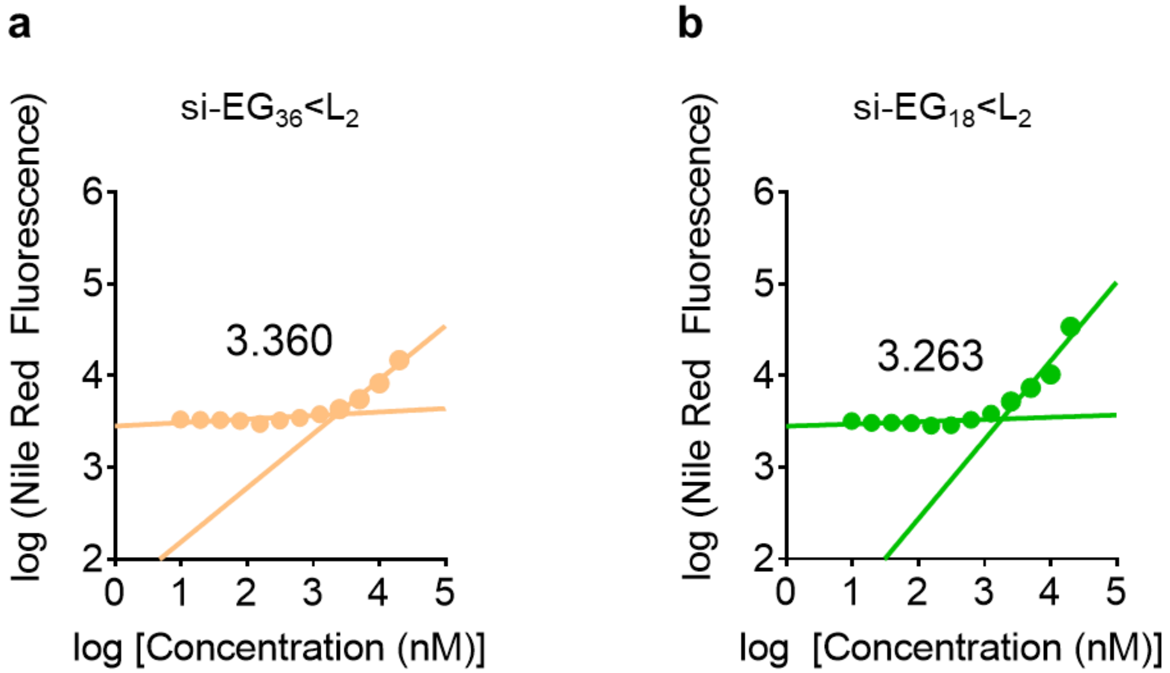


**Supplementary Figure 10:** Critical micelle concentration (CMC) plots for (A) si-$EG_{36}$<$L_2$ and (B) si-$EG_{18}$<$L_2$

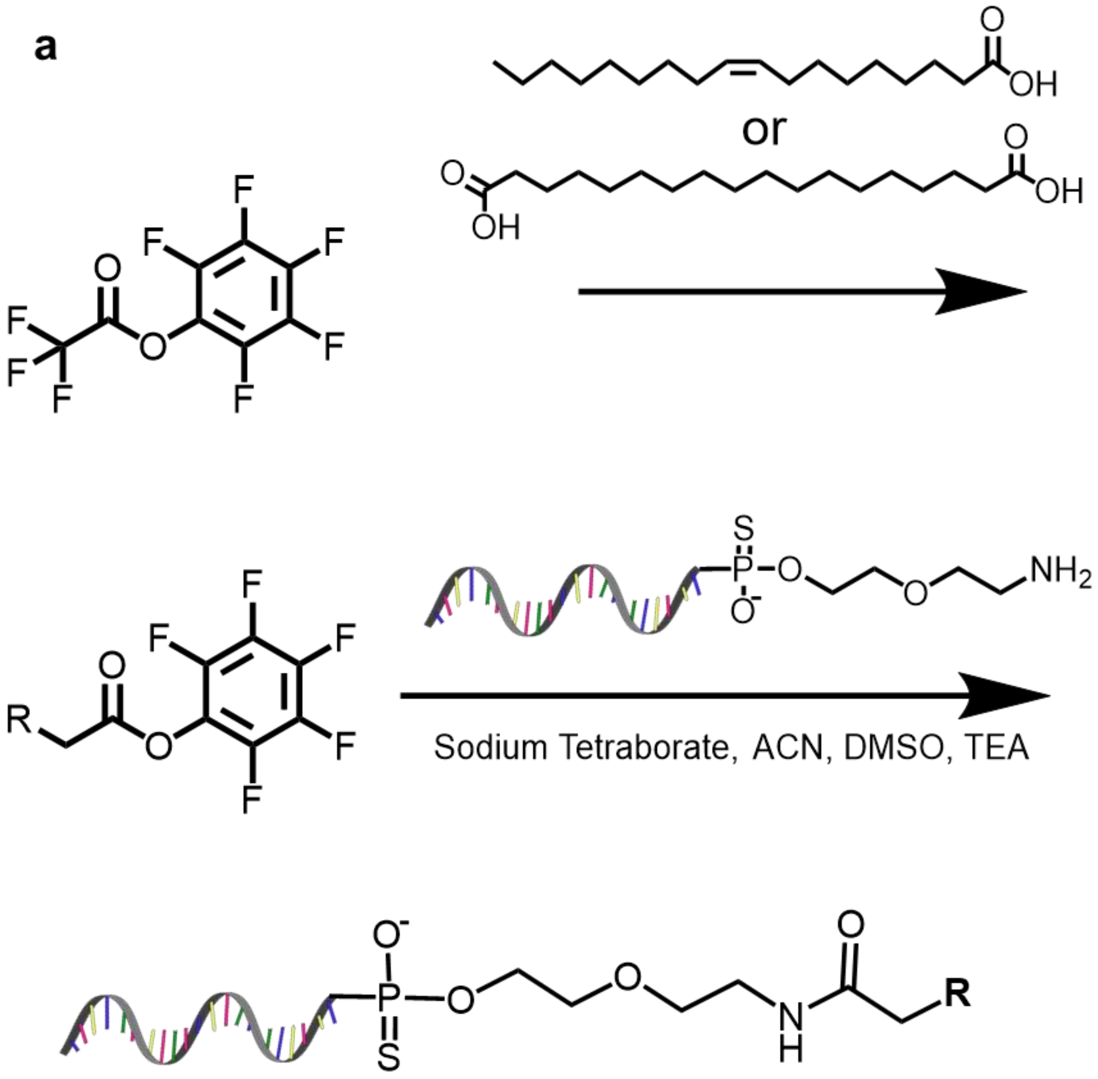


Oleic acid was dissolved at 1mM in anhydrous dichloromethane and placed on ice. A 10x molar excess of triethylamine was then added followed by a 3x molar excess of pentafluorophenyl trifluoroacetate. A similar procedure was used for octadecanoic acid. The lipid was dissolved at 0.5 mM in anhydrous dichloromethane and placed on ice. A 2x molar excess of triethylamine was then added followed by a 0.5x molar excess of pentafluorophenyl trifluoroacetate added dropwise.

For both lipids, the reaction vessel was removed from ice after 15 minutes and allowed to equilibrate to room temperature. The reaction mixture was then stirred for 4h followed by storage at -20°C. The crude product was purified by silica gel column chromatography and eluted with solvents with a gradient from 100% hexane to 100% ethyl acetate to yield the purified product. The compounds were then rotovapped to dryness and characterized by H and F NMR.

b

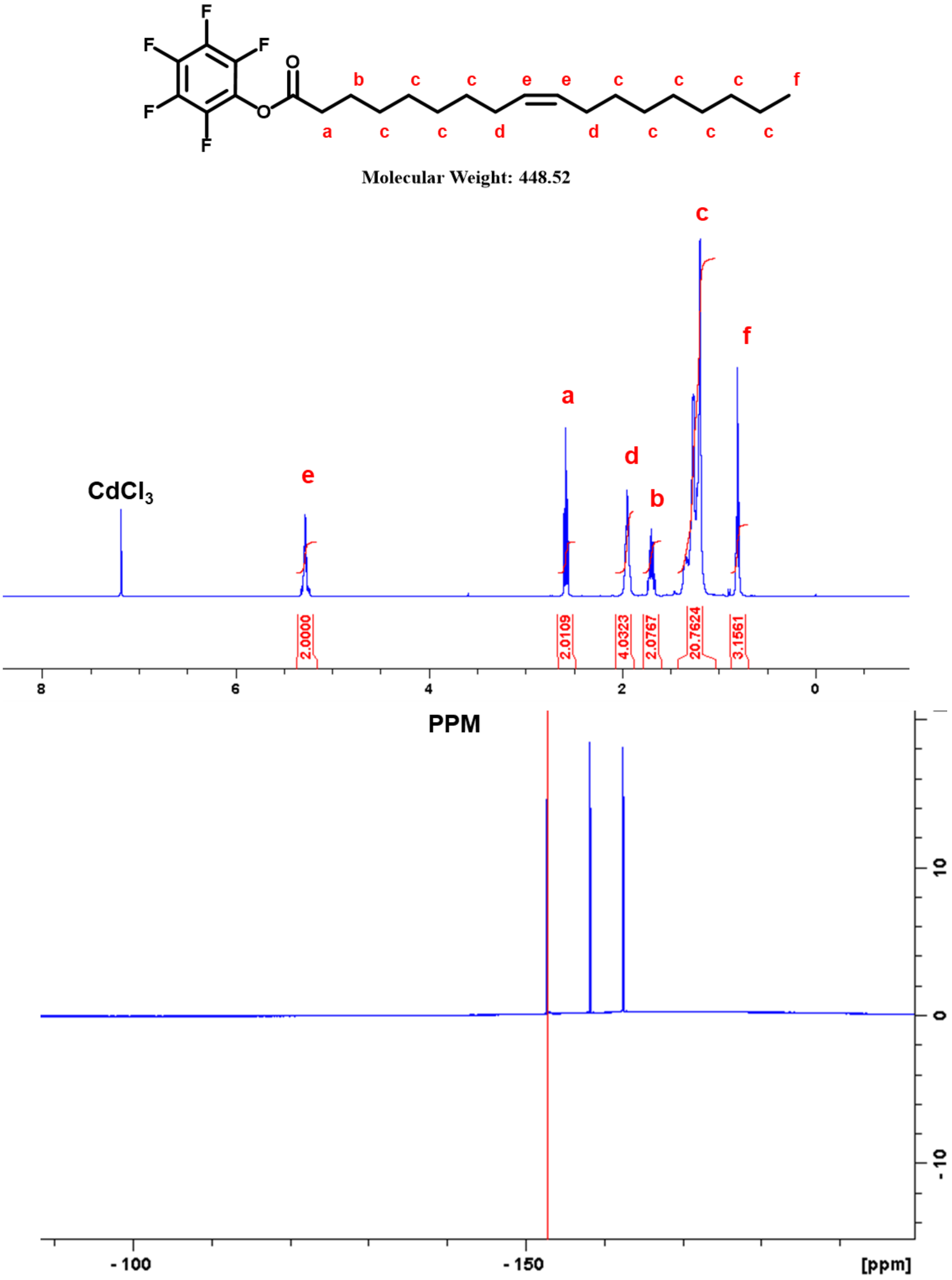

Molecular Weight: 448.52
CdCl$_3$
e
a
d
b
c
f
2.0000
2.0109
4.0323
2.0767
20.7624
3.1561
8
6
4
2
0
PPM
10
0
-10
- 100
- 150
[ppm]

c

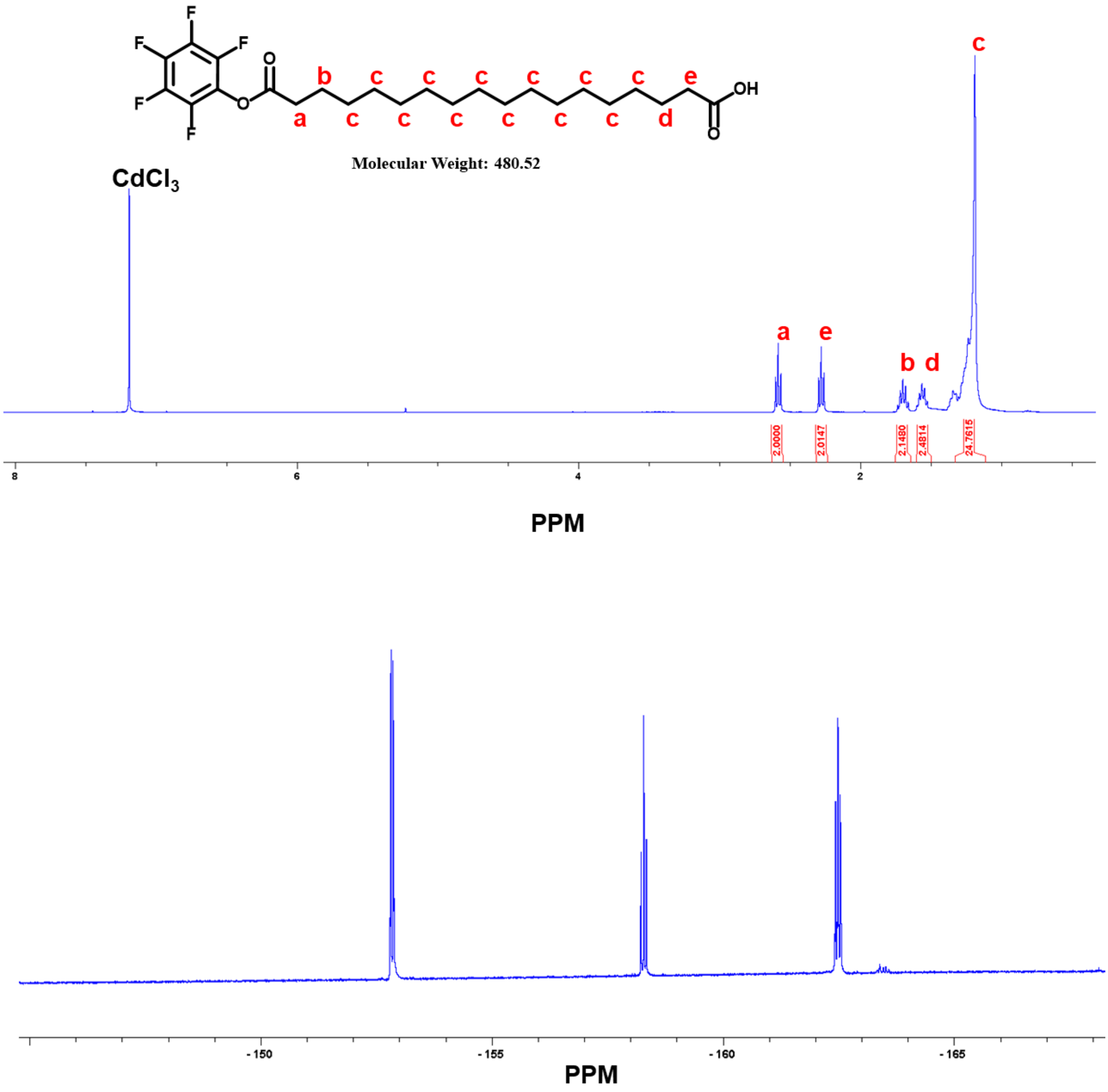


**Supplementary Figure 11: Synthesis of si<(EG$_{18}$L)$_2$ lipid variants**

(A) Overview of PFPA mediated lipid conjugation with (B) unsaturated lipid and (C) diacid lipid characterization by $^{1}$H NMR (top panel) and $^{19}$F NMR (bottom panel)

### Synthesis of lipid-variant siRNA conjugates

Amine-terminated oligonucleotides were speed vacuumed to dryness and desalted to remove MMT groups. Oligonucleotides were then lyophilized followed by reconstitution in 0.1 sodium tetraborate (pH 8.5) to a concentration of 500 uM. PFP-modified lipid was dissolved into a mixture of acetonitrile, DMSO, and triethylamine (70:29:1 by volume) at a concentration of 7 uM. Aqueous oligonucleotide was added dropwise to the organic solution for a 1:40 molar ratio of oligonucleotide-amine:amine-reactive lipid (approximately 25% 0.1M sodium tetraborate, 75% organic mixture). Solution was stirred overnight and desalted prior to purification and characterization detailed in main methods section.

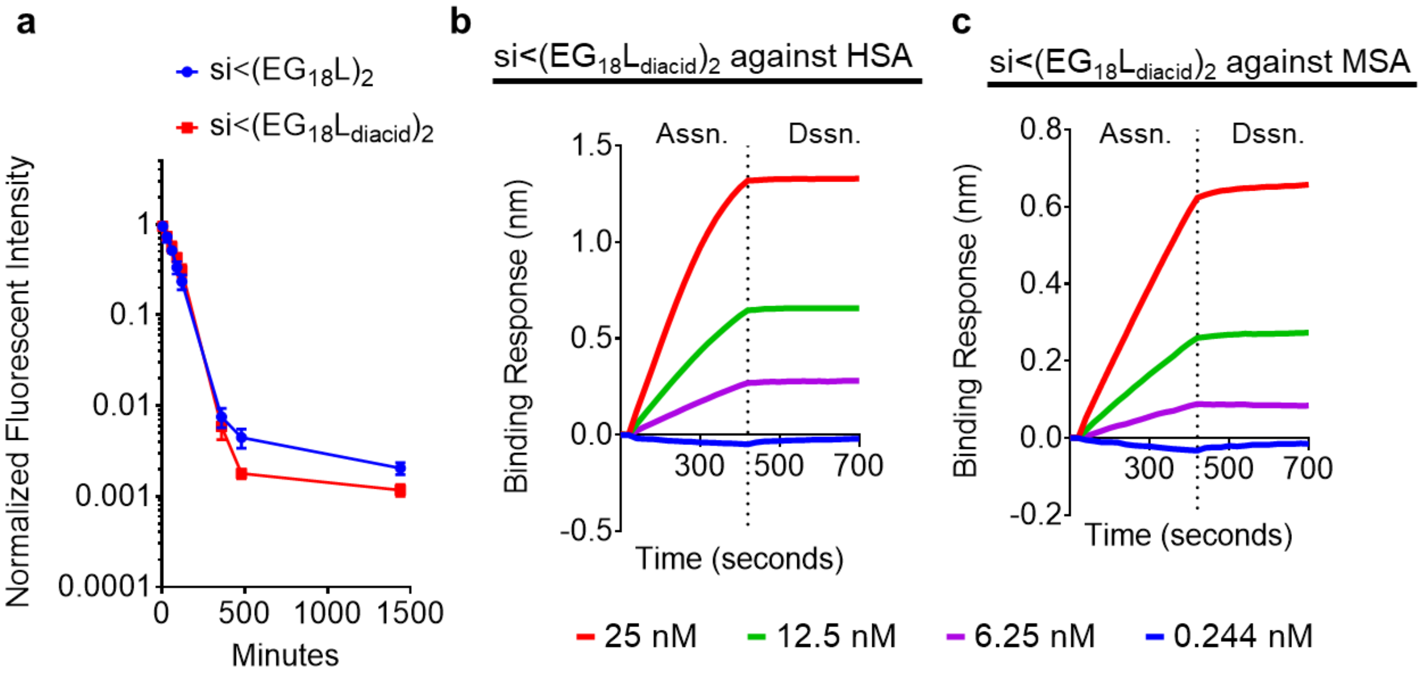


**d** HSA Binding Kinetics

| | **KD (nM)** | **ka (1/Ms)** | **kdis (1/s)** |
|---|---|---|---|
| **si<(EG$_{18}$L$_{diacid}$)$_2$** | 0.15 ± 0.002 | $1.31\times10^5$ ± 712 | $1.93\times10^{-5}$ ± $2.92\times10^{-6}$ |
| **si<(EG$_{18}$L)$_2$** | 30 ± 0.3 | $6.97\times10^3$ ±152 | $2.09\times10^{-4}$ ± $1.75\times10^{-6}$ |

**Supplementary Figure 12: Extended characterization of diacid-modified si<(EG$_{18}$L)$_2$.**

(A) Mice were injected intravenously with 5 mg/kg of Cy5-labeled siRNA conjugate (n=3-4). At various timepoints, blood was sampled from the opposite vein from injection and fluorescence was measured to determine PK profiles at later timepoints. (B-C) Binding responses to HSA and MSA at varied concentrations measured by biolayer interferometry used to determine D) binding kinetic parameters

### Synthesis of siRNA conjugate variant covalently bound to albumin

Conjugate covalently bound to albumin was synthesized by first reacting azido-PEG$_3$-maleimide (Click Chemistry Tools) with the free thiols on human (1 free SH) or mouse (2 free SH). Albumin was dissolved in

PBS with 0.5M EDTA to a final concentration of 10 mM. Anhydrous DMF was used to solubilize and activate azido-PEG$_3$-maleimide. DBCO-modified siRNA duplex in PBS was reacted at a 1:1 ratio of DBCO groups:free SH groups and allowed to incubate at room temperature for 4 hours. To remove any siRNA that did not react with albumin, or reacted only with the azido linker, the resulting solution underwent 10 rounds of centrifugation in a 30 kDa cutoff Amicon filter at 14,000 xg for 10 minutes for each round. Conjugation was confirmed by gel electrophoresis of precusor DBCO-siRNA alongside resulting siRNA-DBCO-albumin.

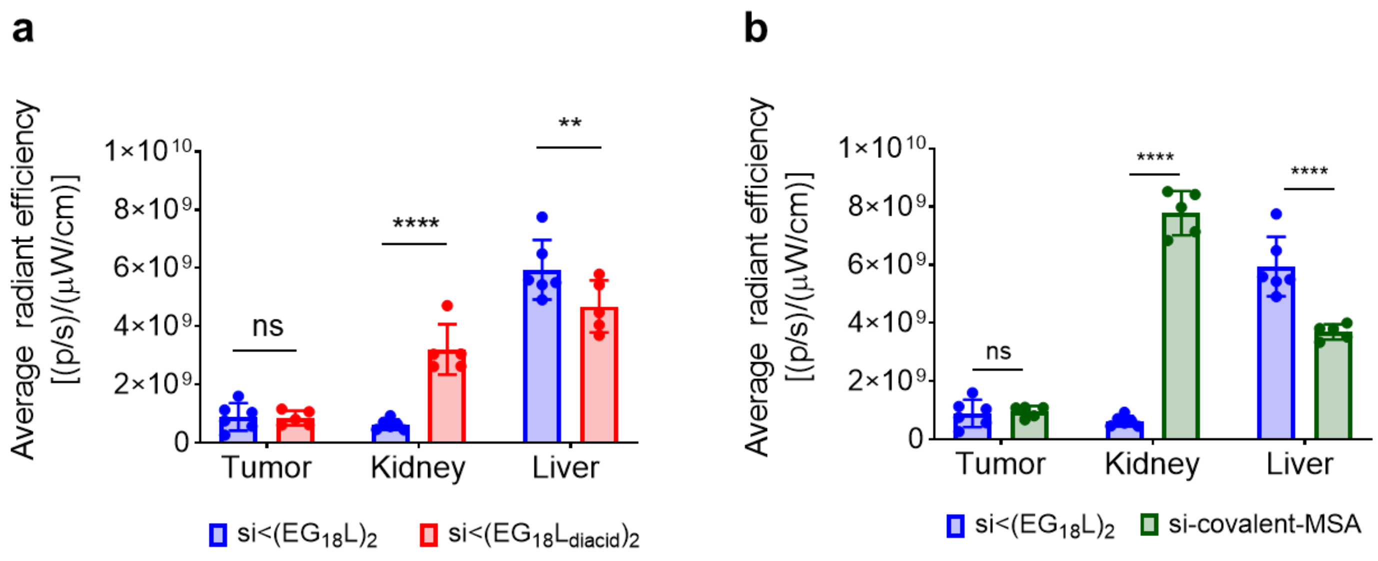


**Supplementary Figure 13: Biodistribution of diacid and covalent siRNA conjugates in tumor bearing mice.** Epifluorescence of whole organs measured by IVIS comparing accumulation of fluorescently labeled A) si<(EG$_{18}$L$_{diacid}$)$_2$ and B) si-covalent-MSA to lead conjugate si<(EG$_{18}$L)$_2$ 18h after i.v. injection of 1 mg/kg siRNA

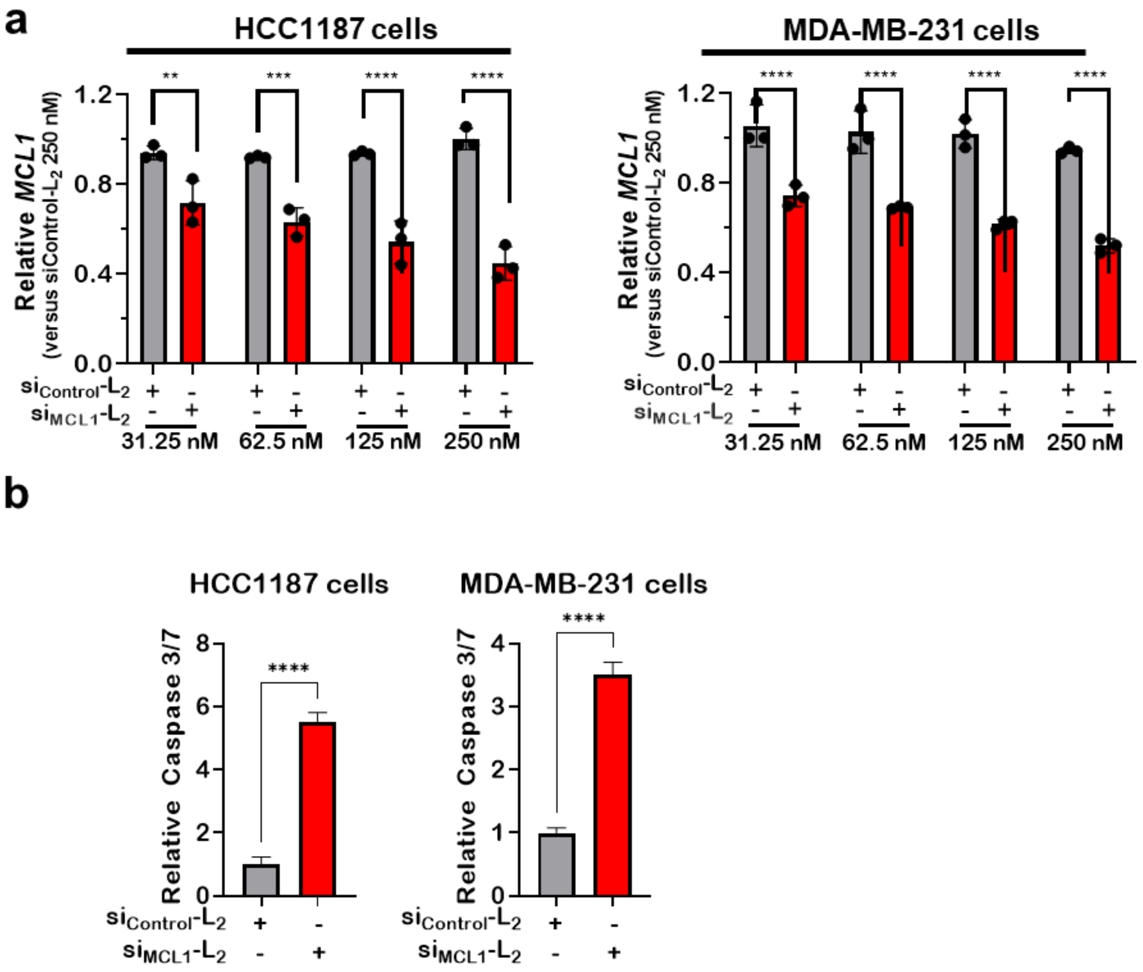


**Supplementary Figure 14:** HCC1187 and MDA-MB-231 cells were treated in serum-free OptiMEM with siRNA-lipid conjugates and assessed 48 hours post-treatment for expression of MCL1 transcript levels, corrected for expression of the housekeeping gene PPIB, and are shown relative to *MCL1* levels seen in $si_{Control}$-$L_2$ (250 nM)-treated cells. **B.** HCC1187 and MDA-MB-231 cells treated with siRNA-lipid conjugates for 96 hours at 250 nM were assessed using Caspase 3/7-Glo.

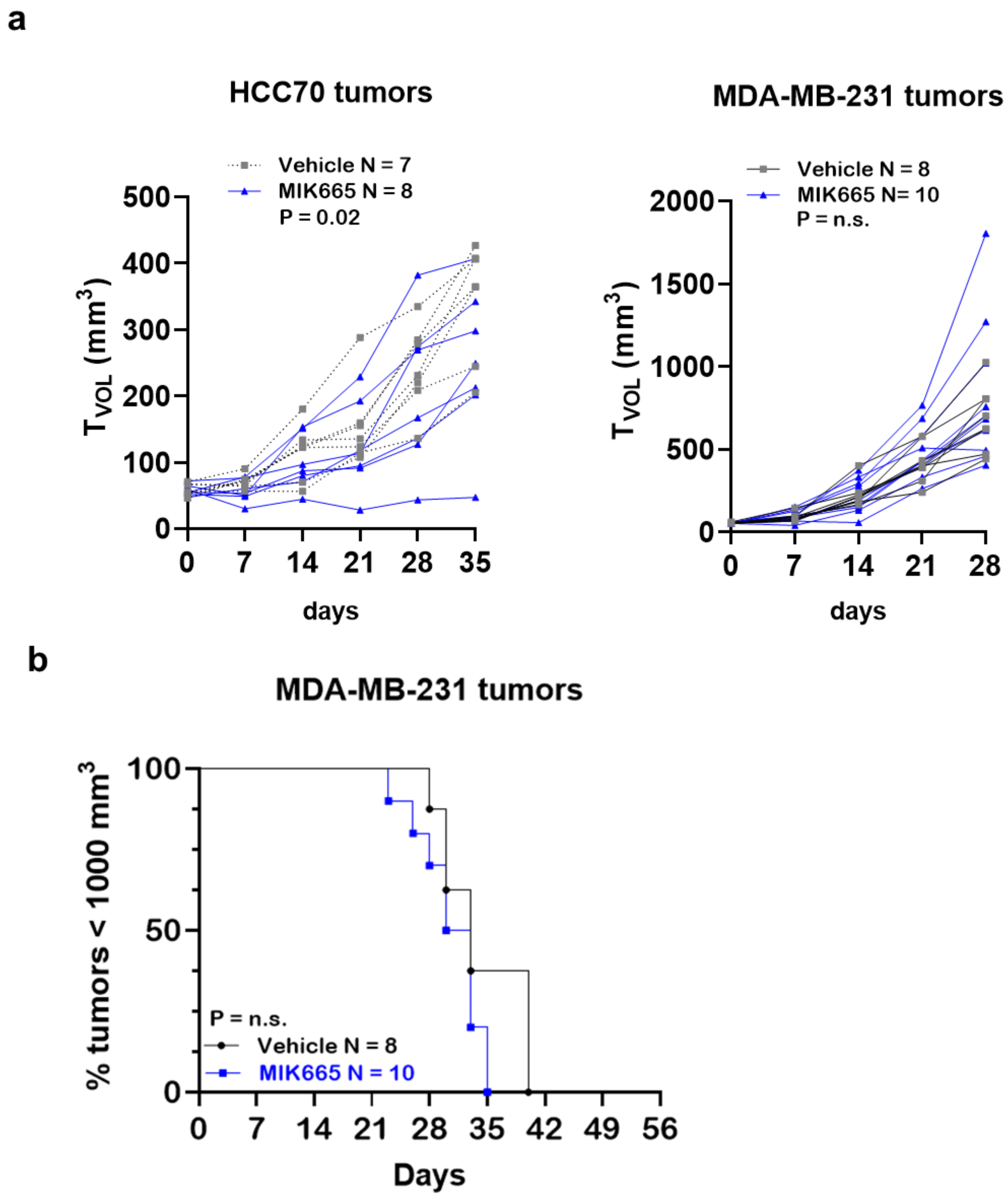


**Supplementary Figure 15**: Mice harboring HCC70 (N = 7-8) and MDA-MB-231 (N = 8-10) orthotopic tumors (50 -100 mm$^3$) received once weekly doses of vehicle (used for delivery of MIK665), or MIK665 through day 28 by i.v. injection. Tumors were collected on day 35 (HCC70) or day 56 (MDA-MB-231) or earlier if tumors ulcerated or exceeded humane size limitations. **A.** Tumor volumes were measured throughout treatment. Statistical values were calculated based on area under the curve. **B.** Kaplan-Meier analysis of tumor bearing mice, defining survival as tumor volume under 1000 mm$^3$. P values are calculated using the log-rank (Mantel-Cox) test.

**a**

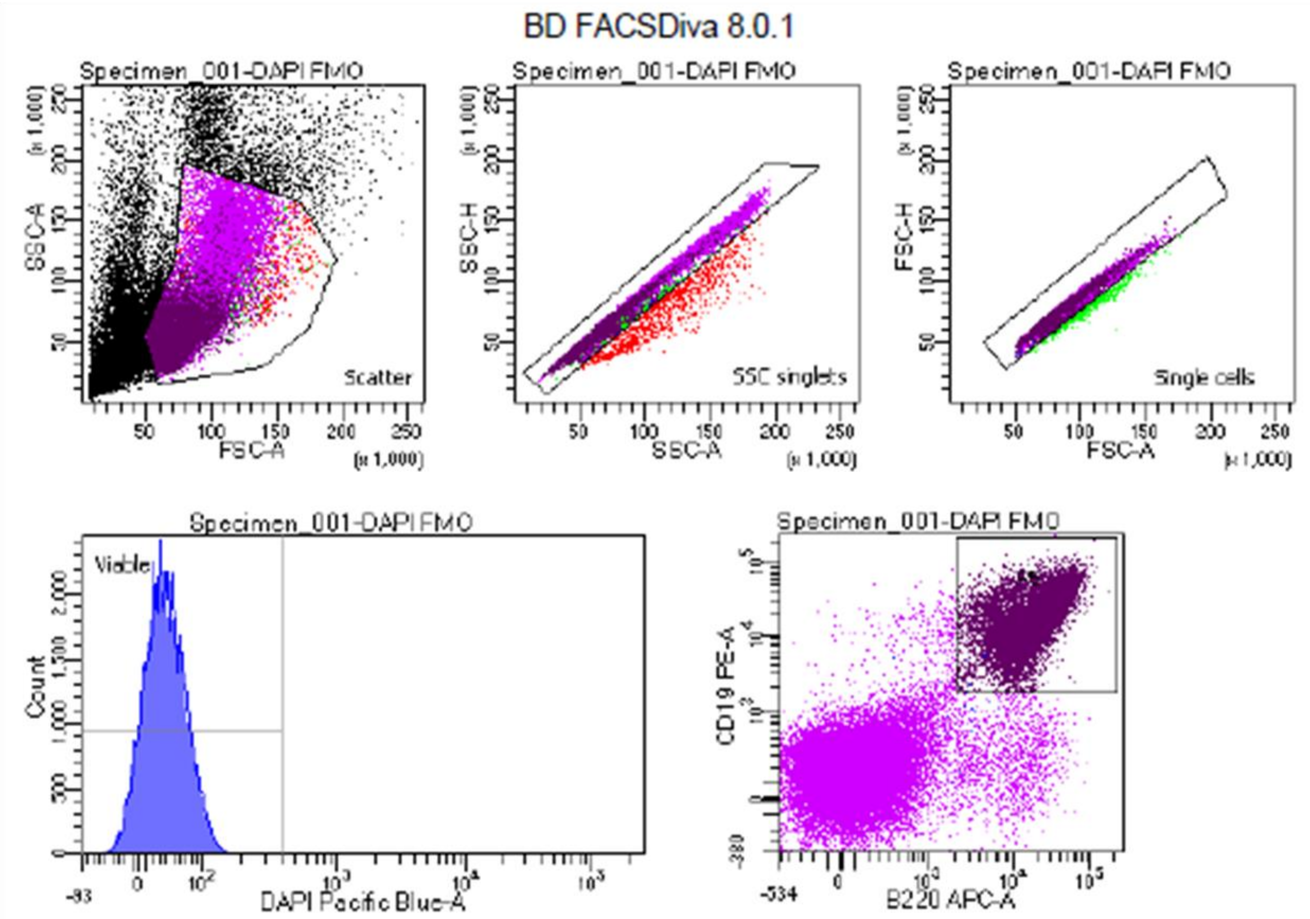


Tube: DAPI FMO

| Population | #Events | %Parent | %Total |
|---|---|---|---|
| All Events | 81,448 | #### | 100.0 |
| Scatter | 52,307 | 64.2 | 64.2 |
| SSC singlets | 51,112 | 97.7 | 62.8 |
| Single cells | 50,376 | 98.6 | 61.9 |
| Viable | 50,302 | 99.9 | 61.8 |
| P5 | 23,910 | 47.5 | 29.4 |

**b**

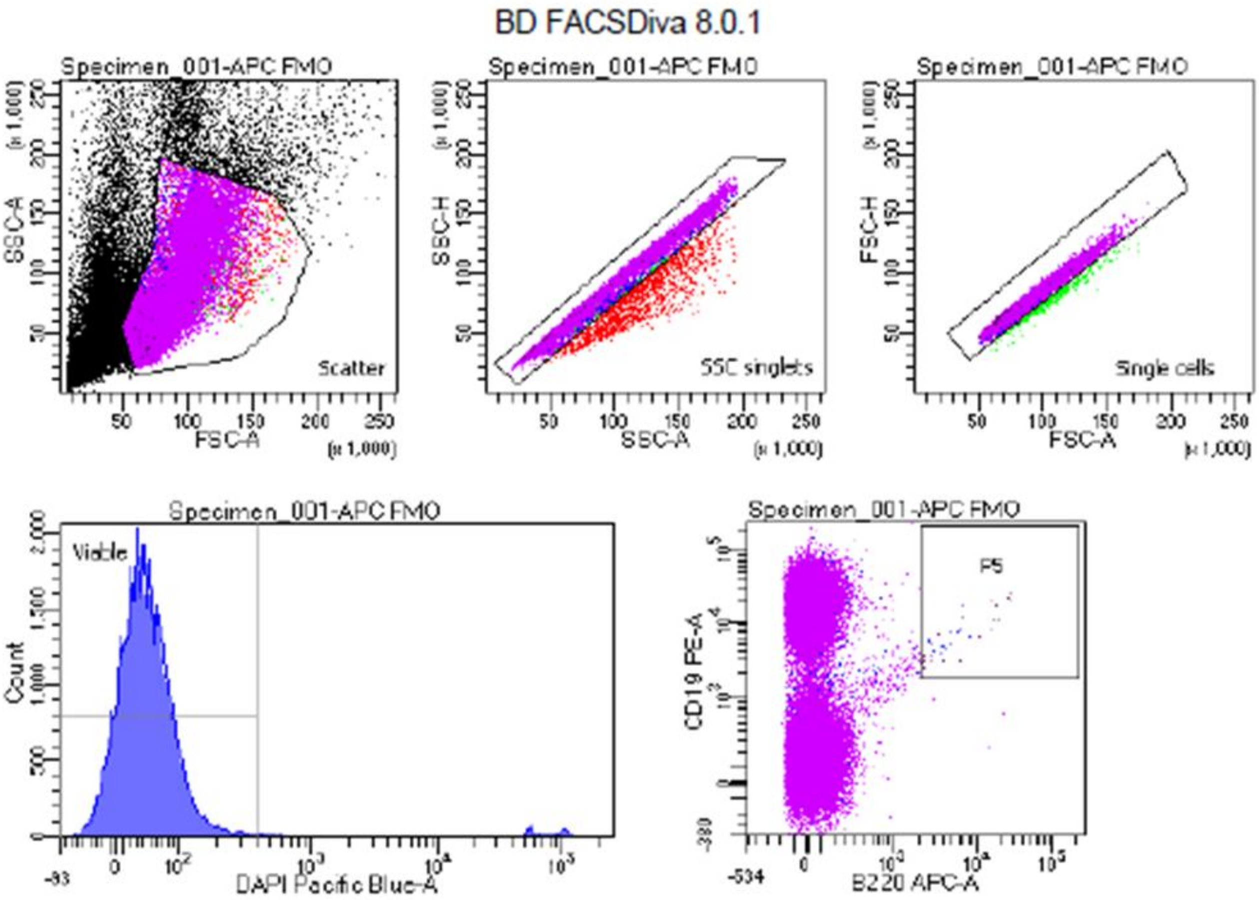


Tube: APC FMO

| Population | #Events | %Parent | %Total |
|---|---|---|---|
| All Events | 91,421 | #### | 100.0 |
| Scatter | 54,222 | 59.3 | 59.3 |
| SSC singlets | 52,697 | 97.2 | 57.6 |
| Single cells | 52,084 | 98.8 | 57.0 |
| Viable | 50,115 | 96.2 | 54.8 |
| P5 | 26 | 0.1 | 0.0 |

c

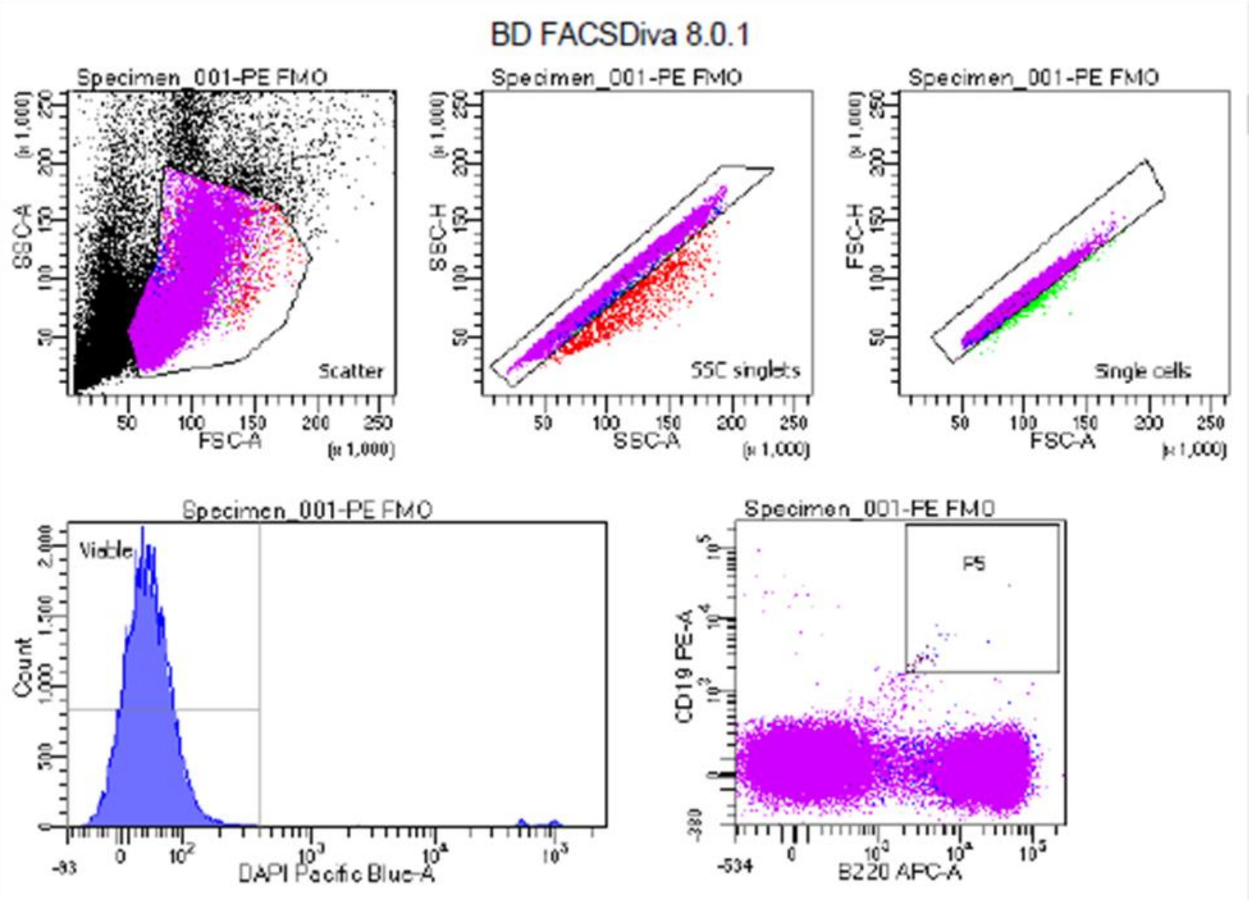


Tube: PE FMO

| Population | #Events | %Parent | %Total |
|---|---|---|---|
| All Events | 87,769 | #### | 100.0 |
| Scatter | 53,736 | 61.2 | 61.2 |
| SSC singlets | 52,424 | 97.6 | 59.7 |
| Single cells | 51,826 | 98.9 | 59.0 |
| Viable | 50,085 | 96.6 | 57.1 |
| P5 | 23 | 0.0 | 0.0 |

**d**

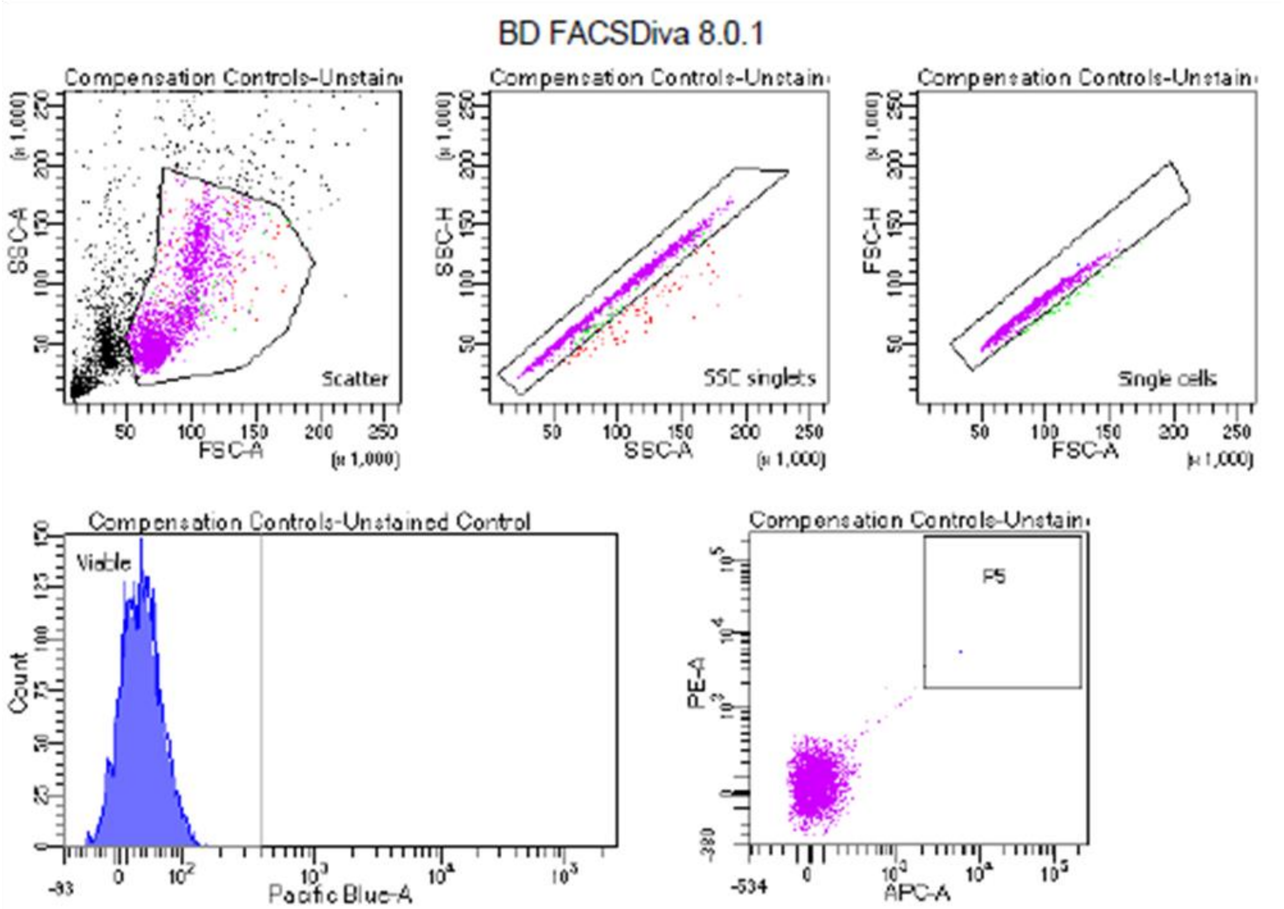


Tube: Unstained Control

| Population | #Events | %Parent | %Total |
|---|---|---|---|
| All Events | 5,000 | #### | 100.0 |
| Scatter | 3,128 | 62.6 | 62.6 |
| SSC singlets | 3,042 | 97.3 | 60.8 |
| Single cells | 2,992 | 98.4 | 59.8 |
| Viable | 2,991 | 100.0 | 59.8 |
| P5 | 1 | 0.0 | 0.0 |

**e**

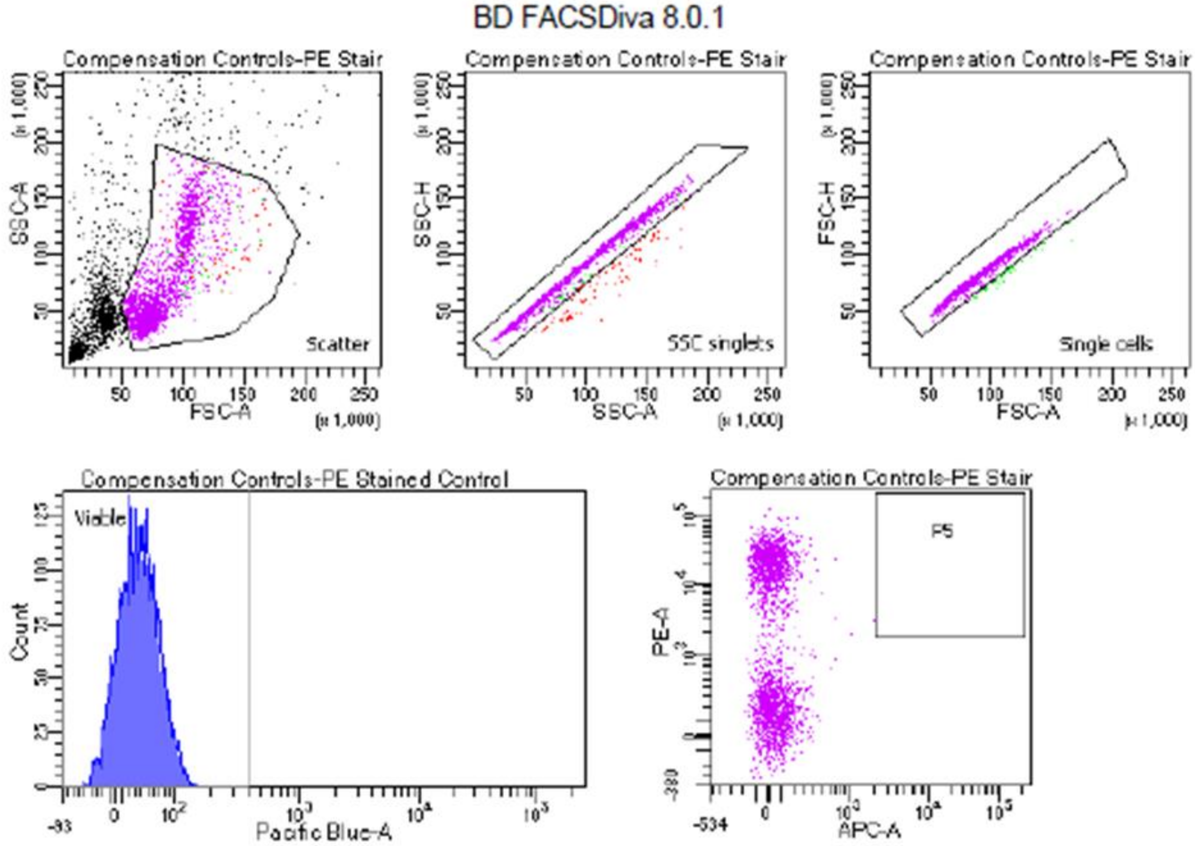


Tube: PE Stained Control

| Population | #Events | %Parent | %Total |
|---|---|---|---|
| All Events | 5,000 | #### | 100.0 |
| Scatter | 3,075 | 61.5 | 61.5 |
| SSC singlets | 2,974 | 96.7 | 59.5 |
| Single cells | 2,925 | 98.4 | 58.5 |
| Viable | 2,925 | 100.0 | 58.5 |
| P5 | 0 | 0.0 | 0.0 |

**f**

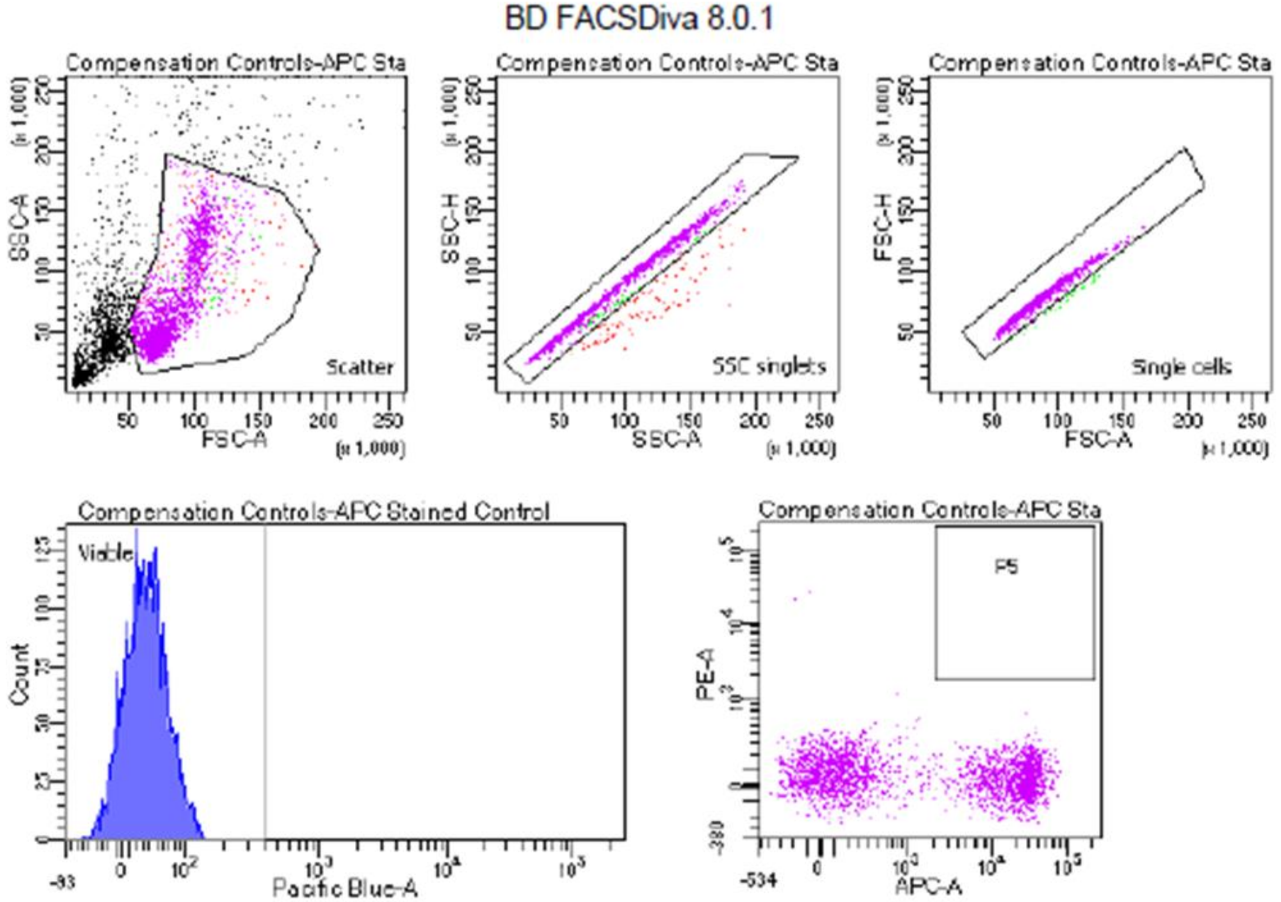


Tube: APC Stained Control

| Population | #Events | %Parent | %Total |
|---|---|---|---|
| All Events | 5,000 | #### | 100.0 |
| Scatter | 3,005 | 60.1 | 60.1 |
| SSC singlets | 2,907 | 96.7 | 58.1 |
| Single cells | 2,862 | 98.5 | 57.2 |
| Viable | 2,862 | 100.0 | 57.2 |
| P5 | 0 | 0.0 | 0.0 |

**g**

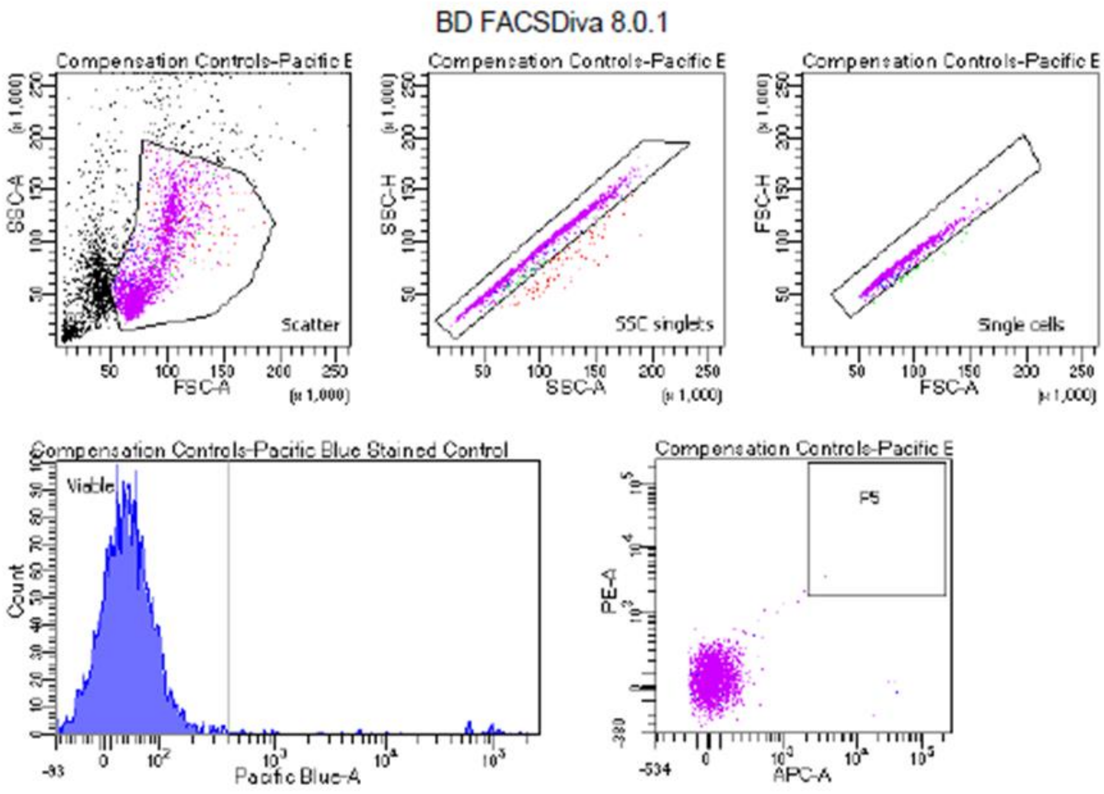


Tube: Pacific Blue Stained Control

| Population | #Events | %Parent | %Total |
|---|---|---|---|
| All Events | 5,000 | #### | 100.0 |
| Scatter | 2,942 | 58.8 | 58.8 |
| SSC singlets | 2,861 | 97.2 | 57.2 |
| Single cells | 2,829 | 98.9 | 56.6 |
| Viable | 2,730 | 96.5 | 54.6 |
| P5 | 1 | 0.0 | 0.0 |

**Supplementary Figure 16: Flow gating for B cells harvested from mouse spleens and livers**

Fluorescence Minus One (FMO) controls using pooled cells from all samples for (A) DAPI (B) APC and (C) PE antibodies to set boundaries for background signal. (D) Unstained pooled cell control (E-G) PE, APC, and DAPI alone stained pooled cell controls used to inform compensation
